# Regulation of PDF receptor signaling controlling daily locomotor rhythms in *Drosophila*

**DOI:** 10.1101/2022.01.03.474875

**Authors:** Weihua Li, Jennifer S. Trigg, Paul H. Taghert

## Abstract

Each day and in conjunction with ambient daylight conditions, neuropeptide PDF regulates the phase and amplitude of locomotor activity rhythms in *Drosophila* through its receptor, PDFR, a Family B G protein-coupled receptor (GPCR). We studied the *in vivo* process by which PDFR signaling turns off, by converting as many as half of the 28 potential sites of phosphorylation in its C terminal tail to a non-phosphorylatable residue (alanine). We report that many such sites are conserved evolutionarily, and their conversion creates a specific behavioral syndrome opposite to loss-of-function phenotypes previously described for *pdfr*. That syndrome includes increases in the amplitudes of both Morning and Evening behavioral peaks, as well as multi-hour delays of the Evening phase. The precise behavioral effects were dependent on day-length, and most effects mapped to conversion of only a few, specific serine residues near the very end of the protein and specific to its A isoform. Behavioral phase delays under entraining conditions were often, but not always correlated with increases in circadian period. The behavioral phenotypes produced by the most severe PDFR variant were ligand-dependent *in vivo*, and not a consequence of changes to their pharmacological properties, nor of changes in their surface expression, as measured *in vitro*. The mechanisms underlying termination of PDFR signaling are complex, subject to regulation that is modified by season, and central to a better understanding of the peptidergic modulation of behavior.

**AUTHOR SUMMARY:** In multi-cellular organisms, circadian pacemakers create output as a series of phase markers across the 24 hour day to allow other cells to pattern diverse aspects of daily rhythmic physiology and behavior. Within circadian pacemaker circuits, neuropeptide signaling is essential to help promote coherent circadian outputs. In the fruit fly *Drosophila* 150 neurons are dedicated circadian clocks and they all tell the same time. In spite of such strong synchronization, they provide diverse phasic outputs in the form of their discrete, asynchronous neuronal activity patterns. Neuropeptide signaling breaks the clock-generated symmetry and drives many pacemakers away from their preferred activity period in the morning. Each day, neuropeptide PDF is released by Morning pacemakers and delays the phase of activity of specific other pacemakers to later parts of the day or night. When and how the PDF that is released in the morning stops acting is unknown. Furthermore, timing of signal termination is not fixed because day length changes each day, hence the modulatory delay exerted by PDF must itself be regulated. Here we test a canonical model of G protein-coupled receptor physiology to ask how PDF receptor signaling is normally de-activated. We use behavioral measures to define sequence elements of the receptor whose post-translational modifications (e.g., phosphorylation) may define the duration of receptor signaling.

## INTRODUCTION

In *Drosophila*, neuropeptide PDF signaling helps pattern the output of the fly circadian pacemaker network that controls rhythmic daily locomotor activity [1–3]. Its functions have been compared to those of neuropeptide VIP in regulating mammalian circadian physiology [4]. Historically PDF was first isolated as an active principle (Pigment Dispersing Hormone) that mediates light-dependent dispersion of pigment granules in diverse chromatophores of crustacea [5, 6]. In the insect circadian system, PDF acts for a specified period within each 24 hr cycle, and works in conjunction with environmental light to set phase and amplitude for locomotor activity rhythms that normally occur around dawn and dusk [7–9]. Each day, the precise times of dusk and of dawn change, which alters the time interval between them. These facts require that the time of PDF signaling, the point when it starts and the point when it stops, must also be adjusted each day to appropriately follow and reflect these daily variations in the light:dark transitions. PDF signaling starts following its release by specific pacemaker neurons, whose period of activity *in vivo* tracks the dawn in a variety of photoperiodic conditions [10]. We lack a comparable understanding of how PDF receptor signaling normally stops: this work addresses that mechanism.

The PDF receptor (PDFR) is a member of the Family B (secretin receptor-like) GPCR group [11–13]: it is G_s_-coupled and its activation elevates cAMP levels *in vivo* [14]. It regulates different adenylate cyclases (AC) in diverse target pacemakers (15, 16), which, through PKA activation, ultimately regulate the pace of the molecular clock through regulation of Timeless [17]. PDFR autoreceptor signaling promotes dramatic, daily morphological changes in the axonal terminals of sLNv pacemakers [18]. In addition to its effects on the pace of the molecular pacemaker, PDFR activation also regulates calcium dynamics in subsets of pacemaker neuron groups to help dictate their group-specific, daily phases of activation (in the sLNv, in the 5^th^ sLNv and in subsets of LNd, and DN3 groups – [10]). Such target cell-specific delays of PER-dependent neuronal activity illustrate the basis by which the circadian network produces a daily series of staggered phasic, neuronal outputs [19, 20]. Finally, PDF/PDFR signaling is long-lasting: its depression of basal calcium levels in target neurons persists without abatement over many hours [19]. These observations raise fundamental questions regarding the mechanism and the time course by which PDFR signaling diminishes in anticipation of the next day’s cycle of signaling.

The canonical model of GPCR phosphorylation and homologous desensitization features G protein-coupled receptor kinases (GRKs) which associate with activated GPCRs and phosphorylate cytosolic segments, thereby recruiting β-arrestins [21, 22]. β-arrestins uncouple the receptors from G proteins [23, 24]. or enhance receptor endocytosis [25]; they can also serve as signal transducers by recruiting distinct signaling molecules [26]. A second major regulatory mechanism to reduce GPCR signaling is heterologous desensitization, whereby second-messenger-dependent kinases (PKA or PKC) phosphorylate GPCRs [27]. Thus, we designed experiments to modify evolutionarily-conserved residues in the C terminal tail of PDFR that could conceivably serve as substrates for phosphorylation and subsequent signal termination. Our working hypothesis was that, by their actions *in vivo*, such modified PDF receptors would reveal extended lifetimes of activation.

We know very little about the mechanisms that underlie normal termination of PDFR signaling. Levels of *pdfr* cycle in many pacemaker groups in the *Drosophila* brain, but they peak at different times, in different groups [28]. Sensitivity to PDF *in vivo* peaks in the early day and is regulated by the PER-dependent clock, via post-transcriptional mechanisms: the EC50 for PDF responses in identified neurons varies systematically 5-10 fold: as a consequence of RalA action, as a function of time of day, and as a function of seasonality [29]. PDFR signaling is long-lasting: it persists for many hours *in vivo* [19], for as long as free peptide ligand is available in the bath [14]. In addition, β-arrestin2-GFP is not efficiently recruited to activated PDFR when the receptor is functionally-expressed in *hEK-293T* cells [29]. In contrast, each of 13 other *Drosophila* neuropeptide GPCRs (including two Family B GPCRs, CG8422 and CG17415) efficiently recruit β-arrestin2-GFP, when they are activated by their cognate ligands in that cellular environment [30–32].

Here we report (i) that PDFR is normally phosphorylated *in vivo* at conserved C terminal residues; (ii) that loss of conserved PDFR phosphorylatable sites leads to a behavioral syndrome opposite to loss-of-function *pdf* and *pdfr* phenotypes, with effects on both the amplitude and phase of the daily locomotor peaks. This ‘gain of function’ approach reveals a multi-hour range of potential phases for both the Morning and Evening activity peaks, within which neuropeptide PDF:PDFR signaling normally specifies rhythmic behavior, according to season. In addition, using a structure-function approach, this work identifies specific PDFR sequence elements that are major points at which the duration of receptor activity is regulated, and through which behavior is modulated in season-specific fashion.

## RESULTS

### PDFR C-Terminal sequences

Based on alternative splicing, the *pdfr* locus in *Drosophila melanogaster* (CG13758) encodes very similar GPCRs which differ in their extreme C terminal sequences, for which the PA and PD isoforms are representative (flybase.org/reports/FBgn0260753). The PD isoform is slightly longer and lacks the final ∼20 AAs of the PA isoform. As described in the Supplemental Information (*PDFR Isoforms*), we focused on the PA protein isoform of PDFR, as it has been used in the majority of genetic studies in the field. To identify residues for mutation, we first used comparative genomic analyses to assess how well specific sequences in the C-terminal region of the PA receptor are conserved. We obtained annotated *pdfr* genomic sequences from 16 additional species of *Drosophila*, in both the *Sophophora* and *Drosophila* sub-families (**Suppl. Table 1**). Together this species collection represents an estimated 40-60 MYr of *Drosophalid* evolution [33]. We defined residue V505 of the *melanogaster* protein as the start of the C terminal sequence, following the consensus 7^th^ transmembrane TM domain (TM7) (**Suppl. Figure 1**). These 17 different PDFRs all contain C-termini of considerable length, and vary between 189 to 215 amino acids (AAs). The *melanogaster* PDFR-A C-terminal contains 28 Ser, Thr, or Tyr residues (these may be subject to post-translational phosphorylation and de-phosphorylation, and/ or other modifications). To survey putative functionality among these, we chose 14 residues that are distributed across the length of the C-terminal and which display high evolutionary conservation (**Suppl. Figure 1** and **Suppl. Table 3)**. For naming purposes, we grouped them into arbitrary clusters (CL) numbered #1 to #7, with positions shown in Figure 1. Following a common paradigm in study of GPCR physiology [e.g., 34-36], we performed an “Alanine Scan”: testing the consequences of their mutation to Alanine, which is a non-phosphorylatable analog [41]. Some clusters have only a single modified AA (e.g. CL4 and CL5), while in others we concurrently modified two AAs (e.g., CL1, CL6 and CL7), and in others, as many as six closely-positioned residues (e.g., CL2-3).

**Figure 1.**
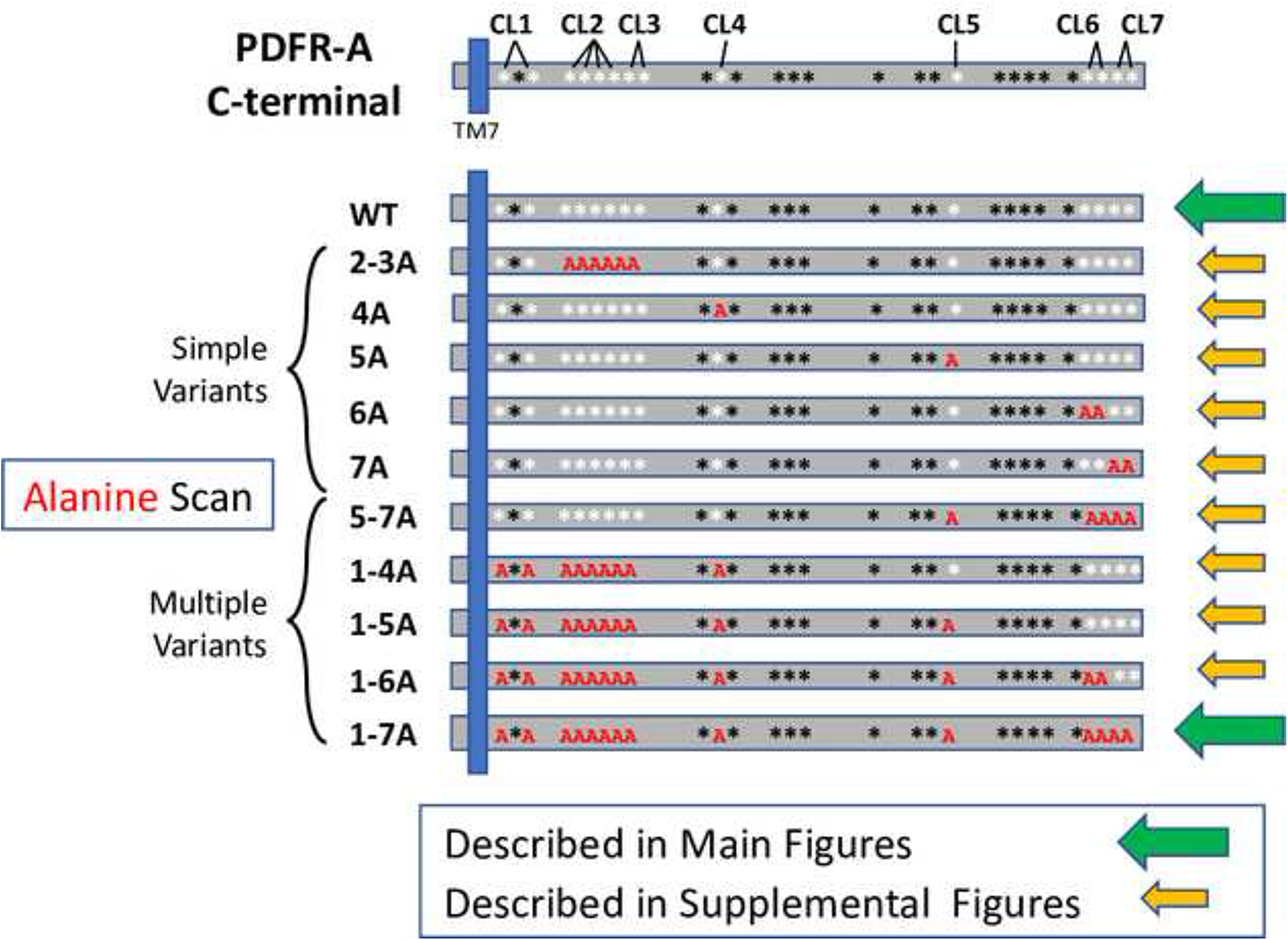
A map illustrating the series of PDFR sequence variants used to test a role for PDFR phosphorylation in regulating the locomotor rhythms. The C Terminal of the PDFR-A isoform is diagrammed (top) to the right of the 7^th^ transmembrane domain (TM7, blue). Asterisks indicate the positions of the 28 serine, threonine or tyrosine residues present within the PDFR C Terminal tail. The residues marked by the white asterisks were those chosen for study by sequence alteration to encode Alanine; they were given arbitrary designations as Clusters (CL) 1 through 7. Each cluster contains a single residue (e.g., CL4 or CL5), or as many as four residues (e.g., CL2). The ten sequence variants are named by the Cluster(s) that was altered followed by the letter ‘A”. Simple Variants include those wherein residues in one or two Clusters were altered. Multiple Variants include those wherein residues within three or more Clusters were altered. All 14 targeted residues were altered in the PDFR 1-7A variant. The green arrows mark the WT and 1-7A PDFR variant: their behavioral effects are featured in the Main Figures. The yellow arrows mark the nine other PDFR variants: their behavioral effects are described in Supplemental Figures.

### Study Overview

This series of variant receptors contains 10 different mutated versions of PDFR (not including the WT ‘parent’ PDFR-A) and is arbitrarily divided into two broad categories as indicated in Figure 1. The first group (termed “<u>Simple Variants</u>”) targeted one or two clusters of conserved Ser/Thr/Tyr AAs (i.e., CL 2-3A; 4A; 5A; 6A; 7A). The second category (termed “<u>Multiple Variants</u>”) targeted three or more of the AA clusters in various combinations (i.e., CL 1-4A; 1-5A; 1-6A; 1-7A and 5-7A). The CL1-7A variant is the most severe as it mutates all 14 of the targeted AA residues. We studied these ten PDFR variants, as well as the WT receptor, following expression in a *pdfr* mutant strain (*han^5537^* [11]; we measured the ability of individual PDFR variants to rescue and to shape the phases and periodicity of rhythmic locomotor behavior. For the most part, we analyzed PDFR variants in the *pdfr* mutant background to permit evaluation of their properties, without competition from endogenous PDFR. Normally, PDFR-A is expressed throughout the ∼150 cell pacemaker network, typically in subsets of each clustered group, as revealed by its expression from an epitope-marked BAC transgene [37]. However, no simple *Pdfr*-Gal4 driver element recapitulates a majority of normal expression sites [37]. Therefore to effect broad PDFR-A expression in the pacemaker system, we used a *timeless*-Gal4 driver element. Because PDF sensitivity varies with seasons [37], we tested this PDFR series in different photoperiodic conditions, as well as in constant darkness. *pdfr* loss-of-function mutants display advanced behavioral peaks under Light:Dark conditions, as well as weak and shortened free-running periods under constant darkness [11].

The behavioral actions of the Ala-mutated PDFRs either resembled those of WT receptor, or exhibited gain-of-function properties (behavioral actions opposite to those seen in loss-of-function *pdfr* mutant flies). For purpose of clarity, the Main Figures report a comparison of properties for the WT PDFR with those of the most extensively-mutated version of PDFR (called 1-7A), as indicated in Fig. 1. Comparable data on the properties of the other variant receptors in the series is described in Supplemental Figures. To better interpret the results produced by the different sequence variants, we also present experiments that test assumptions used in the experimental design. These tested the following hypotheses: i) that the behavioral effects of receptor variants are independent of activation by the endogenous ligand, neuropeptide PDF; and ii) that the bulky C-terminal fusion of GFP (present in all variants tested) strongly influenced the results. Both hypotheses were dis-proved.

### Effects of WT vs 1-7A PDFR on locomotor behavior in Short Day (winter-like) conditions (Figure 2)

In 8L:16D, *pdfr^han^* mutant flies lack a prominent morning peak and their evening peak of activity begins 1-2 hr earlier than controls; the example shown in Fig. 2A-1 (*tim*>*no transgene,* boxed in yellow) is heterozygous for the *tim*-Gal4 element. Rescue by a WT-*pdfr* cDNA (Fig. 2B-1 (*tim*>*pdfr*, boxed in black)) did not produce obvious effects on activity in the time domain preceding or just past Lights-On (Morning). However, it strongly affected the Evening peak: not its amplitude (Fig. 2B-3 **and** B-4) but significantly delaying its phase by about 1 h (Fig. 2E) such that the Evening peak now extends past after Lights-off (Fig. 2B-4). In contrast, expression of the 1-7A Multiple Variant driven by *tim*-Gal4 (*tim> pdfr1-7A*, boxed in red) elevated the amplitude of activity during the period of ZT 18-21 prior to lights-ON (Fig. 2C-1 **and** C-2**, orange arrow**): we speculate this is promotion of a “Morning” peak of activity. It has the same phase as that produced by expression of the WT receptor (Fig. 2D), but significantly larger amplitude (Fig. 2C-2**, orange arrow**). The 1-7A variant also delayed the Evening peak but to a much greater extent than did the WT PDFR (Fig. 2C-1, C-4, blue arrow), producing a conspicuous, large amplitude peak that occurred on average as late as ∼3 h after lights-off, at ZT12, significantly more delayed than the peak produced by WT PDFR (Fig. 2E). Following short day light entrainment, flies were released into constant darkness for ∼7-8 days (DD - constant conditions): resulting locomotor activity is displayed in the double-plotted group actograms (Fig. 2F **-** yellow box**, G -** black box, and **H -** red box). The properties of the persistent circadian rhythmic behavior are provided in Table 1. The period (tau) of the 1-7A variant was considerably lengthened. The phase of the major DD rhythm matched that of the much-delayed evening peak in the 1-7A flies (Fig. 2H, green line). In sum, under winter-like photoperiods, the PDFR 1-7A variant significantly increased the amplitude of the morning activity peak and significantly delayed the phase of the evening peak; the latter phase was also reflected in persistent rhythmic activity under subsequent DD.

**Figure 2.**
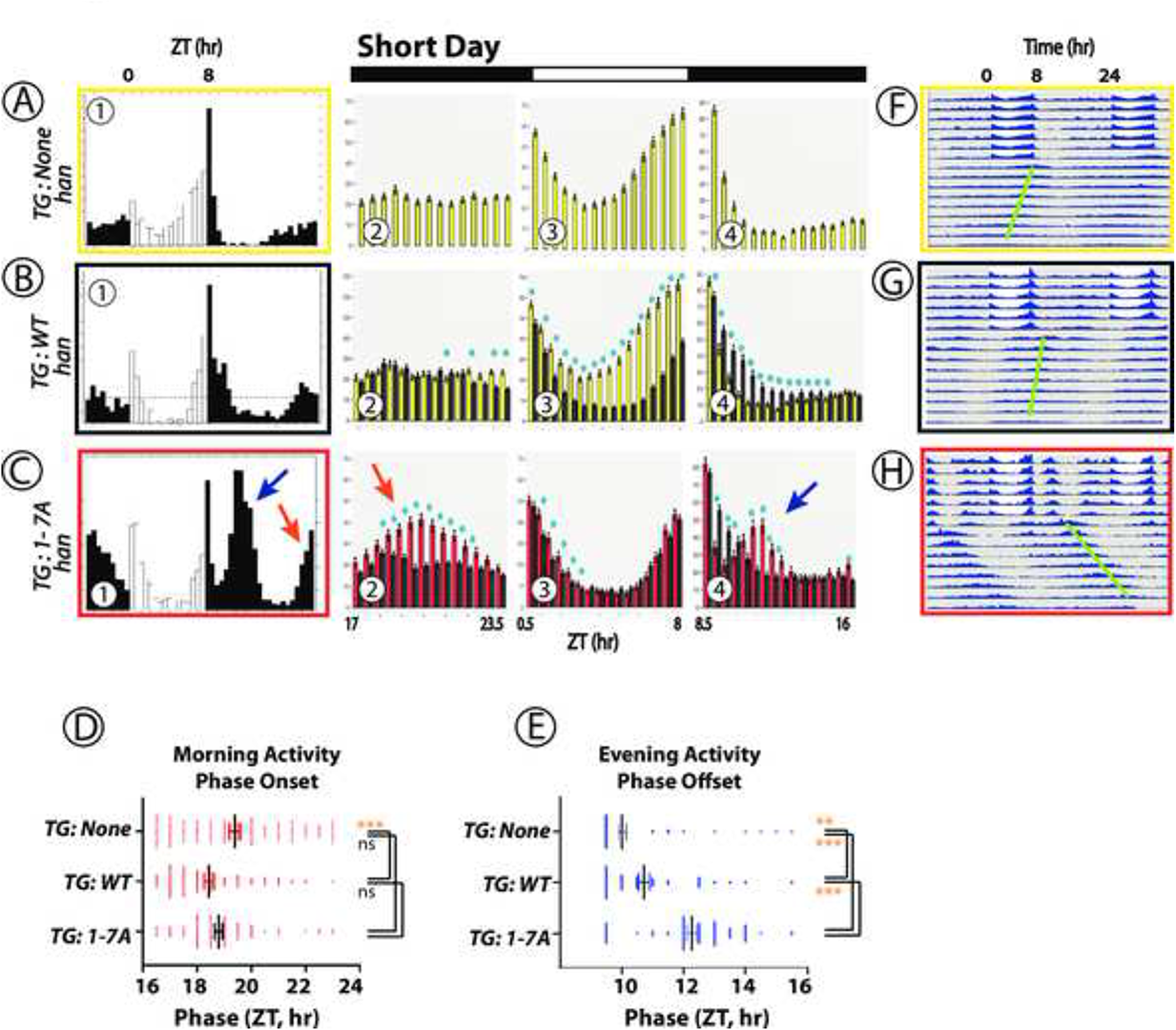
Locomotor Rhythms exhibited by *pdfr* mutant flies expressing the WT PDFR versus the 1-7A PDFR Variant under Winter-like (Short Day) conditions. All behavioral records were recorded from *han* (*pdfr* mutant) flies that expressed either no UAS transgene (A, yellow box), or a UAS-WT *pdfr* transgene (B, black box) or a UAS-1-7A *pdfr* (C, red box)). Panels (A)-(C) contains sub-panels 1-4. Sub-Panel (1) displays group eductions (activity averaged over days 5 and 6 of light entrainment) with open bars indicating the 8-hr periods of Lights-on and filled bars the 16-hr periods of Lights-off. *TG*: transgene. The red and blue arrows indicate Morning and Evening activity periods respectively in the 1-7A records that have distinguished amplitude or phase. Amplitude measures are displayed in Panels 2-4: 30 min bins for each genotype, color-coded and directly-compared (i.e., Panel C-2 through C-4 compares the amplitudes of the WT PDFR activity (black) with that of the 1-7A PDFR activity (red). The bins representing the light-dark transitions were removed. Blue asterisks mark amplitudes that are significantly different between genotypes by ANOVA followed by a Student’s T-test (p < 0.05). Panel (D) displays the average Phase Onset timepoint for the Morning activity for each genotype over the last two days of entrainment (LD 5-6). Panel (E) displays the average Phase Offset timepoint for the Evening activity for each genotype over the last two days of entrainment (LD 5-6). Analyses in (D) and (E) represent ANOVA followed by Dunnett’s post hoc multiple comparisons of all compared to WT: ns = not significant; * = p<0.05; ** = p<0.01; *** = p<0.005; **** = p<0.001. Panels (F) – (H) present double plotted actograms of group activity for each genotype (color-coded) over the ∼6 days of light entrainment, followed by ∼9 days of constant darkness (DD). The green bars indicate the phases of the dominant activity periods in DD.

**Figure 3.**
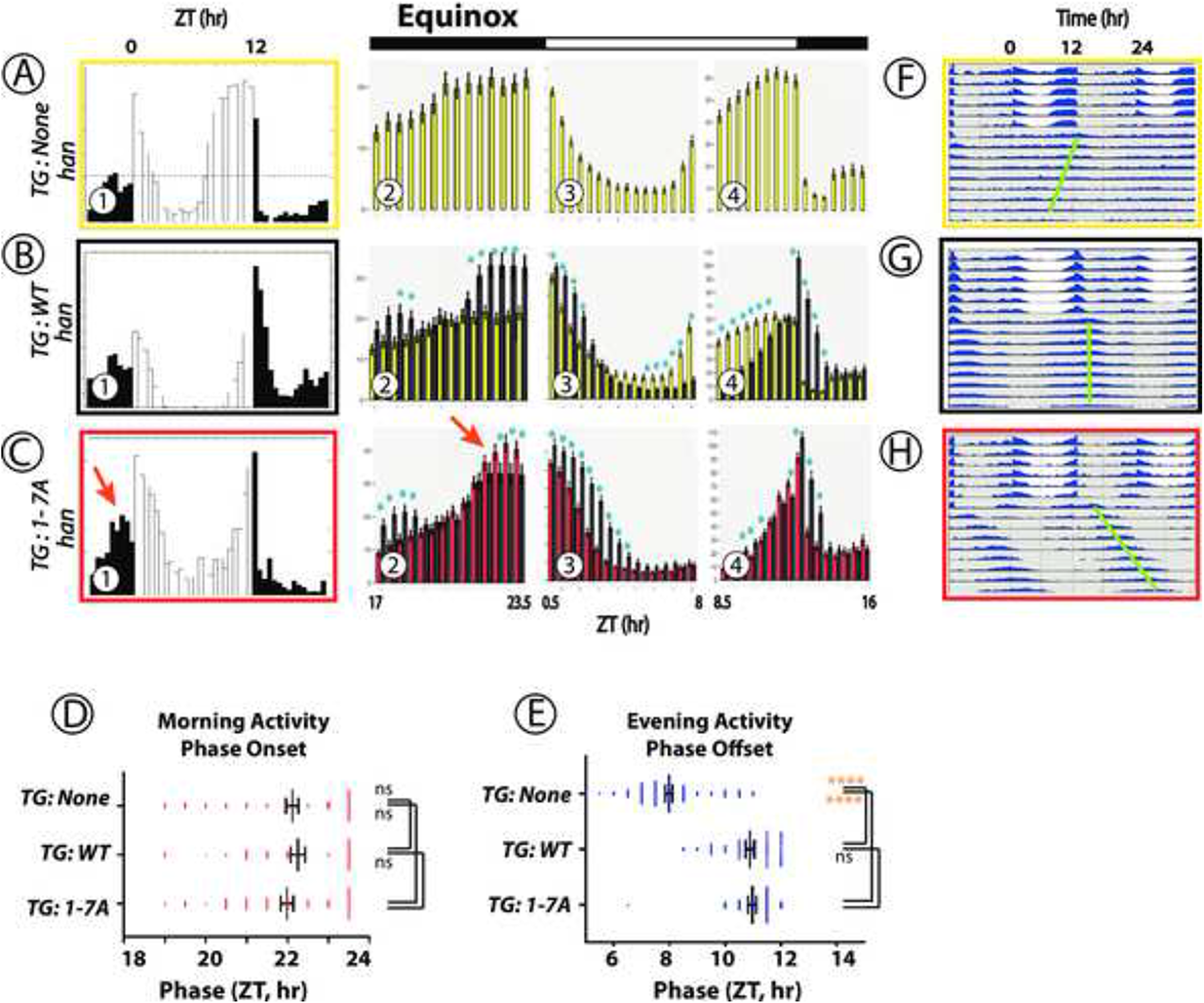
Locomotor Rhythms exhibited by *pdfr* mutant flies expressing the WT PDFR versus the 1-7A PDFR Variant under Equinox-like (12L:12D) conditions. All behavioral records were recorded from *han* (*pdfr* mutant) flies that expressed either no UAS transgene (A, yellow box), or a UAS-WT *pdfr* transgene (B, black box) or a UAS-1-7A *pdfr* (C, red box)). The Panels display behavioral activity with a format similar to the one shown in Figure 2. Panels (A)-(C) display average group eductions (activity averaged over days 5 and 6 of light entrainment, with open bars indicating the 12-hr periods of Lights-on and filled bars the 12-hr periods of Lights-off) and bin-by-bin analyses to compare activity amplitudes between genotypes. *TG*: transgene. The red arrow indicates Morning activity in flies expressing the 1-7A variant with distinguished amplitude. Blue asterisks mark amplitudes that are significantly different between genotypes by ANOVA followed by a Student’s T-test (p < 0.05). Panels (D) and (E) display the phases of morning Onsets and Evening Offsets respectively. Gold asterisks: ANOVA followed by Dunnett’s post hoc multiple comparisons of all compared to WT: ns = not significant; * = p<0.05; ** = p<0.01; *** = p<0.005; **** = p<0.001. Panels (F) – (H) present double plotted actograms of group activity for each genotype over the ∼6 days of light entrainment, followed by ∼9 days of constant darkness (DD). The green bars indicate the phase of the dominant activity period in DD.

**Figure 4.**
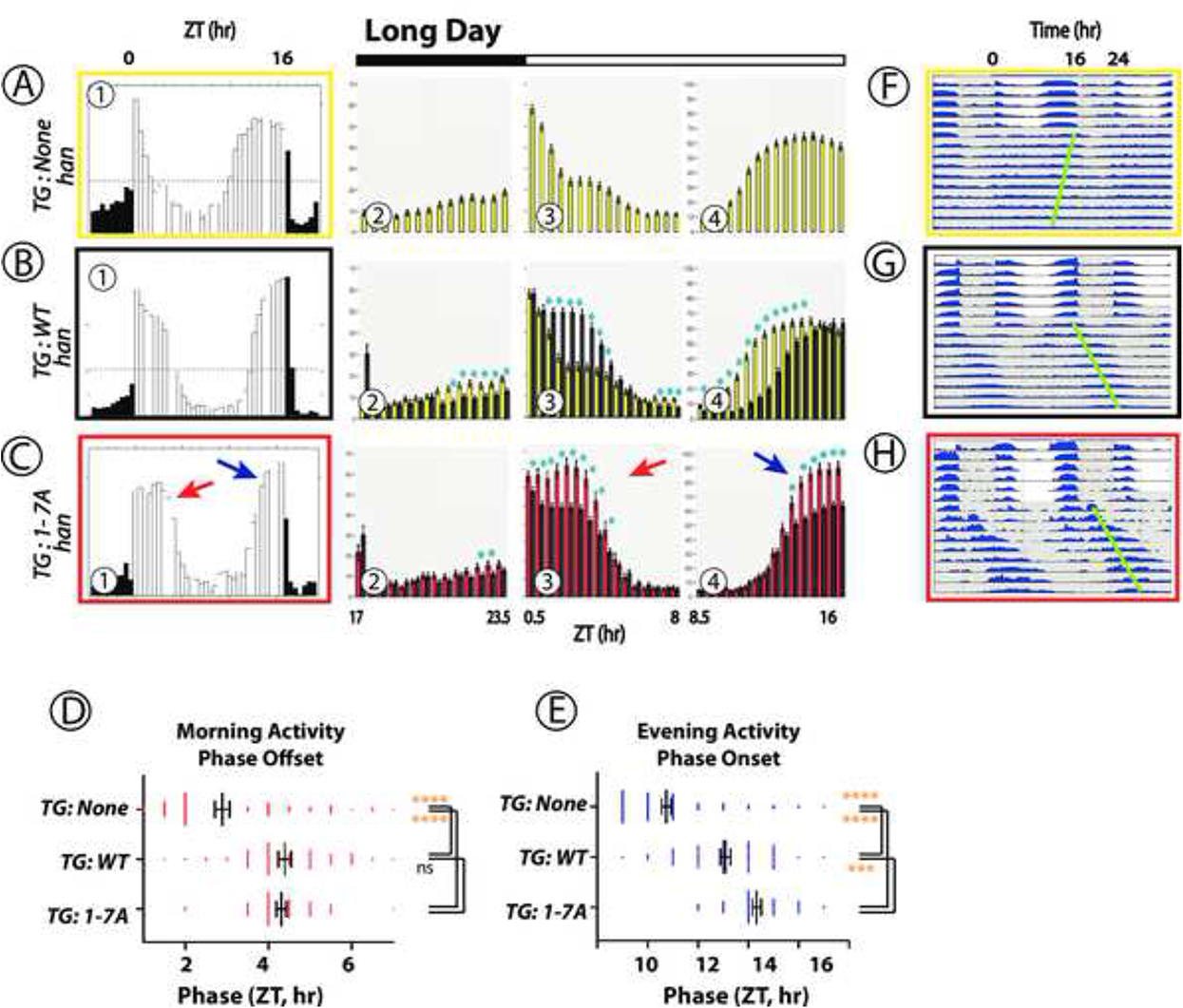
Locomotor Rhythms exhibited by *pdfr* mutant flies expressing the WT PDFR versus the 1-7A PDFR Variant under Long-day (summer-like) conditions. All behavioral records were recorded from *han* (*pdfr* mutant) flies that expressed either no UAS transgene (A, yellow box), or a UAS-WT *pdfr* transgene (B, black box) or a UAS-1-7A *pdfr* (C, red box)). The Panels present behavioral activity with a format similar to the one shown in Figure 2. Panels (A)-(C) display average group eductions (activity averaged over days 5 and 6 of light entrainment, with open bars indicating the 16-hr periods of Lights-on and filled bars the 8-hr periods of Lights-off) and bin-by-bin analyses to compare activity amplitudes between genotypes. *TG*: transgene. The red and blue arrows indicate Morning and Evening activity respectively, in flies expressing the 1-7A variant with distinguished amplitude and/or phase. Blue asterisks mark amplitudes that are significantly different between genotypes by ANOVA followed by a Student’s T-test (p < 0.05). Panels (D) and (E) display the phases of Morning Offsets and Evening Onsets respectively. Gold asterisks: ANOVA followed by Dunnett’s post hoc multiple comparisons of all compared to WT: ns = not significant; * = p<0.05; ** = p<0.01; *** = p<0.005; **** = p<0.001. Panels (F) – (H) display double plotted actograms of activity during the ∼6 days of light:dark conditions, followed by activity during ∼9 days of DD. The green bars indicate the phase of the dominant activity period in DD.

**Table 1.** Locomotor activity measures of various genotypes during constant dark conditions.

| Short Day Entrainment | N | n | %AR | tau | SEM | Average tau different from: |  |  |
| --- | --- | --- | --- | --- | --- | --- | --- | --- |
|  |  |  |  |  |  | w[1118]* | pdfrEgfp** | pdf[01] |
| *han[5304]; tim > w[1118]* | 5 | 72 | 22% | 23.9 | 0.09 | - | ns | nd |
| *han[5304]; tim > pdfr* | 1 | 16 | 6% | 23.7 | 0.13 | ns | ns | nd |
| *han[5304]; tim > pdfr1-7A No GFP* | 3 | 30 | 8% | 23.6 | 0.07 | ns | ns | nd |
| *han[5304]; tim > pdfrEgfp* | 3 | 46 | 35% | 24.0 | 0.09 | ns | - | nd |
| *han[5304]; tim > pdfrEgfp23A* | 2 | 31 | 13% | 24.8 | 0.12 | ns | ns | nd |
| *han[5304]; tim > pdfrEgfp4A* | 2 | 31 | 26% | 24.7 | 0.13 | ns | ns | nd |
| *han[5304]; tim > pdfrEgfp5A* | 2 | 43 | 26% | 24.6 | 0.11 | ns | ns | nd |
| *han[5304]; tim > pdfrEgfp6A* | 3 | 50 | 38% | 24.4 | 0.08 | ns | ns | nd |
| *han[5304]; tim > pdfrEgfp7A* | 3 | 45 | 27% | 24.8 | 0.11 | ns | ns | nd |
| *han[5304]; tim > pdfrEgfp1-4A* | 3 | 40 | 12% | 24.6 | 0.12 | ns | ns | nd |
| *han[5304]; tim > pdfrEgfp1-5A* | 3 | 31 | 16% | 25.0 | 0.23 | *** | * | nd |
| *han[5304]; tim > pdfrEgfp1-6A* | 3 | 42 | 14% | 25.1 | 0.13 | **** | ** | nd |
| *han[5304]; tim > pdfrEgfp1-7A* | 4 | 59 | 16% | 24.9 | 0.11 | **** | * | nd |
| *han[5304]; tim > pdfrEgfp567A* | 2 | 37 | 16% | 23.9 | 0.09 | ns | ns | nd |
| *tim > pdfrEgfp1-7A* | 1 | 15 | 0% | 25.8 | 0.25 | nd | nd | **** |
| *pdf[01]* | 1 | 12 | 67% | 22.5 | 0.12 | nd | nd | - |
| *tim > pdfrEgfp1-7A; pdf[01]/pdf[01]* | 1 | 25 | 20% | 23.6 | 0.12 | nd | nd | ns |

| Equinox Entrainment | N | n | %AR | tau | SEM |  |  |
| --- | --- | --- | --- | --- | --- | --- | --- |
| *han[5304]; tim > w[1118]* | 6 | 68 | 34% | 23.7 | 0.09 | - | ns |
| *han[5304]; tim > pdfr* | 3 | 16 | 25% | 23.5 | 0.16 | ns | ns |
| *han[5304]; tim > pdfr1-7A No GFP* | 3 | 35 | 14% | 23.7 | 0.08 | ns | ns |
| *han[5304]; tim > pdfrEgfp* | 4 | 48 | 13% | 24.1 | 0.06 | ns | - |
| *han[5304]; tim > pdfrEgfp23A* | 2 | 54 | 48% | 25.6 | 0.12 | **** | **** |
| *han[5304]; tim > pdfrEgfp4A* | 2 | 44 | 32% | 24.1 | 0.07 | ns | ns |
| *han[5304]; tim > pdfrEgfp5A* | 2 | 47 | 34% | 24.4 | 0.06 | ns | ns |
| *han[5304]; tim > pdfrEgfp6A* | 3 | 35 | 23% | 24.3 | 0.13 | ns | ns |
| *han[5304]; tim > pdfrEgfp7A* | 3 | 47 | 21% | 24.1 | 0.09 | ns | ns |
| *han[5304]; tim > pdfrEgfp1-4A* | 3 | 45 | 18% | 24.5 | 0.08 | ** | ns |
| *han[5304]; tim > pdfrEgfp1-5A* | 2 | 32 | 6% | 25.0 | 0.13 | **** | *** |
| *han[5304]; tim > pdfrEgfp1-6A* | 2 | 32 | 19% | 25.3 | 0.14 | **** | **** |
| *han[5304]; tim > pdfrEgfp1-7A* | 6 | 88 | 6% | 25.1 | 0.17 | **** | **** |
| *han[5304]; tim > pdfrEgfp567A* | 2 | 45 | 24% | 24.1 | 0.08 | ns | ns |

| Long Day Entrainment | N | n | %AR | tau | SEM |  |  |  |
| --- | --- | --- | --- | --- | --- | --- | --- | --- |
| *han[5304]; tim > w[1118]* | 7 | 105 | 47% | 23.8 | 0.06 | - | ns | nd |
| *han[5304]; tim > pdfr* | 3 | 47 | 17% | 24.9 | 0.15 | ns | ns | nd |
| *han[5304]; tim > pdfr1-7A No GFP* | 3 | 40 | 48% | 24.6 | 0.09 | ns | ns | nd |
| *han[5304]; tim > pdfrEgfp* | 4 | 82 | 17% | 24.9 | 0.08 | ns | - | nd |
| *han[5304]; tim > pdfrEgfp23A* | 2 | 48 | 23% | 25.4 | 0.12 | **** | **** | nd |
| *han[5304]; tim > pdfrEgfp4A* | 1 | 27 | 22% | 25.5 | 0.16 | **** | **** | nd |
| *han[5304]; tim > pdfrEgfp5A* | 1 | 36 | 11% | 24.7 | 0.16 | ns | ns | nd |
| *han[5304]; tim > pdfrEgfp6A* | 2 | 51 | 22% | 25.0 | 0.09 | ns | ns | nd |
| *han[5304]; tim > pdfrEgfp7A* | 2 | 54 | 13% | 25.5 | 0.09 | **** | **** | nd |
| *han[5304]; tim > pdfrEgfp1-4A* | 3 | 40 | 8% | 25.2 | 0.13 | **** | ns | nd |
| *han[5304]; tim > pdfrEgfp1-5A* | 3 | 33 | 6% | 25.2 | 0.14 | **** | * | nd |
| *han[5304]; tim > pdfrEgfp1-6A* | 3 | 48 | 2% | 25.6 | 0.09 | **** | **** | nd |
| *han[5304]; tim > pdfrEgfp1-7A* | 4 | 63 | 14% | 25.3 | 0.08 | **** | *** | nd |
| *han[5304]; tim > pdfrEgfp567A* | 1 | 38 | 18% | 24.3 | 0.09 | ns | ns | nd |
| *tim > pdfrEgfp1-7A* | 1 | 14 | 14% | 25.5 | 0.13 | nd | nd | **** |
| *pdf[01]* | 1 | 32 | 88% | 22.7 | 0.05 | nd | nd | - |
| *tim > pdfrEgfp1-7A; pdf[01]/pdf[01]* | 1 | 29 | 21% | 23.3 | 0.09 | nd | nd | ns |

### Behavioral effects of the other PDFR Variants under Short Day (winter-like) conditions (Suppl. Fig. 2 & 3)

We tested each of the nine other PDFR Ala-variants by comparing their effects on locomotor behavior as we had done for the PDFR 1-7A variant. **Suppl. Figure 2** presents these collected data with Group eductions, Actograms and Phase analysis; **Suppl. Fig. 3** presents these same collected data with bin-by-bin comparisons to WT-PDFR to consider amplitude changes. The phase of the Morning peak was not obviously or systematically effected (**Suppl. Fig. 2L**), while that of the Evening peak was significantly delayed by several (**Suppl. Fig. 2M**). We note two features – first the progressive addition of more Ala substitutions in the series 1-4A, 1-5A, 1-6A (**Suppl. Fig. 2I-K**) tended to produce large and delayed Evening peaks in the time period ZT11.5-12.5 (3-4 h after lights-off), with the variant 1-6A producing the most pronounced delay. The delayed Evening peak activity at ZT12 sometimes appeared at the expense of, the normal Evening peak activity that occurred prior to Lights-off (e.g., **Suppl**. **Fig.2E-1 and F-1**), although when averaged across all days in LD, that effect for specific variants was not significant (**Suppl. Fig. 3E-2 and F-2**). Second, a delayed Evening phase was also produced by two of the Simple PDFR variants (**Suppl. Fig. 2M**), the 6A and 7A variants, each of which contain only a pair of Ser-to-Ala substitutions. The amplitude of the Morning peak (ZT18-20) was increased modestly by variants 2-3A, 7A and 1-5A (**Suppl. Fig 3C-1, G-1** and **J-1, red arrows**) or strongly by 6A (**Suppl. Fig. 3F-1, red arrow**). The amplitude of the Evening peak was also increased by others in the PDFR variant series (**Suppl. Fig. 3, blue arrows**), of which the 6A, 7A, 1-5A and 1-6A variants had effect sizes most similar to that of 1-7A (**Supp. Fig. 3F-3. G-3. J-3 and K-3).** We also note that combining the Single 5A, 6A and 7A variants into a Multiple Variant (5-7A) did not produce the anticipated additive effects on increasing Morning peak amplitude at ∼ZT19 (**Suppl. Fig. 3H-1),** nor on increasing Evening peak amplitude **(Suppl. Fig. 3H-3).** Such anticipation was predicated on the effects seen individually with the 1-5A, 1-6A and 1-7A variants (versus the 1-4A); instead the 5-7A produced only a modest increase in the size of the delayed Evening peak (**Suppl. Fig. 3H-3**).

### Effects of WT versus 1-7A PDFR on locomotor behavior under equinox conditions (Fig. 3)

Under 12:12 conditions, *han* mutant flies (lacking *pdfr* function) typically display elevated nocturnal activity, a lack of morning anticipation prior to Lights-on, and a pronounced advance in the peak of the Evening behavior [11]. Although we note that some reports have observed remnants or a full bout of Morning activity in *han* mutant flies [e.g., 37]. Fig. 3A-1 displays activity patterns of *han* mutants that are heterozygous for *tim*-Gal4; they generally matched prior descriptions of *pdfr* mutant behavior. Restoration of *pdfr* function by *han*; *tim*> WT-*pdfr*, restored a Morning peak (anticipatory activity prior to lights-on – Fig. 3B-1 **and -2**) and delayed the evening peak by 1-2 hrs (Fig. 3B-1 **and** B-4) [7,8,9]. Restoration of *pdfr* function by the 1-7A PDFR variant under these conditions (Fig. 3C-1) increased the amplitude of the Morning activity peak but not its phase (Fig. 3C-2**, red arrow**) but not its phase (Fig. 3D). 1-7A expression in equinox conditions rescued the advanced phase of the Evening activity peak, but did not further delay it past Lights-OFF (Fig. 3C-4 and 3E), unlike what we observed under short-day conditions. The amplitude of the Evening peak was only modestly elevated following expression of WT or 1-7A PDFRs. The period (tau) of the 1-7A variant was considerably lengthened, but see discussion below about the influence of the GFP fusion. The phase of the major DD rhythm was delayed relative to the evening peak under entraining conditions (Fig. 3H, green line). In sum under equinox-like photoperiods, the PDFR 1-7A variant significantly increased the amplitude of the morning activity peak compared to WT PDFR, but did not differentially affect its phase, or the amplitude or phase of the Evening peak; under subsequent DD, the rhythmic activity displayed a delayed phase starting within the very first cycle.

### Behavioral effects of the other PDFR Variants under equinox conditions (Suppl. Fig. 4 & 5)

We tested each of the nine other PDFR Ala-variants by comparing their effects on locomotor behavior as we had done for the PDFR 1-7A variant. The phases of the Morning and Evening peaks were not obviously or systematically effected (**Suppl. Fig. 4L and 4M**). Relative to WT PDFR expression, the Morning Activity peak amplitude was increased by expression of the 6A, 7A and 1-5A variants (**Suppl. Fig. 5F-1, G-1**, and **J-1**, red arrows). Relative to WT PDFR expression, the Evening Activity peak amplitude was increased by expression of the 4A, 6A, 7A, 5-7A and 1-5A variants (**Suppl. Fig. 5D-3, F-3, G-3**, **H-3** and **J-3**, blue arrows). We also note that combining the Single 5A, 6A and 7A variants into a Multiple Variant (5-7A) did not produce the anticipated additive effects on increasing Morning peak amplitude at ∼ZT21 (**Suppl. Fig. 5H-1, versus 5F-1 and G-1).**

### Behavior under Long Day (summer) conditions (Fig. 4)

Flies lacking *pdfr* function in 16L:8D conditions display locomotor patterns similar to those in 12L:12D – lack of a clear Morning peak and display of a broad Evening activity period that peaks ∼2-3hr before Lights-off: the example in Figure 4A-1 **(yellow box)** displays behavior by *han* mutants that are also heterozygous for the *tim*-Gal4 element. Rescue (*han*; *tim*> WT-*pdfr*, Figure 4B-1**, black box**) typically delayed Morning activity offset (Fig. 4B-3 **and** 4D) and delayed the Evening peak onset each by 1.5 h (Fig. 4B-4 **and** 4E). Relative to that of WT PDFR, expression of the 1-7A variant (Figure 4C-1) increased the amplitude of both the morning activity peak (Figure 4C-3) and the evening activity peak (Figure 4C-4). It did not significantly alter the Morning phase relative to effects of WT PDFR (Figure 4D), although there was a pronounced tendency to broaden the duration of the Morning activity peak. 1-7A expression did significantly extend the delay the Evening phase of activity relative the effect of Wt PDFR (Figure 4E). In DD, the 1-7A PDFR variant significantly lengthened circadian period and the dominant rhythmic activity displayed a phase several hours delayed from the evening activity peak phase under light entraining conditions within the first cycle (Fig. 4H – green bar). In summary the 1-7A PDFR variant significantly increased the amplitudes of the Morning and Evening activity peaks under long days, and also slightly delayed the Evening activity phase.

### Behavioral effects of the other PDFR Variants under long day (summer-like) conditions (Suppl. Fig. 6 & 7)

We tested each of the nine other PDFR Ala-variants by comparing their effects on locomotor behavior as we had done for the PDFR 1-7A variant. There was no systematic effect of the variants on the phase of the Morning peak (**Suppl. Fig. 6L**), but several did delay the Evening activity phase (**Suppl. Fig 6M**), especially the 1-4A, 1-5A and 1-6A variants. A few variants affected the Morning peak amplitude – 5A, 6A and 1-6A (**Suppl. Fig. 7E-2, F-2 and K-2**). Several increased the amplitude of the Evening peak under these conditions, namely 2-3A, 4A, 5A, 6A, 7A, 1-5A and 1-6A (**Suppl. Fig. 7 C-3, D-3, E-3, F-3, G-3, J-3** and **K-3**). As observed in the other photoperiodic conditions, when we combining the 5A, 6A and 7A variants into a single 5-7A variant, it did not produce the expected additive effects on behavior in the morning (**Suppl. Fig 7H-2**) or in the evening (**Suppl. Figure 7H-3**): instead these activities appeared similar to effects displayed by the 1-4A variant.

### Effects of PDFR variants on period and phase of rhythmic activity in DD

In DD, following Short Day conditions, the PDFR variants that produced 3-4 h delays in the evening peak often generated periods ∼ 1-2 hr longer than the controls and also lowered % arrhythmicity (Table 1). This was especially true for the Multiple Variant series (e.g., PDFR 1-5A, 1-6A and 1-7A (Table 1). However the correlation between delayed evening peaks and a longer tau in DD was not absolute. For example, the 6A, 7A and 5-7A variants all had strong and significant delaying effects on the Evening phase of activity under Short Day conditions (compared to the effect of the WT PDFR), but they did not lengthen tau values in subsequent DD conditions (Table 1). These observations suggest that delays of activity phases produced by PDFR modulation in LD may not reflect a direct consequence of PDFR effects on the PER and TIM--dependent clock [2, 3, 17]. In addition, the dominant activity periods in DD typically reflected the delayed (∼ZT12) Evening peak (**Suppl. Fig. 2F-2, 2G-2** and **2H-2**). Notably, the early Morning peak (∼ZT18-19) promoted by PDFR 6A and 7A variants clearly persisted in DD (**Suppl. Fig. 2F-2, 2G-2**). In DD following Equinox or Long Day entrainment (with photophases > 8 hr in duration), we noted that several variants produced dominant activity phases which were phase-delayed relative to the phase of the Evening peak displayed in LD (e.g., 6A and 7A - **Suppl. Fig. 4F-2 and 4G-2**; **Suppl. Fig. 6F-2 and 6G-2).** Again, several of the variants significantly increased Tau in DD (Table 1), yet had no delaying effects on phase in LD: examples included 2-3A, 1-4A, and 1-5A, under equinox conditions (**Suppl. Fig. 4C-2, 4I-2, 4J-2**) and 4A and 1-5A in long day conditions (**Suppl. Fig. 6D-2, 6J-2).**

Behavioral activity records from days 3-9 in constant conditions following entrainment in the three indicated photoperiods (Short Day, Equinox and Long Day). N = number of independent experiments performed for each genotype::photoperiod combination. n = total number of flies tested for each genotype::photoperiod combination. % AR: the percentage of flies judged arrhythmic by criteria; tau = average circadian periods calculated according to χ2-periodogram analysis. * complete genotype = *han*[5304]*; tim > w*[1118]*. \*\** complete genotype = *han*[5304]*; tim > pdfrEgfp*. Statistical analysis compared tau’s using Tukey’s multiple comparisons post hoc test following a one-way ANOVA: ns: not <u>significant; * p < 0.05; ** p < 0.01; *** p < 0.001; **** p < 0.0001.; nd: not determined.</u>

### Control experiments

The design of the *PDF*R variants contains assumptions and genotypic constraints, distinct from simply substituting Alanine at potential sites of phosphorylation in the receptor C terminal tail. To assess the potential of some assumptions to affect the results we report, we performed the following two sets of experiments as controls. ***Control experiments (i) - dependence of GPCR PDFR variant effects on the presence of PDF ligand***. We assumed that the altered behavioral phenotypes produced by certain PDFR variants depended on activation by their endogenous cognate ligand, the neuropeptide PDF. However, we could not *a priori* exclude the possibility that PDFR variants may in fact produce novel constitutive activity (neomorphic properties). We therefore placed the 1-7A PDFR variant in the *pdf^01^* (null) background and re-tested its activity when driven by *tim*-Gal4 under both short day and long day conditions, and then in constant darkness. In all conditions, the behavioral phenotypes largely resembled those of the *pdf^01^* background (lack of a morning activity peak, advanced evening peak, and shorter tau/s in DD). The results are shown in Fig. 5 (under short days) and Fig. 6 (under long days) and tabulated in Table 1. The ability of the 1-7A PDFR variant to increase the amplitude of the Morning peak was completely dependent on WT *pdf* function (Fig. 5B-2). Likewise its ability to significantly delay the Evening phase into the dark period was completely dependent on WT *pdf* function (Fig. 5B-4 **and** 5E). Under Long days, the ability of the 1-7A variant to increase the amplitudes of the Morning and Evening activity peaks were strongly diminished by lack of WT pdf function (Fig. 6B-3 and B-4). The 1-7A variant did display some activity in the *pdf* mutant background: for example, it produced a significant delay of the Evening peak phase (akin to the action of the WT receptor, Fig. 5A-3 and Fig.6 A-4) and a reduction in the % arrhythmicity to nearly the same value as found in a WT *pdf* background. These results suggest some degree of constitutive (ligand-independent) activity. Apart from these, we conclude that constitutive activity explains at best a small proportion of the behavioral effects that distinguish the 1-7A PDFR variant from the WT receptor.

**Figure 5.**
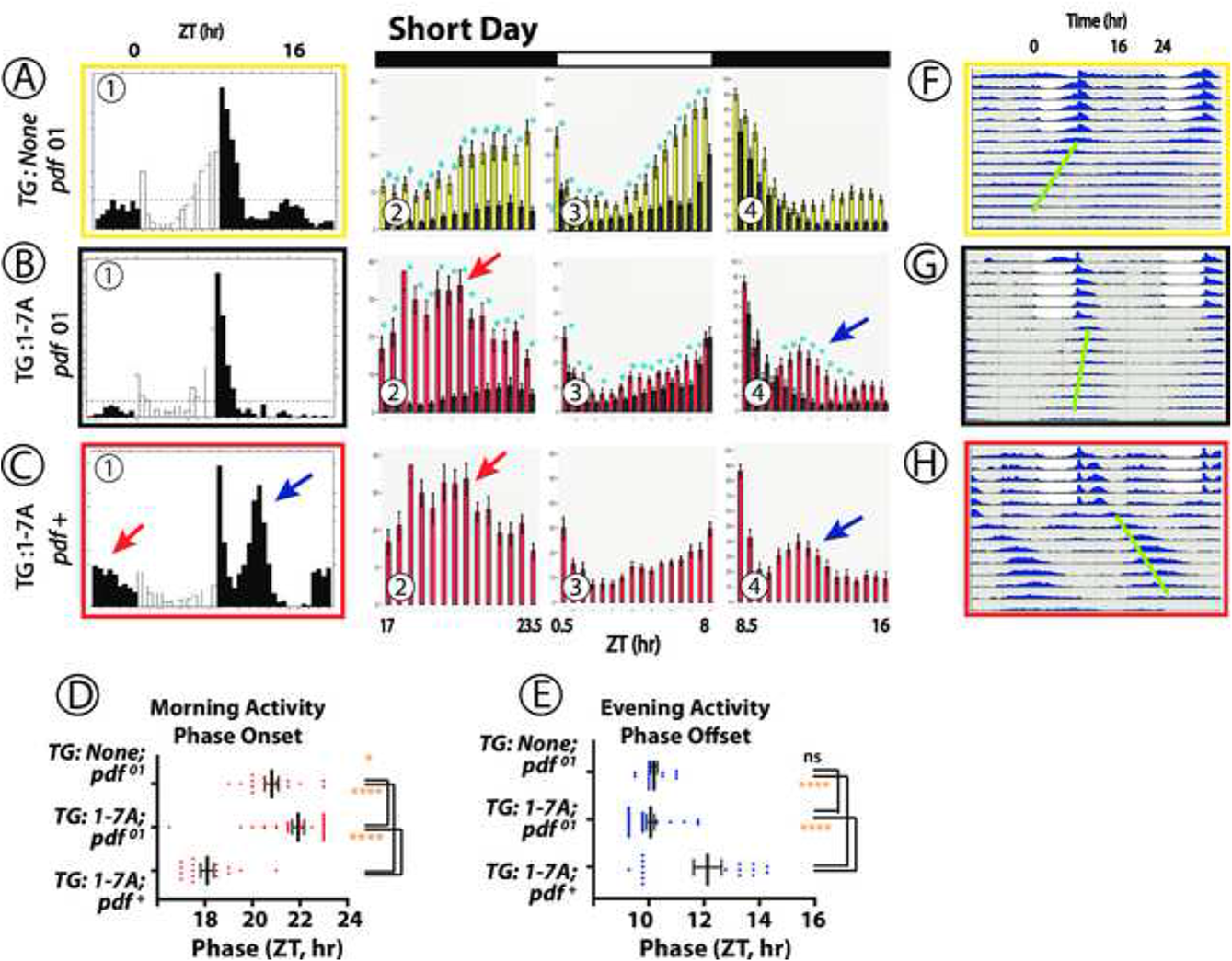
Locomotor Rhythms exhibited by WT versus *pdf* mutant flies expressing the 1-7A PDFR Variant under Short-day (winter-like) conditions. The Panels present behavioral activity with a format similar to that described in Figure Legend 2. Behavioral records were recorded from *pdf^01^* (null) flies that expressed either no UAS transgene (A, yellow box), or a UAS-1-7A *pdfr* transgene (B, black box), or from control flies (*pdf^+^)* that expressed the UAS-1-7A *pdfr* (C, red box)). *TG*: transgene. Panels (A)-(C) display average group eductions (activity averaged over days 5 and 6 of light entrainment, with open bars indicating the 8-hr periods of Lights-on and filled bars the 16-hr periods of Lights-off) and bin-by-bin analyses to compare activity amplitudes between genotypes. The red and blue arrows indicate Morning and Evening activity of control (red) flies with distinguished amplitude and/or phase, respectively. Blue asterisks mark amplitudes that are significantly different between genotypes by ANOVA followed by a Student’s T-test (p < 0.05). Panels (D) and (E) display the phases of Morning activity Onsets and Evening activity Offsets respectively. Gold asterisks: ANOVA followed by Dunnett’s post hoc multiple comparisons of all compared to WT: ns = not significant; * = p<0.05; ** = p<0.01; *** = p<0.005; **** = p<0.001. Panels (F) – (H) display double plotted actograms of activity during the ∼6 days of light:dark conditions, followed by activity during ∼9 days of DD. The green bars indicate the phase of the dominant activity period in DD.

**Figure 6.**
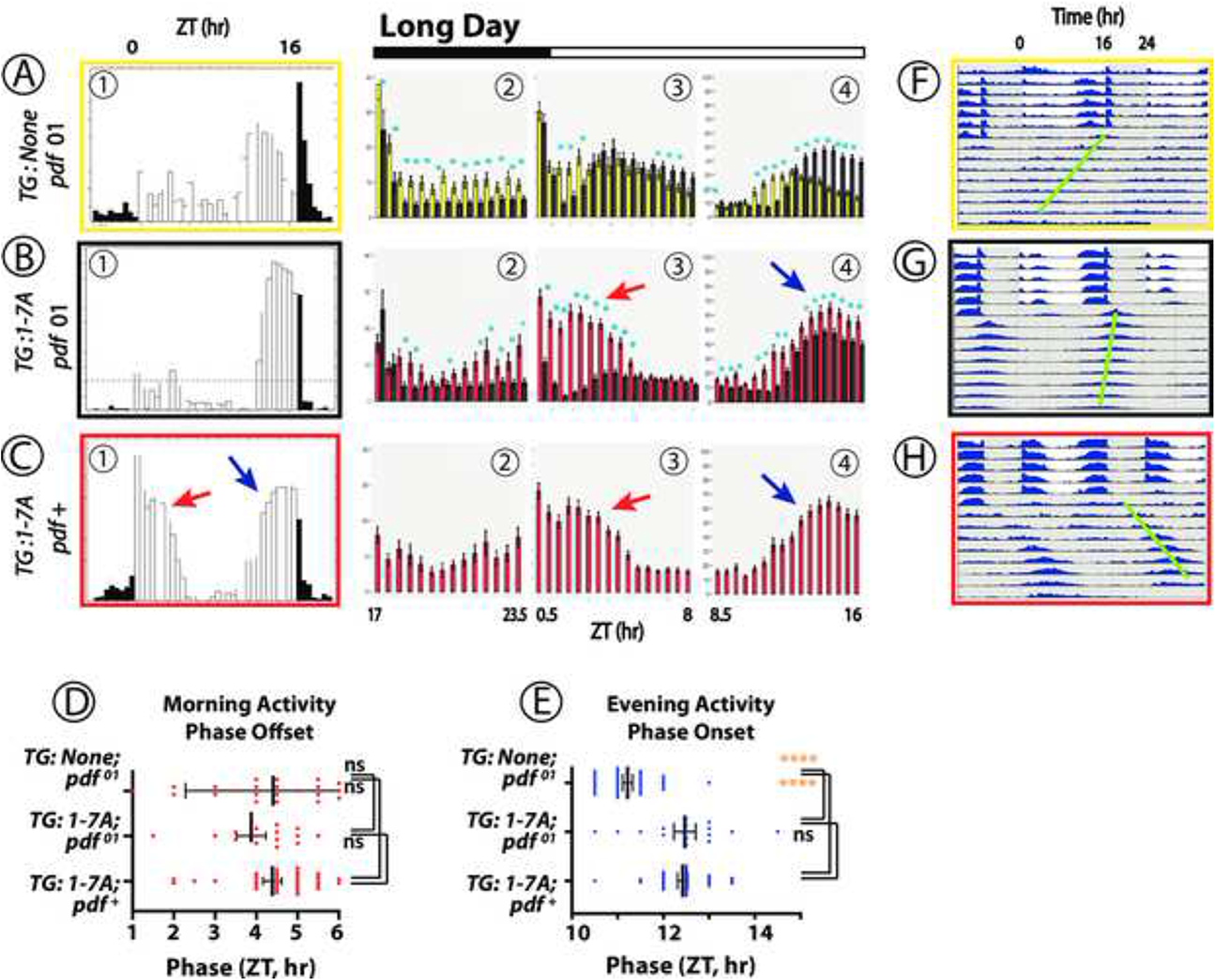
Locomotor Rhythms exhibited by WT versus *pdf* mutant flies expressing the 1-7A PDFR Variant under Long-day (summer-like) conditions. The Panels present behavioral activity with a format similar to that described in Figure Legend 5. Behavioral records were recorded from *pdf^01^* (null) flies that expressed either no UAS transgene (A, yellow box), or a UAS-1-7A *pdfr* transgene (B, black box), or from control flies (*pdf^+^)* that expressed the UAS-1-7A *pdfr* (C, red box)). *TG*: transgene. Panels (A)-(C) display average group eductions (activity averaged over days 5 and 6 of light entrainment, with open bars indicating the 16-hr periods of Lights-on and filled bars the 8-hr periods of Lights-off) and bin-by-bin analyses to compare activity amplitudes between genotypes. The red and blue arrows indicate Morning and Evening activity of control (red) flies with distinguished amplitude and/or phase, respectively. Blue asterisks mark amplitudes that are significantly different between genotypes by ANOVA followed by a Student’s T-test (p < 0.05). Panels (D) and (E) display the phases of Morning activity Offsets and Evening activity Onsets respectively. Gold asterisks: ANOVA followed by Dunnett’s post hoc multiple comparisons of all compared to WT: ns = not significant; * = p<0.05; ** = p<0.01; *** = p<0.005; **** = p<0.001. Panels (F) – (H) display double plotted actograms of activity during the ∼6 days of light:dark conditions, followed by activity during ∼9 days of DD. The green bars indicate the phase of the dominant activity period in DD.

### (i) The influence of epitope fusions

The second control experiment asked whether the 1-7A epitope fusions (4xFLAG and eGFP), might confer some of the differences in behavioral effects that we observed. In part, the concern regarding fused sequences is mitigated by the fact that all UAS-*PDFR* isoforms in the experimental series contained both epitopes (in addition – all were genetically transduced to the same integration site). Hence comparisons across different variants should largely normalize for the effects of the non-receptor sequences, and instead highlight variant-specific properties. However, we wished to test this assumption explicitly to define the extent to which a bulky GFP fused to the C Terminal tail might confer some of the behavioral properties we might otherwise ascribe to PDFR C terminal sequences. We therefore compared a 1-7A version of PDFR to a wild type PDFR, both of which lacked GFP fusions. We found that the additional Evening phase delays in short day and long day produced by 1-7A expression occurred despite the absence of a GFP fusion (**Supp. Fig. 8B-4, 8H** and **8J-4 and 8P**); in addition, the increase of the Morning amplitude under Long days occurred despite the absence of a GFP fusion (**Suppl. Fig. 8J-3**). Finally, the phase of the main DD rhythmic activity, following expression of 1-7A lacking a GFP fusion, reflected the delayed evening phase in short day (**Suppl. Fig. 8F**, green bar) and was delayed relative to that of the LD Evening activity peak in long days (**Suppl. Fig. 8N**, green bar). We note however that the longer period produced by the 1-7A variant was not exhibited by the 1-7A lacking GFP, suggesting that a lengthened period depends at least in part on the GFP fusion. In sum, these results support the hypothesis that most of the key alterations in daily locomotor rhythmicity derived from sequence variation of the PDFR, and not from the properties of the fused GFP.

### PDFR expression and signaling *in vitro*

The PDFR is a G_s_-coupled receptor and several reports have documented the importance of the downstream cAMP pathway to mediate PDF behavioral regulation [15, 17, 38]. In that context, we asked whether the *in vivo* properties of those PDFR variants that affected behavioral phase and amplitude could be correlated with *in vitro* properties when expressed in *hEK-293T* calls. We found no strong correlation. Significant differences in basal signaling levels, in EC50 or in maximum values of signal transduction were rare (**Suppl. Fig. 9, 10 and 11**). When noted, they were poorly correlated with behavioral effects (**Suppl. Table 5**). We also measured surface expression of the PFDR variants in *hEK* cells using β-lactamase N terminal fusions [39] to determine if PDFR variants tended to display longer surface lifetimes. The 1-4 and 1-7 variants displayed higher basal levels, but no others were different from the WT levels (**Suppl. Fig. 12**). Following 20 min exposure to PDF, neither the WT not any of the variants displayed a change in surface expression levels (**Suppl. Fig. 1**3).

### Measuring the phosphorylation state of over-expressed PDFR *in vivo*

To obtain direct evidence that PDFR sequences are phosphorylated *in vivo*, we first over-expressed an epitope-tagged-PDFR WT construct using *tim*-Gal4, which directs expression broadly in cells that feature the PER-dependent molecular oscillator. We immunoprecipitated the receptor from head extracts, then employed tandem mass spectroscopy to determine which if any specific PDFR peptide fragments are phosphorylated. We performed eight biological replicates, with two collections in the morning (ZT2-3) and six in the evening hours (ZT11). **Suppl.Table 4** reports the phosphorylated peptides detected from the PDFR-GFP fusion protein. Among the 28 conserved Ser/Thr and Tyr residues in the PDFR C terminal tail, five were phosphorylated in one or more of these samples. In the two Mornings samples S563 was phosphorylated in one sample; in the Evening samples, S531 (in CL2) was phosphorylated in four of samples, S534 (in CL2) in one, T543 (in CL3) in one, and S560 in two of six samples. The spectra documenting detection of these phosphopeptides are presented in **Suppl. Fig. 14**.

### Testing GRKs and β-arrestin2 contributions to locomotor rhythmicity

Regarding which potential PDFR phosphorylation sites might be β-arrestin binding sites, we considered a recent report by Zhou *et al.* [40], who proposed a conserved GPCR sequence motif that promotes high affinity interactions with β -arrestin. The motif includes three phosphorylation sites that align with three conserved, positively-charged pockets in the arrestin N-domain. In *D. melanogaster* PDFR, the CL1 region contains a match with the proposed motif beginning with S512 (SLATQLS) and shows moderate sequence conservation: of the 17 species we considered, 9 others retained this precise motif (**Suppl. Fig. 1**). In addition, the *D. melanogaster* PDFR sequence beginning with S629 (SRTRGS) also displays a match for the motif, but is retained in only 6 of the other 16 species. The evidence that these two sites represent high affinity β -arrestin2 binding domains is therefore equivocal. We previously reported that β-arrestin2-GFP is not efficiently recruited to activated PDFR when functionally-expressed in *hEK-293T* cells [29]. In contrast, each of 13 other *Drosophila* neuropeptide GPCRs do efficiently recruit β-arrestin2-GFP when expressed and activated in that cellular environment [30–32].

We tested the potential to detect *in vivo* involvement by *Drosophila* orthologues of the canonical desensitization effectors - mammalian G-protein coupled receptor kinases (GRKs – GPRK1 (CG40129) and GPRK2 (CG17998)) and of mammalian β-arrestin2 (βarr2 – *kurtz* (CG1487)) - in the control of locomotor rhythms We drove specific over-expression of WT cDNAs and RNAi constructs using *tim*-Gal4, to broadly affect signaling in the circadian pacemaker system. The PDFR-PA protein is highly restricted in its expression to subsets of the pacemaker neural network [9]. We predicted that elimination of a desensitizing component for PDFR signaling should produce behavioral phenotypes opposite to that of *pdfr* loss of function phenotypes: these would potentially include (for example) a delayed evening activity phase in LD conditions and a longer tau under DD. GPRK1- and *krz*-specific RNAi’s produced normal average locomotor profiles under 12:12, while one of two GPRK2-specific RNAi constructs tested slightly advanced the evening peak (**Suppl. Fig. 15**). Over-expressing GPRK1 cDNAs did not affect morning or evening phase; over-expressing GPRK2 broadened the evening peak. Under DD, *krz* RNAi flies were uniformly arrhythmic, while GPRK RNAi’s were normal or slightly lengthened the circadian period (**Suppl. Table 5**). These results do not support the hypothesis, and suggest that the kinetics by which PDFR signaling terminates do not depend exclusively on the activities of either dGPRK- 1 or -2, or on that of the *Drosophila* β-arrestin2 ortholog, *krz*.

## DISCUSSION

We employed a gain-of-function approach to measuring neuropeptide GPCR function *in vivo*. In doing so, we sought to learn about the maximal consequences of PDFR GPCR signaling and also learn something about the phosphorylation events that normally control the duration of PDFR’s signaling lifetime. We found that expression of a PDF receptor variant, containing numerous Ala-substitutions of conserved phosphorylatable residues in the C terminal domain (1-7A), fundamentally altered rhythmic locomotor behavior in *Drosophila*. The changes included increases in the amplitudes of the Morning and/or Evening activity peaks, delays in their phases and/ or lengthening of the free-running period. Such results are generally consistent with a prediction whereby non-phosphorylatable PDFR variants – those with a potential to increase the duration of PDFR signaling - would produce behavioral actions opposite to those seen in loss-of-function *pdf* [1], or *pdfr* [11] mutant stocks. Accordingly, we speculate that phosphorylation of the PDFR at these conserved residues normally defines the duration of its active signaling state each day. The longer duration that follows the substitutions of Ala may allow receptor signal strength to also increase by accrual: such an effect could explain the increased amplitude of the Morning and Evening peaks, as seen with expression of some of the variants. In this context, we acknowledge that modification of GPCRs by phosphorylation does not exclusively lead to signal termination. GRK2-dependent phosphorylation of the smoothened (smo) GPCR follows reception of the Hh signal and helps mediate its signal: phosphorylation leads to smo <u>activation</u> in both *Drosophila* and mammalian systems [41]. Thus modification of putative phosphorylation sites on GPCRs (to preclude phosphorylation) will not exclusively promote extended signaling. Therefore our interpretations must correspondingly consider outcomes without *a priori* assumption of mechanisms.

### GPCR signal termination

PDFR belongs to the Secretin Receptor Family (Family B) of neuropeptide receptors [11–13]: there is no clear consensus regarding mechanisms of desensitization and internalization for this receptor family. For VPAC2 receptors, phosphorylation and internalization is mediated exclusively by GRK2 [42]. Likewise, *Drosophila* orthologues of Cortocotrophin Releasing Factor receptors (CG8422, DH44-R1) and Calcitonin receptors (CG17415, DH31-R1) are internalized in *hEK* cells following recruitment of β-arrestin2 [31–32]. In contrast, PDFR, which is also related to the mammalian Calcitonin receptor, is not internalized following exposure to PDF [29]. VPAC2 receptors are also regulated by heterologous receptor signaling: M3 cholinergic receptors via PKC signaling can block VPAC2 phosphorylation, desensitization and internalization [43]. Furthermore, secretin receptors and VPAC1 receptors undergo phosphorylation by GRKs and β-arrestin2-dependent desensitization, but these are not sufficient to facilitate or mediate internalization [44–46]. Finally, GLP2-Receptor associates with β-arrestin2 via its distal C terminal sequences, but that receptor domain is required neither for GLP2-R desensitization nor its internalization [47]. Thus kinases other than GRKs and effectors other than β-arrestin2 may regulate internalization of diverse Family B receptors.

Following activation of rhodopsin in the mammalian retina, visual arrestin is recruited with a time constant of < 80 ms [48]; for many *Drosophila* neuropeptide receptors, β-arrestin2 is recruited within a minute of exposure to ligand [30]. The PDFR GPCR signals over a time base of many hours [19]: could phosphorylation/de-phosphorylation also regulate its activity? The mammalian blue-light sensitive GPCR melanopsin (OPN4) mediates intrinsic light sensitivity in certain classes of retinal ganglion cells (RGCs): melanopsin signaling is distinguished by long latencies, graded responses and sustained RGC depolarization that can outlast the duration of the light stimulus [49]. Melanopsin-expressing RGCs produce exceptionally long time-course integration because that GPCR behaves differently from other opsins in two fundamental ways. Firstly, it undergoes photoequilibration between signaling and silent states: a property that maintains the availability of pigment molecules for activation, termed ‘tristability’ by Emmanual and Do [50]. Secondly, phosphorylation and β-arrestin recruitment do contribute to the kinetics of melanopsin signaling, but over a long time-course [51–52]. Mure *et al.* [34] proposed that the distal portion of the melanopsin C terminal tail creates a steric blockade covering a cluster of specific serine residues situated in more proximal regions. The time required for relief of that blockade dictates the time course of GPCR phosphorylation and therefore delays subsequent desensitization.

### Specific versus non-specific phosphorylation regulating GPCR activity

The 1-7A PDFR variant modified the greatest number of phosphorylatable residues in the series we tested (Fig. 1), and typically produced the strongest behavioral effects. Notably, the 6A variant – which modified only two specific Ser residues – produced effects that were nearly identical to those exhibited by flies expressing the 1-7A form. This suggests that much PDFR post-translational modification, that which is capable of restricting its signaling time-course, may be directed to specific phosphorylation sites near the extreme C terminal end. There are two broadly divergent hypotheses to describe the mechanisms by which GPCR phosphorylation promotes desensitization. The first proposes that modification of specific residues have the greatest significance for downstream desensitizing mechanisms: such a mechanism can explain the effects we have seen with the highly limited 6A PDFR variant. The second hypothesis invokes triggering of a termination processes by the aggregate negative charge accumulated with bulk phosphorylation, regardless of where it might occur along a GPCR’s intracellular sequences. For some GPCRs (e.g., OPN4), the evidence supports both models [34, 35]. Our data concerning PDFR desensitization also support both viewpoints and we present a model in Figure 7. In particular we regard CL6 and CL7 to be the specific residues especially critical in terminating the time course of PDFR signaling. Notably, converting just the single pair of CL6 AAs, or the single pair of CL7 AAs, is enough to generate hours-long delays in the peaks of locomotor rhythms under Light:Dark conditions, as well as generating a significant increase in tau under Constant Dark conditions. We note that the splicing event that distinguishes the PDFR-A and PDFR-D isoforms occurs just prior to the position of sequences encoding CL6 and 7, such that the D form lacks these two highly conserved domains (<u>flybase.org/download/sequence/FBgn0260753/FBpp</u>). Other clusters (like CL2-3) also appear to have potential for ‘specific’ contributions to PDFR regulation. Together, these results speak to the potency of the inferred <u>specific</u> termination mechanisms for PDFR GPCR signaling. In contrast, the evidence for the bulk phosphorylation hypothesis comes from comparison of the 1-4A, versus the 1-5A, 1-6A and 1-7A variants. This series inactivates increasing numbers of phosphorylatable residues, and with it we observed increasingly delayed evening phases in short day conditions (e.g., **Suppl. Fig. 2I-2, 2J-2 and 2K-2**). The tandem mass-spectroscopy results, indicating endogenous phosphorylation of certain PDFR residues (**Supplemental Table 4**) demonstrates such post-translational modifications can occur. However, we caution that these phospho-peptide measurements derive from whole head extracts and may not provide a complete or relevant accounting of those sites that are modified in critical E pacemaker neurons.

**Figure 7.**
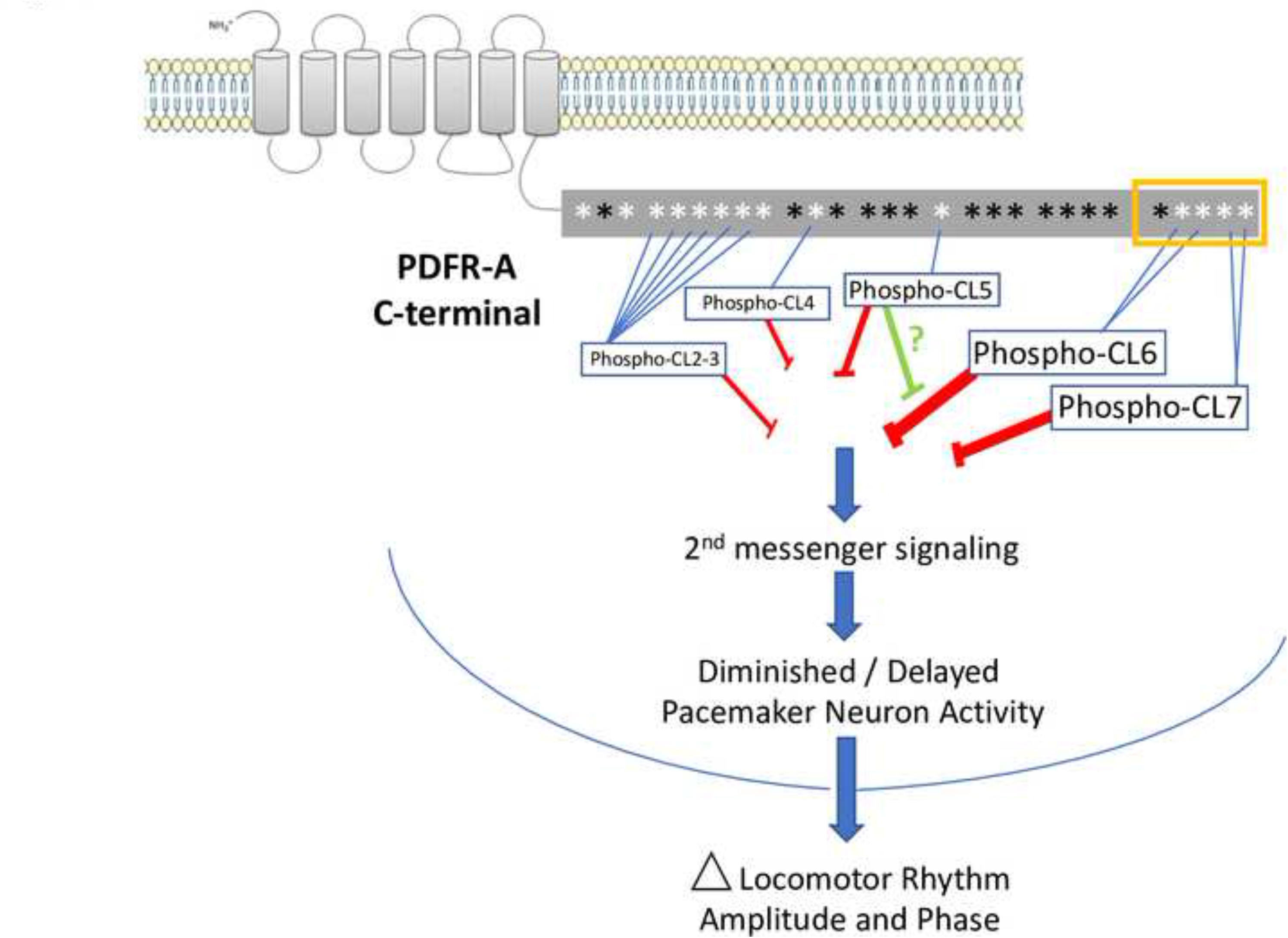
A model predicting the effects of phosphorylating different sites on the PDFR CT on downstream signaling. The CT is diagrammed as a grey box running horizontally. Asterisks indicate the positions Ser/ Thr/ and Tyr residues as described in Fig. 1 and Suppl. Fig. 1: white asterisks are the subset of Ser/Thr/ and Tyr residues that were targets of mutational analysis in this study. Mutation of CL6 and CL7 consistently demonstrated the greatest delaying effects on the phases of Evening locomotor peaks; they were also among the sites with greatest influence on the amplitudes of the Morning and Evening peaks. The results suggest their phosphorylation normally will have the greatest effect to slow or terminate PDFR signaling. Other sites are also effective, although to lesser degrees (as indicated by their font sizes), including CL2-3, CL4 and CL5. Primarily, such phosphorylation will decrease the duration of PDFR signaling (red bars) and so reduce (for example) the delay that PDF imposes on the period of neuronal activation displayed by PDFR-responsive pacemaker groups (like the Evening cells and the DN3 [10, 19]). The model also predicts that the effects of phosphorylating CL5 will depend on the phosphorylation status of neighboring sites. In some contexts (e.g., PDFR 1-5A), it will help terminate PDFR signaling, but in others (e.g., PDFR 567A) it may promote the duration or extent of PDFR signaling, perhaps by blocking the effects of phosphorylating CL6 and CL7 (green bar).

Our results also point to what we propose as “context-dependent” effects of modifying specific GPCR residues – ones that may produce opposing behavioral effects, depending on the phosphorylation status of neighboring sites. In particular we point to the opposing results we observed with Ala-variants of the CL5 site. In comparing results from the series 1-4A, 1-5A, and 1-6A, we found that under Short Day conditions, the 1-4A had only mild effects on the evening peak phase compared to over-expressing WT PDFR (e.g., **Suppl. Fig. 2B-1** vs. **2I-1**), whereas 1-5A and 1-6A both produced significant delays (**Suppl. Fig. 2M and Suppl. Fig. 3J-3 through 3K-3)**. The difference between the 1-4A and 1-5A variants is a change in a single S residue at CL5 (S633A): Those observations suggest modification of the CL5 residue normally promotes desensitization of PDFR and termination of PDFR signaling. However, the PDFR variant 5-7A was constructed with the expectation that it would be as effective as 1-5A, 1-6A or 1-7A in delaying the evening phase. Instead it proved only weakly effective. This same mis-match of expected behavioral effects for 5-7A was also seen in other photoperiodic conditions (**Suppl. Fig. 6M).** Likewise, Langlet *et al*. [36] reported that effects on mutating Serine residues in the carboxy terminus of VPAC1 did not produce additive effects. We propose that phosphorylation of CL5 will have either positive or negative consequences on PDFR desensitization depending on which other neighboring residues are also modified. Thus we speculate that CL5 takes on outsize importance in determining desensitization rates for PDFR and so may itself be subject to exceptional regulation. Such complex interactions between phosphorylation sites is reminiscent of interactions documented between diverse phosphorylation sites in the circadian clock protein PERIOD [53–54]. Better resolution of these two paradoxical mechanisms ((i) specific phosphorylation versus bulk negative charge, and (ii) context-dependent effects of CL5 site phosphorylation, awaits precise molecular definition of where and when PDFR is modified in key pacemaker neurons *in vivo*, and by which post-translational modifications.

### Validating the behavioral phenotypes produced by PDFR variants

The actions of PDFR variants we have described, following substitutions of Ala for various Ser/Thr or Tyr residues, are not easily explained by a hypothesis invoking neomorphic or constitutive GPCR properties. The evidence for this conclusion is three-fold. First, the effects on both amplitude and phase of activity peaks by these variants are not of a random assortment: rather they are all strictly opposite to that of loss-of-function models for *pdf* and *pdfr*. Second, the actions of PDFR variants display strong dependence on wild type *pdf* gene function: this strongly argues that the actions of PDFR variants reflect responses to normal, endogenous PDF signaling. Third, even the strongest actions of PDFR variants on the phases and amplitudes of locomotor activity (that of the 1-7A variant) reflected PDFR sequence variation, and not the properties of the GFP C-terminal fusion. Together these observations support the hypothesis that phosphorylation of some or all of the C terminal residues we studied are normally modified to attenuate the strength and duration of PDFR signaling during the 24-hr day.

### PDF Signaling and seasonal adaptation

PDF neuropeptide actions were discovered in the context of physiological adaptions to daylight in crustacea [5, 6]. Helfrich-Förster [55] discovered PDF expression within a defined subset of the insect *Drosophila* circadian pacemaker network. Renn *et al.* [1] demonstrated that *pdf* makes a fundamental contribution by setting the normal behavioral phase of Evening activity in *Drosophila* during light:dark entrainment and promoting normal rhythmicity during constant darkness. PDF signaling coordinates with that driven by light: It works in parallel to direct photosensitivity along with the CRY blue light photoreceptor. Flies doubly-mutant for *cry* and *pdf* display pronounced deficits of locomotor rhythmicity [7, 8, 9], and can be as severe as measured in clock-deficient flies. PDF and environmental light also work coordinately at the level of neuronal activity patterns: in the case of the evening pacemakers (LNd and the 5^th^ s-LNv), both light and PDF signaling promote a delay in their PER-dependent activation period, and together help align it to a phase just prior to dusk [19]. Based on the close association of PDF signaling and photoperiodic signaling, it is not surprising that the duration of the photoperiod strongly influenced the effects of PDFR sequence variants on the Morning versus the Evening activity peaks. We only observed delayed Evening activity peak produced by PDFR gain-of-function variants under Short Day conditions: that delay appears strongly inhibited by light durations > 8 hr. It remains unclear at what molecular level light exerts such an inhibitory effect, however, we note a similarity of these observations with effects recently reported from loss-of-function states for the phosphatase *PRL-1* [56]. Like the most active PDFR variants we herein describe (e.g., 6A, 7A, 1-5A 1-6A and 1-7A), *PRL-1* mutants produce a 3-4 h delay in the phase of the evening activity peak, but only under winter-like (short day) conditions, and not under equinox or summer-like (long day) conditions. Kula-Eversole *et al.* [56] have shown that TIM phosphorylation is affected by PRL-1 activity and suggest the seasonal action of PRL-1 to advance the evening locomotor activity phase is mediated by modification of TIM levels within the transcription-translation feedback loop. The extensive similarity of these two sets of behavioral phenotypes reveals either serial or parallel pathways to effect comparable outcomes on locomotor activity. If serial, then according to the simplest model, PRL1 acts downstream of PDFR, and the combined results to-date suggest PRL-1 would be inhibited by PDFR activation. Further genetic and biochemical experiments are needed to evaluate if and how these pathways converge.

### Relations between in vitro and in vivo PDFR signaling measures

Using an *in vitro* assay for cyclic AMP generation, we found that modifying phosphorylation properties of PDFR does not affect the strength of signaling. That conclusion suggests that *in vivo* other GPCR features are normally affected to regulate the extent/duration of PDFR signaling according to season. This speculation corresponds to findings described with other Family B GPCRs like VPAC1 [46]: Mutation of all the Ser and Thr residues of the C terminal tail, and of Ser250 to Ala, led to a receptor with binding properties and adenylate cyclase activity similar to the wild type receptor; however that variant receptor was neither phosphorylated nor internalized. We propose that the PDFR variants we have described do not modify locomotor behavior by virtue of greater second messenger signaling. Rather they do so by generating cAMP (and perhaps other second messengers) over time periods longer than that normal sustained by a WT receptor over a portion of each 24-hr cycle.

### Is PDFR modulation of diurnal phase independent of its modulation of circadian period?

Many PDFR variants strongly delayed the peak of evening activity (past lights-off) and also significantly lengthened the circadian period (measured under constant darkness). A conventional interpretation of these linked phenotypes is that PDFR signaling affects the pace of the molecular clock (re-setting the period) and consequently affects behavioral phase indirectly, which is downstream of the clock. This explanation holds for all the variants that both increased tau and delayed the evening phase (e.g., 1-5A, 1-6A, and 1-7A). However, the correlation was not always observed. For example, under winter-like conditions, the 5A, 6A, 7A and 5-7A variants all significantly delayed the offsets of Evening activity peaks versus that seen with WT PDFR, but these variants did not significantly lengthen circadian period. Likewise, under long day, 6A produced a significant increase in tau versus that seen with WT PDFR, but did not significantly delay the onset of the Evening activity peak. Therefore, we cannot rule out an alternative hypothesis: that the duration of PDF>PDFR signaling to target pacemakers in the *Drosophila* brain is modulated each day to effect (i) entrainment of the molecular clock, and independently (ii) to directly delay to Evening pacemaker neuronal activity. In fact, the *pdfr han* mutant flies exhibited quasi-normal tau values in this report (Table 1), although they more traditionally exhibit shortened ones in our experience [37]. Evening locomotor activity derives in large part from the activity of the primary Evening oscillators, termed E cells, the LNd and the 5^th^ small LNv [10, 20, 57 – 59]. However, numerous network interactions are also known to provide critical contributions to the accuracy, precision and adaptability of that timing system [e.g., 59 – 64]. The Evening oscillator group itself is known to be heterogeneous and to constitute at least three separate, functionally distinct oscillators [60], only some of which express PDFR-A and respond to PDF [37, 60]. Vaze and Helfrich-Förster [65] reported that the phase of Period protein accumulation in E neurons is phase-locked to the previous lights-off transition and also sensitive to PDF signaling. They conclude that the peak of PER accumulation is key to determining the phase of the evening activity peak, but also note that the correlation between the Period-clock timing cue and the Evening activity peak is not perfect. They suggest the Evening behavioral phase is better described by also factoring in the delay in E cell neuronal activity driven by PDF neuromodulation [10, 19]. We speculate that it is the combination of two separate PDFR signaling effects that is essential for proper rhythmic behavioral outcomes across seasons. If valid, the hypothesis predicts a bifurcation of signaling pathways downstream of PDFR, to independently regulate diurnal phase and also circadian period within E pacemaker neurons.

## MATERIALS and METHODS

### Fly Rearing and Stocks

*Drosophila* were raised on a cornmeal agar diet supplemented with yeast at 25°C in 12 hr:12 hr LD cycles. The UAS-*pdfr* mutant transgenic series was created by injecting *yw P{nos phiC31\int.NLS}X;P{CaryP}attP40* embryos (Rainbow Transgenic Flies, Inc, Camarillo, CA). *pdfr* mutant flies are described in [11]; *tim*(UAS)-Gal4 flies are described in [66] – *BL80941*). The recombinant fly stock - *han^5304^*; *tim* (UAS)-Gal4 - was confirmed by PCR and sequencing. The UAS-*pdfr*-tandem construct was injected into *y*[1] *w[*] P{y[+t7.7]=nos-phiC31\int.NLS}X; P{y[+t7.7]=CaryIP}su(Hw)attP6* embryos. For PDFR-tandem fusion protein expression, we made a stable *yw*; UAS-*pdfr*-tandem; *tim*-Gal4 stock.

### *hEK-293* Cell Culture

*hEK-293* cells were maintained in DMEM, 10% FBS and 100U/mL penicillin and streptomycin in 5% CO_2_ atmosphere at 37°C. For all transient transfections, 1.5 x 10^6^ cells were used to inoculate T25 flasks, incubated overnight, then transfected with 10ug plasmid DNA and 20 uL lipofectamine 2000 reagent (Invitrogen Life Technologies). Five hours after transfection, cells were split 4 x 10^4^ cells/well into a 96-well assay plate. We created a series of stably-transfected *hEK-293* cells expressing WT and sequence-variant PDFRs, using the Flp-In System (Invitrogen Life Technologies, Waltham, Massachusetts) per manufacturers recommendations, and maintained them in DMEM supplemented with 150 μg/ml hygromycin B.

### cAMP Assays

We measured PDF Receptor signaling activity using a CRE-*Luciferase* reporter gene, following methods described by Johnson *et al.* [30]. The reporter gene construct was transiently transfected to each stable cell line and luminescence measured using Firefly Luciferase Assay Kit (Biotium, Inc., Fremont, California) and a Wallac 1420 VICTOR2 microplate reader (PerkinElmer, Inc., Waltham, Massachusetts). Concentration-effect curves, EC_50_, top values and p-values were calculated using the dose response, in a nonlinear regression using GraphPad Prism 8.0 software (San Diego, California).

### Locomotor activity Measures

All locomotor activity experiments were conducted with 2-5 days-old male flies at 25°C using Trikinetics Activity Monitors as previously described [9]. We crossed Gal4 lines to *w^1118^* to create control progeny. Locomotor activities were monitored for 6 days under different photoperiodic conditions, and then for 9 days under constant dark (DD). To analyze rhythmicity under constant conditions, we normalized the activity of flies from DD day 3 to day 9 and used χ2-periodogram analysis with a 95% confidence cut-off, as well as SNR analysis [67] Arrhythmic flies were defined by a power value ≤ 10 and width value ≤ 2, and period outside the range, 18 to 30 hours. To analyze periods, we used Graphpad Instat (v. 8) software to run one-way ANOVA measures followed by the Tukey-Kramer Multiple Comparisons Test. We used Clocklab (Actimetrics) software to produce actograms and the Brandeis Rhythms Package [73] to produce average activity plots (group eductions). To analyze onset and offset phases of Morning and Evening activity bouts on the final two days of light:dark entrainment (LD 5&6), we followed the method of Kula-Eversole *et al.* [56]: [(A_n+2_ + A_n+1_) – (A_n-1_ + A_n-2_) = ΔActivity]. Experimental genotypes were tested in the *han* mutant background*, pdfr^5304^* [11], or in the *pdf^01^* (null) mutant background [1], or in the *w^1118^* background, as noted.

### Presentation of behavioral data with expression of PDFR variants

Figures 2-4 all describe rhythmic locomotor behavior of adult *pdfr^han^* mutant flies that are expressing either a WT *pdfr* cDNA or the 1-7A variant. Figures 5 and 6 use a similar format but they introduce behavioral experiments performed in *pdfr^+^* or *pdf* mutant backgrounds, as described. All behavioral experiments extend across ∼6 days of light entrainment (LD), followed by activity during nine or more days of constant darkness (DD). For each genotype, we display locomotor activity during light entrainment in four ways: (i) group eductions (in panels marked **A, B and C**-**1** in these Figures), (ii) direct comparisons between two genotypes of the amplitudes of behavioral peaks using bin-by-bin analyses during the last two days of entrainment (days 5-6) (in panels marked **A, B and C-2-4**); (iii) derivations of Morning and Evening behavioral phases (in Panels **D and E**); and (iv) double-plotted group actograms (in Panels **F, G and H**). and The Bin-by-Bin analyses directly compare experimental genotypes to a control one [cf. 42]. They present data averaged across 2 to 5 independent experiments, to help identify the differences that most consistently correlate with genotype. Finally, Table 1 compiles measures of rhythmic activity displayed by the different genotypes in days 3-9 of constant darkness.

Additional details on Methods and procedures are found in **Supplemental Information**.

## ACKNOWLEDGEMENTS

We thank many colleagues for generously sharing reagents, fly strains and providing technical advice, all of which helped us pursue these studies. These include Xuemei Si, and Drs. Ali Salahpour, Dmitri Nusinow, Johanna Chiu, Xitong Liang, Paul Hardin, Zsolt Lenkei, Ken Blumer and David Wheeler. We thank the staff that manages the Flybase resource which provides valuable information, and the Danforth Plant Science Center Mass Spectroscopy Facility for expert assistance, and the Bloomington *Drosophila* Stock Center, and the Vienna *Drosophila* Research Center for providing many valuable fly stocks. We also thank Joel Levine and Orie Shafer for kindly providing comments on the manuscript.

## AUTHOR CONTRIBUTIONS

W.L., J.T. and P.H.T designed experiments. W.L. and J.T. performed experiments. W.L., J.T. and P.H.T analyzed experiments. P.H.T wrote the manuscript.

## LITERATURE CITED

1. Renn SC, Park JH, Rosbash M, Hall JC, Taghert PH. A pdf neuropeptide gene mutation and ablation of PDF neurons each cause severe abnormalities of behavioral circadian rhythms in Drosophila. Cell. 1999; 99:791–802. PMID: 10619432.

2. Peng Y, Stoleru D, Levine JD, Hall JC, Rosbash M. Drosophila free-running rhythms require intercellular communication. PLoS Biol. 2003; 1:E13. PMID: 12975658.

3. Lin Y, Stormo GD, Taghert PH. The neuropeptide pigment-dispersing factor coordinates pacemaker interactions in the Drosophila circadian system. J Neurosci. 2004; 24:7951–7. PMID: 15356209.

4. Vosko AM, Schroeder A, Loh DH, Colwell CS. Vasoactive intestinal peptide and the mammalian circadian system. Gen Comp Endocrinol. 2007; 152:165–75. PMID: 17572414.

5. Fernlund, P. Structure of a light-adapting hormone from the shrimp, Pandalus borealis. Biochim Biophysica Acta 1976; 439: 17–25. PMID: 95295.

6. Rao KR, Riehm JP, Zahnow CA, Kleinholz LH, Tarr GE, Johnson L, Norton S, Landau M, Semmes OJ, Sattelberg RM, Jorenby WH, Hintz MF. Characterization of a pigment-dispersing hormone in eyestalks of the fiddler crab Uca pugilator. Proc Nat’l Acad Sci (Wash) U S A. 1985; 82:5319–22. PMID: 16593589.

7. Cusumano P, Klarsfeld A, Chélot E, Picot M, Richier B, Rouyer F. PDF-modulated visual inputs and cryptochrome define diurnal behavior in Drosophila. Nat Neurosci. 2009; 12:1431–7. PMID: 19820704.

8. Zhang L, Lear BC, Seluzicki A, Allada R. The CRYPTOCHROME photoreceptor gates PDF neuropeptide signaling to set circadian network hierarchy in Drosophila. Curr Biol. 2009; 19:2050–5. PMID: 19913424.

9. Im SH, Li W, Taghert PH. PDFR and CRY signaling converge in a subset of clock neurons to modulate the amplitude and phase of circadian behavior in Drosophila. PLoS One. 2011; 6:e18974. PMID: 21559487.

10. Liang X, Holy TE, Taghert PH. Synchronous Drosophila circadian pacemakers display nonsynchronous Ca²⁺ rhythms. Science (Wash). 2016; 351:976–81. PMID: 26917772.

11. Hyun S, Lee Y, Hong ST, Bang S, Paik D, Kang J, Shin J, Lee J, Jeon K, Hwang S, Bae E, Kim J. Drosophila GPCR Han is a receptor for the circadian clock neuropeptide PDF. Neuron. 2005; 48:267–78. PMID: 16242407.

12. Lear BC, Merrill CE, Lin JM, Schroeder A, Zhang L, Allada R. A G protein-coupled receptor, groom-of-PDF, is required for PDF neuron action in circadian behavior. Neuron. 2005; 48:221–7. PMID: 16242403.

13. Mertens I, Vandingenen A, Johnson EC, Shafer OT, Li W, Trigg JS, De Loof A, Schoofs L, Taghert PH. PDF receptor signaling in Drosophila contributes to both circadian and geotactic behaviors. Neuron. 2005; 48:213–9. PMID: 16242402.

14. Shafer OT, Kim DJ, Dunbar-Yaffe R, Nikolaev VO, Lohse MJ, Taghert PH. Widespread receptivity to neuropeptide PDF throughout the neuronal circadian clock network of Drosophila revealed by real-time cyclic AMP imaging. Neuron. 2008; 58:223–37. PMID: 18439407.

15. Duvall LB, Taghert PH. The circadian neuropeptide PDF signals preferentially through a specific adenylate cyclase isoform AC3 in M pacemakers of Drosophila. PLoS Biol. 2012; 10:e1001337. PMID: 22679392.

16. Duvall LB, Taghert PH. E and M circadian pacemaker neurons use different PDF receptor signalosome components in Drosophila. J Biol Rhythms. 2013; 28:239–48. PMID: 23929551.

17. Seluzicki A, Flourakis M, Kula-Eversole E, Zhang L, Kilman V, Allada R. Dual PDF signaling pathways reset clocks via TIMELESS and acutely excite target neurons to control circadian behavior. PLoS Biol. 2014; 12(3):e1001810.. PMID: 24643294.

18. Depetris-Chauvin A, Fernández-Gamba A, Gorostiza EA, Herrero A, Castaño EM, Ceriani MF. Mmp1 processing of the PDF neuropeptide regulates circadian structural plasticity of pacemaker neurons. PLoS Genet. 2014; doi:10.1371/journal.pgen.1004700. PMID: 25356918.

19. Liang X, Holy TE, Taghert PH. A Series of Suppressive Signals within the Drosophila Circadian Neural Circuit Generates Sequential Daily Outputs. Neuron. 2017; 94:1173–1189. PMID: 28552314.

20. Liang X, Ho MCW, Zhang Y, Li Y, Wu MN, Holy TE, Taghert PH. Morning and Evening Circadian Pacemakers Independently Drive Premotor Centers via a Specific Dopamine Relay. Neuron. 2019; 102:843–857. PMID: 30981533.

21. Pincas H, González-Maeso J, Ruf-Zamojski F, Sealfon SC. G Protein -Coupled Receptors. In: Belfiore A, LeRoith D, editors. Principles of Endocrinology and Hormone Action.

22. Lefkowitz RJ, Shenoy SK. Transduction of receptor signals by β-Arrestins. Science (Wash). 2005; 308:512–7. PMID: 15845844.

23. Benovic JL, Kühn H, Weyand I, Codina J, Caron MG, Lefkowitz RJ. Functional desensitization of the isolated beta-adrenergic receptor by the beta-adrenergic receptor kinase: potential role of an analog of the retinal protein arrestin (48-kDa protein). Proc Nat’l Acad Sci (Wash) U S A. 1987; 84:8879–82. PMID: 2827157.

24. Lohse MJ, Benovic JL, Codina J, Caron MG, Lefkowitz RJ. β-Arrestin: a protein that regulates beta-adrenergic receptor function. Science (Wash). 1990; 248:1547–50. PMID: 2163110.

25. Ferguson SS, Downey WE 3rd, Colapietro AM, Barak LS, Ménard L, Caron MG. Role of β-arrestin in mediating agonist-promoted G protein-coupled receptor internalization. Science (Wash). 1996; 271:363–6. PMID: 8553074.

26. Smith JS, Rajagopal S. The β-Arrestins: Multifunctional Regulators of G Protein-coupled Receptors. J Biol Chem. 2016; 291:8969–77. PMID: 26984408.

27. Kelly E, Bailey CP, Henderson G. Agonist-selective mechanisms of GPCR desensitization. Brit J of Pharm. 2008; 153 Suppl 1: S379–88. PMID 18059321.

28. Abruzzi, K. C., Zadina, A., Luo, W., Wiyanto, E., Rahman, R., Guo, F., Shafer, O., & Rosbash, M. RNA-seq analysis of Drosophila clock and non-clock neurons reveals neuron-specific cycling and novel candidate neuropeptides. PLoS Genetics, 2017; 13(2), e1006613 doi.org/10.1371/journal.pgen.1006613

29. Klose M, Duvall L, Li W, Liang X, Ren C, Steinbach JH, Taghert PH. Functional PDF Signaling in the Drosophila Circadian Neural Circuit Is Gated by Ral A-Dependent Modulation. Neuron. 2016; 90:781–794. PMID: 27161526.

30. Johnson EC, Bohn LM, Barak LS, Birse RT, Nässel DR, Caron MG, Taghert PH. Identification of Drosophila neuropeptide receptors by G protein-coupled receptors-beta-arrestin2 interactions. J Biol Chem. 2003; 278(52):52172–8. PMID: 14555656.

31. Johnson EC, Bohn LM, Taghert PH. Drosophila CG8422 encodes a functional diuretic hormone receptor. J Exp Biol. 2004; 207:743–8. PMID: 14747406.

32. Johnson EC, Shafer OT, Trigg JS, Park J, Schooley DA, Dow JA, Taghert PH. A novel diuretic hormone receptor in Drosophila: evidence for conservation of CGRP signaling. J Exp Biol. 2005; 208:1239–46. PMID: 15781884.

33. Throckmorton, LH. The phylogeny, ecology and geography of Drosophila. In: King RC., editor. Handbook of Genetics. Vol 3. New York: Plenum; 1975. pp 421–469.

34. Mure LS, Hatori M, Zhu Q, Demas J, Kim IM, Nayak SK, Panda S. Melanopsin-Encoded Response Properties of Intrinsically Photosensitive Retinal Ganglion Cells. Neuron. 2016; 90:1016–27. PMID: 27181062.

35. Valdez-Lopez JC, Gulati S, Ortiz EA, Palczewski K, Robinson PR. Melanopsin Carboxy-terminus phosphorylation plasticity and bulk negative charge, not strict site specificity, achieves phototransduction deactivation. PLoS One. 2020; 15:e0228121. PMID: 32236094.

36. Langlet C, Langer I, Vertongen P, Gaspard N, Vanderwinden JM, Robberecht P. Contribution of the carboxyl terminus of the VPAC1 receptor to agonist-induced receptor phosphorylation, internalization, and recycling. J Biol Chem. 2005; 280: 28034 –28043. PMID: 15932876.

37. Im SH, Taghert PH PDF receptor expression reveals direct interactions between circadian oscillators in Drosophila. J Comp Neurol. 2010; 518:1925–45. PMID: 20394051.

38. Zhang Y, Emery P. GW182 controls Drosophila circadian behavior and PDF-receptor signaling. Neuron. 2013; Apr 10;78(1):152-65. PMID: 23583112.

39. Beerepoot P, Lam VM, Salahpour A. Measurement of G protein-coupled receptor surface expression. J Recept Signal Transduct Res. 2013; Jun;33(3):162-5. PMID: 23557016.

40. Zhou XE, He Y, de Waal PW, Gao X, Kang Y, Van Eps N, Yin Y, Pal K, Goswami D, White TA, Barty A, Latorraca NR, Chapman HN, Hubbell WL, Dror RO, Stevens RC, Cherezov V, Gurevich VV, Griffin PR, Ernst OP, Melcher K, Xu HE. Identification of Phosphorylation Codes for Arrestin Recruitment by G Protein-Coupled Receptors. Cell. 2017; 170:457–469.e13. PMID: 28753425.

41. Li S, Li S, Han Y, Tong C, Wang B, Chen Y, Jiang, J. (2016). Regulation of Smoothened Phosphorylation and High-Level Hedgehog Signaling Activity by a Plasma Membrane Associated Kinase. PLoS Biology, 2016; 14(6), e1002481. PMID: 27280464.

42. Murthy KS, Mahavadi S, Huang J, Zhou H, and Sriwai W. Phosphorylation of GRK2 by PKA augments GRK2-mediated phosphorylation, internalization, and desensitization of VPAC2 receptors in smooth muscle. Am J Physiol Cell Physiol. 2008; 294: C477–C487. PMID: 18077607.

43. Huang J, Mahavadi S, Sriwai W, Grider JR, Murthy KS. Cross-regulation of VPAC(2) receptor desensitization by M(3) receptors via PKC-mediated phosphorylation of RKIP and inhibition of GRK2. Am J Physiol Gastrointest Liver Physiol. 2007; 292:G867–74. PMID: 17170028.

44. Shetzline MA, Premont RT, Walker JK, Vigna SR, Caron MG. A role for receptor kinases in the regulation of class II G protein-coupled receptors. Phosphorylation and desensitization of the secretin receptor. J Biol Chem. 1998 Mar 20;273(12):6756–62. PMID: 9506976.

45. Walker JK, Premont RT, Barak LS, Caron MG, Shetzline MA. Properties of secretin receptor internalization differ from those of the 2-adrenergic receptor. J Biol Chem 1999; 274: 31515– 31523. PMID: 10531354.

46. Marie JC, Rouyer-Fessard C, Couvineau A, Nicole P, Devaud H, El Benna J, Laburthe M. Serine 447 in the carboxyl tail of human VPAC1 receptor is crucial for agonist-induced desensitization but not internalization of the receptor. Mol Pharmacol. 2003; 64: 1565–1574. PMID: 14645688.

47. Estall JL, Koehler JA, Yusta B, Drucker DJ. The glucagon-like peptide-2 receptor C terminus modulates beta-arrestin-2 association but is dispensable for ligand-induced desensitization, endocytosis, and G-protein-dependent effector activation. J Biol Chem. 2005; 280:22124–34. PMID: 15817468.

48. Krispel CM, Chen D, Melling N, Chen YJ, Martemyanov KA, Quillinan N, Arshavsky VY, Wensel TG, Chen CK, Burns ME. RGS expression rate-limits recovery of rod photoresponses. Neuron. 2006; 51:409–16. PMID: 16908407.

49. Berson DM, Dunn FA, Takao M. Phototransduction by retinal ganglion cells that set the circadian clock. Science (Wash). 2002; 295:1070–3. PMID: 11834835.

50. Emanuel AJ, Do MT. Melanopsin tristability for sustained and broadband phototransduction. Neuron. 2015; 85:1043–55. PMID: 25741728.

51. Somasundaram P, Wyrick GR, Fernandez DC, et al. C-terminal phosphorylation regulates the kinetics of a subset of melanopsin-mediated behaviors in mice. Proc Nat’l. Acad Sci (Wash) U S A. 2017; 114:2741–2746. PMID: 28223508.

52. Dephoure N, Gould KL, Gygi SP, Kellogg DR. Mapping and analysis of phosphorylation sites: a quick guide for cell biologists. Mol Biol Cell. 2013; 24:535–42. PMID: 23447708.

53. Chiu JC, Ko HW, Edery I. (2011) NEMO/NLK phosphorylates PERIOD to initiate a time-delay phosphorylation circuit that sets circadian clock speed. Cell. 2011; 145:357–70. PMID: 21514639.

54. Yu W, Houl JH, Hardin PE. NEMO kinase contributes to core period determination by slowing the pace of the Drosophila circadian oscillator. Curr Biol. 2011; 21:756–61. PMID: 21514156.

55. Helfrich-Förster C. The period clock gene is expressed in central nervous system neurons which also produce a neuropeptide that reveals the projections of circadian pacemaker cells within the brain of Drosophila melanogaster. Proc Nat’l Acad Sci (Wash) U S A. 1995; 92:612–6. PMID: 7831339.

56. Kula-Eversole E, Lee DH, Samba I, Yildirim E, Levine DC, Hong HK, Lear BC, Bass J, Rosbash M, Allada R. Phosphatase of Regenerating Liver-1 Selectively Times Circadian Behavior in Darkness via Function in PDF Neurons and Dephosphorylation of TIMELESS. Curr Biol. 2021; 31:138–149.e5. PMID: 33157022.

57. Grima B, Chélot E, Xia R, Rouyer F. Morning and evening peaks of activity rely on different clock neurons of the Drosophila brain. Nature. 2004; Oct 14;431(7010):869-73. PMID: 15483616.

58. Stoleru D, Peng Y, Agosto J, Rosbash M. Coupled oscillators control morning and evening locomotor behaviour of Drosophila. Nature. 2004; 431:862–8. PMID: 15483615.

59. . Rieger D, Shafer OT, Tomioka K, Helfrich-Förster C. Functional analysis of circadian pacemaker neurons in Drosophila melanogaster. J Neurosci. 2006; Mar 1;26(9):2531-43. PMID: 16510731.

60. Zhang Y, Liu Y, Bilodeau-Wentworth D, Hardin PE, Emery P. Light and temperature control the contribution of specific DN1 neurons to Drosophila circadian behavior. Curr Biol. 2010; Apr 13;20(7):600-5. PMID: 20362449.

61. Yao Z, Shafer OT. The Drosophila circadian clock is a variably coupled network of multiple peptidergic units. Science (Wash). 2014; 343(6178):1516-20. PMID: 24675961.

62. Schlichting M, Díaz MM, Xin J, Rosbash M. Neuron-specific knockouts indicate the importance of network communication to Drosophila rhythmicity. Elife. 2019 Oct 15;8:e48301. PMID: 31613223

63. 63. Delventhal R, O’Connor RM, Pantalia MM, Ulgherait M, Kim HX, Basturk MK, Canman JC, Shirasu-Hiza M. Dissection of central clock function in Drosophila through cell-specific CRISPR-mediated clock gene disruption. Elife. 2019;8:e48308. PMID: 31613218.

64. Menegazzi P, Beer K, Grebler V, Schlichting M, Schubert FK, Helfrich-Förster C. A Functional Clock Within the Main Morning and Evening Neurons of D. melanogaster Is Not Sufficient for Wild-Type Locomotor Activity Under Changing Day Length. Front Physiol. 2020; Mar 26;11:229. PMID: 32273848.

65. Schlichting M, Menegazzi P, Lelito KR, Yao Z, Buhl E, Dalla Benetta E, Bahle A, Denike J, Hodge JJ, Helfrich-Förster C, Shafer OT. A Neural Network Underlying Circadian Entrainment and Photoperiodic Adjustment of Sleep and Activity in Drosophila. J Neurosci. 2016; 36(35):9084–96. PMID: 27581451.

66. Vaze KM, Helfrich-Förster C. The Neuropeptide PDF Is Crucial for Delaying the Phase of Drosophila’s Evening Neurons Under Long Zeitgeber Periods. J Biol Rhythms. 2021;36(5):442–460. PMID: 34428956.

67. Blau J, Young MW. Cycling vrille expression is required for a functional Drosophila clock. Cell. 1999; Dec 10;99(6):661-71 PMID: 10612401.

68. Levine JD, Funes P, Dowse HB, Hall JC. Signal analysis of behavioral and molecular cycles. BMC Neurosci. 2002; 3:1. PMID: 1182533.

69. Wheeler DA, Hamblen-Coyle MJ, Dushay MS, Hall JC. Behavior in light-dark cycles of Drosophila mutants that are arrhythmic, blind, or both. J Biol Rhythms. 1993; 8:67–94. PMID: 8490212.

70. Shafer OT, Taghert PH. RNA-interference knockdown of Drosophila pigment dispersing factor in neuronal subsets: the anatomical basis of a neuropeptide’s circadian functions. PLoS One. 2009; 4:e8298. PMID: 20011537..

