## Supplemental Information for "Regulation of PDF receptor signaling controlling daily locomotor rhythms in *Drosophila*"

##### *PDFR Isoforms*

Flybase reports evidence for 4 distinct *pdfr* RNAs and 3 PDFR protein isoforms – (http://flybase.org/cgi-bin/gbrowse2/dmel/?Search=1;name=FBgn0260753): together these encode either of two N terminal sequence variants (RA vs RB) and either of two different C terminal sequence variants (RA and RB vs. RC and RD). The C terminal sequence encoded by isoforms RC and RD replaces the final 20 residues encoded by RA and RB with an alternative 50 residues, by splicing to a downstream exon (#10). Exon #10 is specifically deleted in a fly stock we previously described (P2-1, containing a unilateral deletion of the entire Exon 10, derived from *P{BG12523}*: [1]. That deletion did not affect locomotor rhythmicity adversely, nor did it produce any other locomotor phenotypes that resembled those of the *han* loss of function *pdfr* alleles, which include deletions of the C terminus-encoding sequences of all isoforms. Hence, we experimentally considered the two RNA isoforms RA and RB which display the shorter C terminal tail. The RA isoform is represented in the construct UAS-*pdfr*-16 which we have used in other published studies (e.g., [1-4]. Therefore, we focused our experimental design on the contributions of the sequence features of the C terminal tail of the PDFR-PA protein isoform to the strength and time course of PDF signaling that shapes locomotor rhythms.

##### *Genomic sequence analysis*

We used the PDFR sequence *D. melanogaster* NP_570007.2 as a blastp query (https://blast.ncbi.nlm.nih.gov/Blast.cgi?PAGE=Proteins) to search among non-redundant protein sequences (nr). Species representatives were not accepted if the C terminal sequence was incomplete or if only non-PDFR-A isoforms were retrieved. Accession numbers for the 16 additional *Drosophalid* species that we accepted are listed in Supplemental Table 1. Sequences were aligned using Clustal (https://www.ebi.ac.uk/Tools/msa/clustalo/) followed by minor manual adjustments.

***DNA Construction***

**Primers.** All primers used in creating DNA constructs are listed in Supplemental Table 2.

**β-lactamase Fusions.** The expression vector β-lactamase: β-AR2 pcDNA3.1 was provided by Dr. Ali Salahpour [5, 6] and used to make a β-lactamase pcDNA3.1 vector. The *CG13758* cDNA [1] was cloned into the β-lactamase pcDNA3.1 vector in frame with β-*lac* sequence, with restriction sites AscI and Not1. We also cloned the β *lac* sequence into a pcDNA5/frt vector via Nhe1 and HindIII. The *pdfr* wild type and C-terminal mutation series (see C-terminal construct oligos) was cloned into β-lac pcDNA5/frt vector with HindIII and Not1.

**FLAG::PDFR::EGFP fusions.** The expression vector Flag CB1R EGFP-N1 was provided by Dr. Zsolt Lenkei [7]. 4xFlag CG13758 EGFP-N1 vector was constructed by cloning 3xFlag from p3xFLAG-CMV-14 vector (Millipore-Sigma) in frame with the 1xFlag, with restriction site BglII on N-terminus and BamHI-BglII on C-terminus. *CG13758* WT sequence was cloned in frame with 4xFlag at N-terminus and EGFP on the C-terminus, using restriction sites BglII and Age1 with the CG13758 PCR fragment and BamHI and Age1 with the 4xFlag EGFP vector. Finally, site-directed mutagenesis was performed to mutate potential C-terminal phosphorylation sites (see C-terminal construct oligos). All constructs from 4xFlag 13758 EGFP were transferred to pcDNA5/frt vector (using HindIII and Not1) to make stable cell lines and to UAS-*attb* vector (using EcoRI and Not1) to create transgenic *Drosophila* lines.

**PDFR-Tandem Fusions.** The 2007bp *pdfr* coding sequence was amplified from a pcDNA3.1 *pdfr* construct [1] and the stop codon removed. The primers used were, *pdfr*-F 5'-GGAGATCTGCCACCATGACCCTCCTGTCGAACATTCTCG-3' and *pdfr*-R 5'-

GGCGACCGGTCCCTGCTCTGACAACTCAAATACAACTGACTC-3’. The product was blunt end-cloned into Eco-RV-pBluescript. The 451bp Tandem tag fragment was amplified from pENTR tandem plasmid (kind gift of Dmitri Nusinow, Donald Danforth Plant Science Center) to include a stop site at its N-terminus using primers (Integrated DNA Technologies, Coralville, Iowa) Tandem-foward 5’-GGGGAATTCGGAAGCTGGAGCCACCCTCAATTTGAAAAGGG-3’ and Tandem-reverse 5’-CCTCTAGATTACTATCACTTCTCGAACTGAGGATGACTCCAAGATCC-3’, then ligated in-frame to the *Pdfr* gene by digesting both with EcoRI and XbaI. Following sequenced verification, the PDFR-Tandem was transferred to the BlII/XbaI-digested UAS-attb vector.

***β-Lactamase (βlac)-fusion protein Assay.***

To quantify PDFR surface expression in cell lines, we employed the β-Lactamase method [5, 6]. $4\times10^4$ cells/well of *hEK-293* stably-expressing a β-lactamase-fusions to the C-termini of WT and sequence-variant PDFRs were split into a poly-lysine coated 96-well plate. Twenty-four hours after plating, the cells were incubated for 20 minutes at 37$^o$C, with and without PDF peptide (Phoenix Pharmaceuticals, Inc., Burlingame, California) diluted in MEM to $10^{-5}$M. The cells were washed with PBS and treated with nitrocefin (Cayman Chemical, Ann Arbor, Michigan) diluted to a final concentration of 100 µM in PBS. Immediately after adding the nitrocefin solution, the absorbance of each well was read at 490 nm every minute, for 30 min at 37$^o$C, with a Synergy H4 Hybrid Multi-Mode microplate reader (Bio Tek Instruments, Inc., Winooski, VT). Change in absorbance per minute readings were calculated into linear slopes and $R^2$ values. Basal levels of PDFR cell surface expression were obtained from cells not exposed to PDF. We divided the slope obtained from exposed cells exposed to peptide by the slope from cells not exposed to calculate % change in PDFR cell surface expression following 20 m exposure to PDF. Graphs and one-way ANOVA statistics were calculated using GraphPad Prism 8 software (San Diego, California).

***Tandem Affinity Purification.***

We grew *yw*; UAS-*pdfr*-tandem; *tim*-gal4 flies at 25°C on standard cornmeal medium. Flies were collected on ice in 25 ml aliquots and stored at -80°C. Fly heads were collected by sieving, cooled in liquid nitrogen, ground in a Retsch 400 mixer mill (4 x; 30 Hz; 45 sec), and re-suspended in 5 ml 0.3% IGEPAL lysis buffer (50mM Tris-HCl pH7.5, 125mM NaCl, 5% Gycerol, IGEPAL 0.3%**,** 1.5mM $MgCl_2$, 25mM NaF, 1mM $Na_3VO_4$, 0.05mM MG-115, 1mM PMSF, Protease inhibitor mix (Sigma P8340), Protease Inhibitor Cocktail (04693159001 Roche, Switzerland)) [8]. The head extract was sonicated (3-4 x; 10 sec), clarified by centrifugation at 4°C for 20 min at 50,000 x g and again for 40 minutes at 250,000 x g. The extract was then incubated with anti-flag M2 magnetic beads (M8823 Sigma, St. Louis, MO) on a rotator (1 hr at 4°C). The beads were washed twice in 0.3% IGEPAL lysis buffer, twice in Flag-to-HIS buffer (100 mM sodium phosphate, pH 8.0, 150 mM NaCl, 0.1% Triton X-100) and finally three times with Flag-to-HIS buffer without Triton X-100. The immunoprecipitated proteins were eluted from the anti-flag beads twice, at 4°C and then also at 30°C, using 500µg/ml 3x FLAG peptide (Sigma - F4799) prepared in flag-to-HIS buffer without Triton X-100. The combined eluates were incubated with His-tag Dynalbeads (10103D Invitrogen-ThermoFisher, Waltham, Ma, USA) for 0.5 h at RT with rotation. The His-tag beads were washed twice with flag-to-His buffer without triton X-100 and three times with 50mM ammonium bicarbonate buffer. After the last wash all the buffer was removed, and the beads were flash frozen in liquid nitrogen and stored at -80°C until submission for trypsin digest and mass spectrometry. Prior to submission, small samples from different purification steps were run on polyacrylamide gels (BioRad Laboratories, Inc., Hercules, Ca., USA) which were then rinsed in double distilled water and silver stained using a BioRad silver stain plus kit, according to manufacturer's instructions. Protein samples were reduced and alkylated using 10 mM TCEP in 50 mM ammonium bicarbonate for 1 h at 37 °C and 20 mM iodoacetamide for 30 min in the dark at RT, respectively. 1 µg of trypsin was added to the beads and samples were incubated at 37 °C overnight. The samples were then acidified and supernatants dried down in new tubes. The tryptic peptides were dissolved in 5% ACN/0.1% formic acid and 5 µL was injected to an LTQ-Orbitrap Velos Pro (ThermoFisher Scientific, MA) coupled with a U3000 RSLCnano HPLC (ThermoFisher Scientific). The liquid chromatography and mass spectrometer settings followed those previously described [9].

***Data Analysis*.**

Scaffold (Proteome Software Inc., Portland, OR; v.4.8.9) was used to validate MS/MS based peptide and protein identifications. The UniProt Database was used in determining peptide and protein identification which were accepted if they could be established at greater than 99.0% probability. The Scaffold Local FDR was used and only peptides probabilities with FDR 0.1% were used for further analysis. The Normalized Total Spectral was selected for the quantitative measurement estimate of the protein abundance of individual proteins in samples.

***SUPPLEMENTAL LITERATURE CITED***

1. Mertens I, Vandingenen A, Johnson EC, Shafer OT, Li W, Trigg JS, De Loof A, Schoofs L, Taghert PH. PDF receptor signaling in *Drosophila* contributes to both circadian and geotactic behaviors. Neuron. 2005; 48:213-9. PMID: 16242402.
2. Klose M, Duvall L, Li W, Liang X, Ren C, Steinbach JH, Taghert PH. Functional PDF Signaling in the *Drosophila* Circadian Neural Circuit Is Gated by Ral A-Dependent Modulation. Neuron. 2016; 90:781-794. PMID: 27161526.
3. Im SH, Taghert PH PDF receptor expression reveals direct interactions between circadian oscillators in *Drosophila.* J Comp Neurol. 2010; 518:1925-45. PMID**:** 20394051.
4. Liang X, Holy TE, Taghert PH. A Series of Suppressive Signals within the *Drosophila* Circadian Neural Circuit Generates Sequential Daily Outputs. Neuron. 2017; 94:1173-1189. PMID: 28552314.
5. Lam VM, Beerepoot P, Angers S, Salahpour A. A novel assay for measurement of membrane-protein surface expression using a β-lactamase. Traffic. 2013 14(7):778-84. PMID: 23574269.
6. Beerepoot P, Lam VM, Salahpour A. Measurement of G protein-coupled receptor surface expression. J Recept Signal Transduct Res. 2013; 33(3):162-5. PMID: 23557016.
7. Roland AB, Ricobaraza A, Carrel D, Jordan BM, Rico F, Simon A, Humbert-Claude M, Ferrier J, McFadden MH, Scheuring S, Lenkei Z. Cannabinoid-induced actomyosin contractility shapes neuronal morphology and growth. Elife. 2014; Sep 15;3:e03159. PMID: 25225054.
8. Tian X, Zhu M, Li L, Wu C. Identifying protein-protein interaction in Drosophila adult heads by Tandem Affinity Purification (TAP). J Vis Exp. 2013; 5;(82):50968. PMID: 24335807.

9. Huang H, Alvarez S, Bindbeutel R, Shen Z, Naldrett MJ, Evans BS, Briggs SP, Hicks LM, Kay SA, Nusinow DA. Identification of Evening Complex Associated Proteins in Arabidopsis by Affinity Purification and Mass Spectrometry. Mol Cell Proteomics. 2016; 15(1):201-17. PMID: 26545401;

***SUPPLEMENTAL TABLES***

**Supplemental Table 1. Accession numbers for PDFR-A from 17 *Drosophalid* species used to assess evolutionary conservation of individual AA residues in the C terminal regions.**

**Supplemental Table 2. Oligonucleotides used in this study for cloning**

**Supplemental Table 3. Potential phosphorylatable residues in the C terminal of the *D. melanogaster* PDFR-A isoform.**

**Supplemental Table 4. Phosphopeptides derived from PDFR-Tandem expression, detected *in vivo*.**

**Supplemental Table 5. Rhythmic behavior under Constant Dark Conditions for flies in which *GRK1, GRK2* and *-arrestin2* are manipulated.**

**Supplemental Table 1. *Drosophalid* PDFR Accession Numbers**

| Genus/ Species | Accession Number | Total AA length |
|---|---|---|
| *Drosophila melanogaster* | NP_570007.2 | 669 |
| *Drosophila takahashii* | XP_016994843.1 | 693 |
| *Drosophila ficusphila* | XP_017048318.1 | 679 |
| *Drosophila rhopaloa* | XP_016982487.1 | 671 |
| *Drosophila eugracilis* | XP_017067611.1 | 675 |
| *Drosophila erecta* | EDV45631.1 | 671 |
| *Drosophila simulans* | KMZ07909.1 | 717 |
| *Drosophila persimilis* | EDW26013.1 | 591 |
| *Drosophila miranda* | XP_033242917.1 | 685 |
| *Drosophila willistoni* | EDW82885.2 | 663 |
| *Drosophila bipectinate* | XP_017095130.1 | 684 |
| *Drosophila albomicans* | XP_034119693.1 | 702 |
| *Drosophila grimshawi* | EDW00227.1 | 679 |
| *Drosophila virilis* | XP_032288784.1 | 623 |
| *Drosophila hydei* | XP_030080101.1 | 668 |
| *Drosophila novamexicana* | XP_030566778.1 | 623 |
| *Drosophila navajoa* | XP_017964591.1 | 671 |

#### Supplemental Table 2.
#### Oligonucleotides used for cloning

**β-Lac 13758 pcDNA3 vector construction :** digest 13758 PCR & β-Lac GBR1 pcDNA3 with AscI & Not1 and ligate.

| | |
|---|---|
| c13758 AscI-F | CCCGGCGCGCCaccctcctgtcgaacattctcgactgcggaggc |
| c13758 Not1-R | CCCGCGGCCGCctactgctctgacaactcaaatacaactgactc |

**β-Lac pcDNA5 vector construction:** digest β-Lac PCR & pcDNA5 with Nhe1 & HindIII and ligate

| | |
|---|---|
| Blac atg Nhe-F | ggGCTAGCatggctagtgagacagacacactcctgcta |
| Blac end HindIII-R | ggAAGCTTccaatgcttaatcagtgaggcacctatctcag |

**β-Lac 13758 WT pcDNA5 vector construction:** Digest 13758 PCR & β-Lac pcDNA5 with HindIII & Not1 and ligate.

| | |
|---|---|
| c13758 HindIII-F | ggaagcttGCCACCATGACCCTCCTGTCGAACATTCTC |
| c13758 Not1-R | CCCGCGGCCGCctactgctctgacaactcaaatacaactgactc |

**β-Lac 13758 SNA pcDNA3 C-terminal mutagenesis:**

| | |
|---|---|
| c13758 AscI-F | CCCGGCGCGCCaccctcctgtcgaacattctcgactgcggaggc |
| c13758 S1A-R | gacctcgcacTGCcagctgggtggccagTGCctttagtag |
| c13758 S2A-R | cgtatccggcgccgtgttTGCagcaccTGCTGCcatTGCtgcccttttcggcgccca |
| c13758 S3A-R | tgcaggctgcactgcatcTGCatccggcgcTGCgttataagcacccgag |
| c13758 S4A-R | gcggtgatattcgctttccTGCggccgatggatctcctgc |
| c13758 S5A-R | cgcgctgcccgcctcctttTGCgtgaatgtgggagatggtc |
| c13758 S6A-R | caactgactcgggtggcacTGCTGCcgctgacggtactcttag |
| c13758 S7A-R | ggttttggtctactgctcTGCcaactcaaatacaacTGCctcgggtggcacggatgac |

**β-Lac S23A 13758 pcDNA3 C-terminal mutagenesis:**

Performed mutagenesis on β-Lac S2A 13758 pcDNA3 construct:

| | |
|---|---|
| c13758 AscI-F | CCCGGCGCGCCaccctcctgtcgaacattctcgactgcggaggc |
| c13758 S3A-R | tgcaggctgcactgcatcTGCatccggcgcTGCgttataagcacccgag |

**β-Lac S567A 13758 pcDNA3 C-terminal mutagenesis:**

Performed mutagenesis on β-Lac S5A 13758 pcDNA3 construct:

| | |
|---|---|
| c13758 AscI-F | CCCGGCGCGCCaccctcctgtcgaacattctcgactgcggaggc |
| c13758 S6A-R | caactgactcgggtggcacTGCTGCcgctgacggtactcttag |

followed by mutagenesis with B-Lac S56A 13758 pcDNA3 construct:

| | |
|---|---|
| c13758 AscI-F | CCCGGCGCGCCaccctcctgtcgaacattctcgactgcggaggc |
| c13758 S7A-R | ggttttggtctactgctcTGCcaactcaaatacaacTGCctcgggtggcacggatgac |

**β-Lac S1-4A 13758 pcDNA3 C-terminal mutagenesis:**

Performed mutagenesis on β-Lac S1A 13758 pcDNA3 construct:

| | |
|---|---|
| c13758 AscI-F | CCCGGCGCGCCaccctcctgtcgaacattctcgactgcggaggc |
| c13758 S2A-R | cgtatccggcgccgtgttTGCagcaccTGCTGCcatTGCtgcccttttcggcgccca |

followed by mutagenesis with β-Lac S12A 13758 pcDNA3 construct:

| | |
|---|---|
| c13758 AscI-F | CCCGGCGCGCCaccctcctgtcgaacattctcgactgcggaggc |
| c13758 S3A-R | tgcaggctgcactgcatcTGCatccggcgcTGCgttataagcacccgag |

followed by mutagenesis with β-Lac S123A 13758 pcDNA3 construct:

| | |
|---|---|
| c13758 AscI-F | CCCGGCGCGCCCaccctcctgtcgaacattctcgactgcggaggc |
| c13758 S4A-R | gcggtgatattcgctttccTGCggccgatggatctcctgc |

**β-Lac S1-7A 13758 pcDNA3 C-terminal mutagenesis:**

Performed mutagenesis on β-Lac S1-4A 13758 pcDNA3 construct:

| | |
|---|---|
| c13758 AscI-F | CCCGGCGCGCCCaccctcctgtcgaacattctcgactgcggaggc |
| c13758 S5A-R | cgcgctgcccgcctcctttTGCgtgaatgtgggagatggtc |

followed by mutagenesis with β-Lac S1-5A 13758 pcDNA3 construct:

| | |
|---|---|
| c13758 AscI-F | CCCGGCGCGCCCaccctcctgtcgaacattctcgactgcggaggc |

**Supplemental Table 3. Potential phosphorylatable residues in the C terminal of the PDFR-A isoform**

| Residue | conservation among 17 species | Cluster designation | phosphorylated *in vivo?* |
|---|---|---|---|
| S512 | 100% | CL1 | |
| T514 | 53% | | |
| S518 | 100% | CL1 | |
| S531 | 100% | CL2 | YES |
| Y533 | 100% | CL2 | |
| S534 | 100% | CL2 | YES |
| Y537 | 100% | CL2 | |
| T539 | 100% | CL3 | |
| T543 | 100% | CL3 | |
| S554 | 35% | | |
| T556 | 100% | CL4 | |
| S560 | 94% | | YES |
| S572 | 94% | | |
| S573 | 94% | | |
| S574 | 88% | | |
| T607 | 59% | | |
| T616 | 6% | | |
| S618 | 18% | | |
| S622 | 88% | CL5 | |
| S627 | 18% | | |
| S630 | 88% | | |
| T632 | 47% | | |
| S635 | 76% | | |
| S653 | 59% | | YES |
| S655 | 100% | CL6 | |
| S656 | 100% | CL6 | |
| S661 | 100% | CL7 | |
| S667 | 100% | CL7 | |

**Supplemental Table 4. Phosphopeptides derived from PDFR-Tandem, detected *in vivo***

Spectra are shown in Supplemental Figure 2

| Experiment | collection time | phosphorylated residue | Tryptic Peptide | CL # |
|---|---|---|---|---|
| 2 | evening | S531 | (R)AS*MYSGAYNTAPDTAVQPAGDPSATGK | CL2 |
| 3 | evening | S531 | (R)AS*MYSGAYNTAPDTAVQPAGDPSATGK | CL2 |
| | | T543 | (R)ASMYSGAYNTAPDT*AVQPAGDPSATGK | CL3 |
| | | S560 | (K)RIS*PPNKR | --- |
| 5 | evening | S531 | (R)AS*MYSGAYNTAPDTAVQPAGDPSATGK | CL2 |
| | | S534 | (R)ASMYS*GAYNTAPDTAVQPAGDPSATGK | CL2 |
| 7 | morning | S653 | (R)VPS*ASSVPPESVVFELSEQGPVAIEFGSWSHPQFEK | --- |
| 8 | evening | S531 | (R)AS*MYSGAYNTAPDTAVQPAGDPSATGK | CL2 |
| | | S560 | (K)RIS*PPNKR | --- |

**Supplemental Table 5. Rhythmic behavior under Constant Dark Conditions for flies in which *GRK1*, *GRK2* and *KRZ* (β-arrestin2) are manipulated**

| Genotype | Fly # | AR% | TAU | PWR | WI | SNR | T-test |
|---|---|---|---|---|---|---|---|
| timg > + | 16 | 0% | 24.28 | 157.71 | 6.13 | 1.863 | |
| timg > gprk1 | 15 | 20% | 24.42 | 49.17 | 4.17 | 0.541 | ns |
| timg > gprk2.2 | 15 | 47% | 24.56 | 53.83 | 4.00 | 0.616 | ns |
| dcr2; timg > + | 16 | 0% | 24.34 | 193.14 | 6.19 | 2.784 | |
| dcr2; timg > gprk1 RNAi | 15 | 53% | 23.86 | 62.70 | 4.43 | 0.583 | ns |
| dcr2; timg > gprk2 RNAi | 15 | 13% | 24.38 | 73.25 | 3.92 | 0.864 | ns |
| dcr2; tim >+ | 11 | 0% | 24.05 | 123.30 | 7.18 | 2.69 | |
| dcr2; timg > KK10463 gprk2 RNA | 15 | 7% | 23.57 | 64.76 | 4.50 | 0.84 | *** |
| dcr2; timg > KK103756 krz RNAi | 14 | 100% | --- | --- | --- | --- | --- |

tau values compared to controls with Student's t-Test:

ns - not different;

*** - different, p < 0.01

*SUPPLEMENTAL FIGURES LEGENDS*

**Supplemental Figure 1 [Supporting Figure 1]. Alignment of the C terminal PDFR-A sequences from 17 different *Drosophalid* species.** The predicted 7$^{th}$ transmembrane domain (TM7) is marked in GREY. The C terminal tail starts with V505 (numbering for the *melanogaster* protein). The 28 potentially phosphorylated residues in the D.m. C terminal tail are highlighted in color: the 14 residues chosen for analysis are marked by their Cluster (CL) designation (1 to 7) and marked in AQUA; the 14 non-selected residues are marked in YELLOW. See Supplemental Table 2 for additional sequence information.

**Supplemental Figure 2 [Supporting Figure 2]. Locomotor Rhythms exhibited by WT PDFR and by other PDFR Variants under Short-day (winter-like) condition.** All behavioral records were recorded from *han* (*pdfr* mutant) flies that expressed either no UAS transgene (A), or a UAS-WT *pdfr* transgene (B) or a variety of Simple *pdfr* Variants, including 2-3A (C), 4A (D), 5A (E), 6A (F) or 7A (G), or Multiple *pdfr* variants, including 5-7A (H), 1-4A (I), 1-5A (J), and 1-6A (K). <u>Top Right Box:</u> A schematic of the PDFR C terminal segment for the WT and all variants studied: see Figure Legend 1 for details. Letters to the right of each variant C terminal segment correspond to the Panels in this Figure that display the behavior observed following its expression. Each Panel (A)-(K) contains sub-panels (1) and (2): Sub-Panel (1) displays a daily plot of locomotor activity (a group eduction) averaged over the last two days of entrainment (LD 5-6). Open bars indicate the 8 hr periods of Lights-on and filled bars indicate 16 hr periods of Lights-off. Sub-panel (2) displays a double-plotted group actogram throughout the 6 days of LD entrainment, followed by ~9 days of (DD, grey background). Green lines indicate the phase of the dominant activity period in DD. Panel L displays the average Morning activity Phase Onset timepoint, and Panel M displays the average Evening activity Phase Offset (marked by a Blue Arrow) for each genotype over the last two days of entrainment (LD 5-6). The positions of the Red and Blue arrows in panels A-K are representative phase points; panels L and M present their true values respectively. Analyses represent ANOVA followed by Dunnett's post hoc multiple comparisons of all compared to WT: ns = not significant; * = $p<0.05$; ** = $p<0.01$; *** = $p<0.005$; **** = $p<0.001$.

**Supplemental Figure 3 [Supporting Figure 2]. Amplitude measures of locomotor rhythms exhibited by WT PDFR and by other PDFR Variants under Short-day (winter-like) conditions.** All behavioral records were recorded from *han* (*pdfr* mutant) flies that expressed either no UAS transgene (A – marked in YELLOW), or a WT *pdfr* cdNA (B – marked in BLACK) or a variety of Simple *pdfr* Variants (all marked in RED) including 2-3A (C), 4A (D), 5A (E), 6A (F), 7A (G), 5-7A (H), 1-4A (I), 1-5A (J), and 1-6A (K). These measures were averaged across all experiments run for each individual genotype (see Table 1 for N and n values). Top Right Box: A schematic of the PDFR C terminal segment for the WT and all variants studied: see Figure Legend 1 for details. Letters to the right of each variant C terminal segment correspond to the Panels in this Figure that display the behavior observed following its expression. Each Panel (A)-(L) contains sub-panels (1) through (3), each of which displays Bin-by-Bin analyses of activity levels sorted by 30 min bins, for three different time periods: (1) ZT17-23.5; (2) ZT 0.5 – 8; (3) ZT 8.5-16. The missing bin at timepoint 0 contains the startle response that accompanies the sudden lights-on signal. Blue asterisks indicate significantly-different activity levels according to a Student's T-test following an ANOVA ($p< 0.05$). Red arrows highlight elevated Morning activity levels displayed by different PDFR variants. Blue arrows highlight elevated Evening activity levels produced by different PDFR variants.

**Supplemental Figure 4 [Supporting Figure 3]. Locomotor Rhythms exhibited by WT PDFR and by other PDFR Variants under Equinox (12:12) condition.** All behavioral records were recorded from *han* (*pdfr* mutant) flies that expressed either no UAS transgene (A), or a UAS-WT *pdfr* transgene (B) or a variety of

Simple *pdfr* Variants, including 2-3A (C), 4A (D), 5A (E), 6A (F) or 7A (G), or Multiple *pdfr* variants, including 5-7A (H), 1-4A (I), 1-5A (J), and 1-6A (K). Top Right Box: A schematic of the PDFR C terminal segment for the WT and all variants studied: see Figure Legend 1 for details. Letters to the right of each variant C terminal segment correspond to the Panels in this Figure that display the behavior observed following its expression. Each Panel (A)-(L) contains sub-panels (1) and (2): Sub-Panel (1) displays a daily plot of locomotor activity (a group eduction) averaged over the last two days of entrainment (LD 5-6). Open bars indicate the 12 hr periods of Lights-on and filled bars indicate 12 hr periods of Lights-off. Sub-panel (2) displays a double-plotted group actogram throughout the 6 days of LD entrainment, followed by ~9 days of (DD, grey background). Green lines indicate the phase of the dominant activity period in DD. Panel L displays the average Morning activity Phase Onset timepoint, and Panel M displays the average Evening activity Phase Onset (marked by a Blue Arrow) for each genotype over the last two days of entrainment (LD 5-6). The positions of the Red and Blue arrows in panels A-K are representative phase points; panels L and M present their true values respectively. Analyses represent ANOVA followed by Dunnett's post hoc multiple comparisons of all compared to WT: ns = not significant; * = $p<0.05$; ** = $p<0.01$; *** = $p<0.005$; **** = $p<0.001$.

**Supplemental Figure 5 [Supporting Figure 3]. Amplitude measures of locomotor rhythms exhibited by WT PDFR and by other PDFR Variants under Equinox (12:12) conditions.** All behavioral records were recorded from *han* (*pdfr* mutant) flies that expressed either no UAS transgene (A – marked in YELLOW), or a WT *pdfr* cdNA (B – marked in BLACK) or a variety of Simple *pdfr* Variants (all marked in RED) including 2-3A (C), 4A (D), 5A (E), 6A (F), 7A (G), 5-7A (H), 1-4A (I), 1-5A (J), and 1-6A (K). These measures were averaged across all experiments run for each individual genotype (see Table 1 for N and n values). Top Right Box: A schematic of the PDFR C terminal segment for the WT and all variants studied: see Figure Legend 1 for details. Letters to the right of each variant C terminal segment correspond to the

Panels in this Figure that display the behavior observed following its expression. Each Panel (A)-(L) contains sub-panels (1) through (3), each of which displays Bin-by-Bin analyses of activity levels sorted by 30 min bins, for three different time periods: (1) ZT17-23.5; (2) ZT 0.5 – 8; (3) ZT 8.5-16. The missing bin at timepoint 0 contains the startle response that accompanies the sudden lights-on signal. Blue asterisks indicate significantly-different activity levels according to a Student's T-test following an ANOVA ($p < 0.05$). Red arrows highlight elevated Morning activity levels displayed by different PDFR variants. Blue arrows highlight elevated Evening activity levels produced by different PDFR variants.

**Supplemental Figure 6 [Supporting Figure 4]. Locomotor Rhythms exhibited by WT PDFR and by other PDFR Variants under Long-day (summer-like) condition.** All behavioral records were recorded from *han* (*pdfr* mutant) flies that expressed either no UAS transgene (A), or a UAS-WT *pdfr* transgene (B) or a variety of Simple *pdfr* Variants, including 2-3A (C), 4A (D), 5A (E), 6A (F) or 7A (G), or Multiple *pdfr* variants, including 5-7A (H), 1-4A (I), 1-5A (J), and 1-6A (K). <u>Top Right Box:</u> A schematic of the PDFR C terminal segment for the WT and all variants studied: see Figure Legend 1 for details. Letters to the right of each variant C terminal segment correspond to the Panels in this Figure that display the behavior observed following its expression. Each Panel (A)-(L) contains sub-panels (1) and (2): Sub-Panel (1) displays a daily plot of locomotor activity (a group eduction) averaged over the last two days of entrainment (LD 5-6). Open bars indicate the 16 hr periods of Lights-on and filled bars indicate 8 hr periods of Lights-off. Sub-panel (2) displays a double-plotted group actogram throughout the 6 days of LD entrainment, followed by ~9 days of (DD, grey background). Green lines indicate the phase of the dominant activity period in DD. Panel L displays the average Morning activity Phase Offset timepoint, and Panel M displays the average Evening activity Phase Onset (marked by a Blue Arrow) for each genotype over the last two days of entrainment (LD 5-6). The positions of the Red and Blue arrow in panels A-K are representative phase points; panels L and M present their true values respectively.

Analyses represent ANOVA followed by Dunnett's post hoc multiple comparisons of all compared to WT: ns = not significant; * = p<0.05; ** = p<0.01; *** = p<0.005; **** = p<0.001.

**Supplemental Figure 7 [Supporting Figure 4]. Amplitude measures of locomotor rhythms exhibited by WT PDFR and by other PDFR Variants under Long-day (summer-like) conditions.** Behavioral records recorded from *han* (*pdfr* mutant) flies that expressed either no UAS transgene (A – marked in YELLOW), or a WT *pdfr* cdNA (B – marked in BLACK) or a variety of Simple *pdfr* Variants (all marked in RED) including 2-3A (C), 4A (D), 5A (E), 6A (F), 7A (G), 5-7A (H), 1-4A (I), 1-5A (J), and 1-6A (K). These measures were averaged across all experiments run for each individual genotype (see Table 1 for N and n values). Top Right Box: A schematic of the PDFR C terminal segment for the WT and all variants studied: see Figure Legend 1 for details. Letters to the right of each variant C terminal segment correspond to the Panels in this Figure that display the behavior observed following its expression. Each Panel (A)-(L) contains sub-panels (1) through (3), each of which displays Bin-by-Bin analyses of activity levels sorted by 30 min bins, for three different time periods: (1) ZT17-23.5; (2) ZT 0.5 – 8; (3) ZT 8.5-16. The missing bin at timepoint 0 contains the startle response that accompanies the sudden lights-on signal. Blue asterisks indicate significantly-different activity levels according to a Student's T-test following an ANOVA (p< 0.05). Red arrows highlight elevated Morning activity levels displayed by different PDFR variants. Blue arrows highlight elevated Evening activity levels produced by different PDFR variants.

Supplemental Figure 8. Average daily locomotor rhythms in flies expressing WT and 1-7A variant PDFR transgenes that lack GFP fusions. All behavioral records were recorded from *han* (*pdfr* mutant) flies that expressed either no UAS transgene (panels (A, D, I, and L) - yellow), or a WT *pdfr* cdNA (panels (B, E, J and M) - black) or the PDFR 1-7A Multiple Variant (panels (C, F, K and N) – red). Panels (A-F) present data recorded under Short-Day (winter-like) conditions; panels (I-N) present data recorded under Long Day (summer-like) conditions. Each Panel (A - C) and (I -K) contains four sub-panels: Sub-Panel (1) displays an average daily plot of locomotor activity averaged over the final two days of light entrainment (a group eduction): Open bars indicate the periods of Lights-on and filled bars indicate periods of Lights-off. Sub-Panels 2-4 display Bin-by-Bin analyses of activity levels sorted by 30 min bins, for three different time periods: (1) ZT17-23.5; (2) ZT 0.5 – 8; (3) ZT 8.5-16. The missing bin at timepoint 0 contains the startle response that accompanies the sudden lights-on signal. Blue asterisks indicate significantly-different activity levels according to a Student's T-test following an ANOVA ($p< 0.05$). Panels (D - F) and (L - N) display double-plotted group actograms throughout the 6 days of Light: dark entrainment, followed by 9 days of constant darkness (DD, grey background). Panels (G) and (H) display the average Phase Onsets and Offsets (respectively) for the Morning and Evening activity periods for each genotype over the last two days of entrainment under short days (LD 5-6). Blue arrows highlight the elevated Evening activity amplitudes. Panels (O) and (P) display the average Phase Offsets and Onsets (respectively) for the Morning and Evening activity periods for each genotype over the last two days of entrainment under long days (LD 5-6). Red Arrows highlight the elevated amplitude of the Morning peak. Ns – not significant; * - $p < 0.05$; *** - $p < 0.005$; **** - $p < 0.001$

**Supplemental Figure 9. Basal cAMP signaling displayed by the PDFR variant series following functional expression *in vitro*.** Luciferase measurements in *hEK-293T* cells stably expressing WT PDFR or its variants and transiently expressing *CRE-Luciferase*. The histogram represents basal levels of 2$^{nd}$ messenger signaling, i.e., in the absence of stimulation by neuropeptide PDF. Values represent the mean +/-SEM of three independent measurements, and were analyzed by Student 's T-test: * = $p < 0.05$; ns = not significantly different.

**Supplemental Figure 10. EC50 values for cAMP generation displayed by the PDFR variant series following functional expression *in vitro*.** Luciferase measurements in *hEK-293T* cells stably expressing WT PDFR or its variants and transiently expressing *CRE-Luciferase*. The histograms represent EC50 values for PDF-stimulated 2nd messenger signaling. Values represent the mean +/-SEM of three independent measurements, and were analyzed by Student 's T-test: * = $p < 0.05$; ns = not significantly different.

**Supplemental Figure 11. Top Values for cAMP generation displayed by the PDFR variant series following functional expression *in vitro*.** Luciferase measurements in *hEK-293T* cells stably expressing WT PDFR or its variants and transiently expressing *CRE-Luciferase*. The histograms represent the top values achieved for 2nd messenger signaling following PDF-stimulation. Values represent the mean +/-SEM of three independent measurements, and were analyzed by Student 's T-test: * = $p < 0.05$; ns = not significantly different.

**Supplemental Figure 12**. **Surface expression of the PDFR variant series following functional expression *in vitro***. β-Lactamase activity measurements in *hEK-293T* cells stably expressing WT PDFR or its variants fused to β-lactamase at the N terminus. The histograms represent the basal values for surface receptor expression in the absence of stimulation by neuropeptide PDF. Values represent the mean +/-SEM of three independent measurements, and were analyzed by Student 's T-test: * = $p < 0.05$; ns = not significantly different.

**Supplemental Figure 13. Percentage change in surface expression of the PDFR variant series after exposure to PDF, following functional expression *in vitro*.** β-Lactamase activity measurements in *hEK-293T* cells stably expressing WT PDFR or its variants fused to β-lactamase at the N terminus. The histograms represent the values for surface receptor expression 20 m after exposure to neuropeptide PDF. Values represent the mean +/-SEM of three independent measurements, and were analyzed by Student 's T-test: * = $p < 0.05$; ns = not significantly different.

**Supplemental Figure 14. *In vivo* detection of phosphopeptides from the PDFR C terminal tail.** Nine spectra of phosphopeptides derived from a UAS-*PDFR*-Tandem construct from eight independent immunoprecipitation experiments.

**Supplemental Figure 15. Average daily locomotor rhythms in flies with manipulations of *GRK1, GRK2* and *⍰-arrestin2*.** Group eductions of locomotor activity profiles for different genotypes averaged over six days of light entrainment. Open bars indicate periods of Lights-On and filled bars indicate periods of Lights-Off. The column present manipulations for each of four different gene targets (none; *Gprk1*; *Gprk2* and ⍰*-arr2* ***(kurtz))*** using *tim*(UAS)-Gal4. The Upper panels (D and F) present over-expression experiments. The Middle (E and G) and Lower panels (H and I) present RNAi experiments. UAS-RNAi constructs illustrated in panels E and G were created and shared by the Paul Hardin laboratory. Values for N and n are found in Supplemental Table 5.

### Supplemental Figure 1.

TM7 — Transmembrane Segment 7 (grey)
Conserved Ser/Thr/Tyr; Modified in Experiments (cyan)
Conserved Ser/Thr/Tyr; Not Modified in Experiments (yellow)

```
                    -----TM7--------     CL1   CL1        CL2-  -CL3
melanogaster  FAVWSYGTHFLTSFQGFFIALIYCFLNGEVRAVLLKSLATQLSVRGHPEWAPKRASMYSGAYNTAPDTDAV--QPAGD---   552
takahashi     FAVWSYVTHFLTSFQGFFIALIYCFLNGEVRAVLLKSLATQLSVRGHPEWAPKRASMYSGAYNTAPDTDAV--QPAGD---   619
ficusphila    FAVWSYGTHFLTSFQGFFIALIYCFLNGEVRAVLLKSLATQLSVRGHPEWAPKRASMYSGAYNTAPDTDAV--HPAGD---   608
rhopaloa      FAVWSYGTHFLTSFQGFFIALIYCFLNGEVRAVLLKSLATQLSVRGHPEWAPKRASMYSGAYNTAPDTDAV--QPAGD---   599
eugracilis    FAVWSYGTHFLTSFQGFFIALIYCFLNGEVRAVLLKSLATQLSVRGHPEWAPKRASMYSGAYNTAPDTDAV--QPAGD---   605
erecta        FAVWSYGTHFLTSFQGFFIALIYCFLNGEVRAVLLKSLATQLSVRGHPEWAPKRASMYSGAYNTAPDTDAV--QPAGD---   602
simulans      FAVWSYGTHFLTSFQGFFIALIYCFLNGEVRAVLLKSLATQLSVRGHPEWAPKRASMYSGAYNTAPDTDAV--QPAGD---   643
persimilis    FAVWSYGTHFLTSFQGFFIALIYCFLNGEVRACLLKSLSNQLSLRGHPEWAPKRPSMYSGAYNTAPDTDAVGQQPVAETQ-   525
miranda       FAVWSYGTHFLTSFQGFFIALIYCFLNGEVRACLLKSLSNQLSLRGHPEWAPKRPSMYSGAYNTAPDTDAVGQQPVAETQ-   619
willistoni    FAVWSYGTHFLTSFQGFFIALIYCFLNGEVRAVLLKSLATQMSVRGHPEWVPKRASMYSGAYNTAPDTDAVQTQPQGGPDI   605
bipectinata   FAVWSYGTHFLTSFQGFFIALIYCFLNGEVRAVLLKSLATQMSVRGHPEWVPKRASMYSGAYNTAPDTDAVIQQP-GD---   626
albomicans    FAVWSYVTHFLTSFQGFFIALIYCFLNGEVRTVLLKSLAVWMSVRGHPEWVPKRASMYSGAYNTAPDTDAL-QQP-GDAPP   641
grimshawi     FAVWSYVTHFLTSFQGFFIALIYCFLNGEVRTVLLKSLAVWMSVRGHPEWAPKRASMYSGAYNTAPDTDVV-QQP-GE---   623
virilis       FAIWSYVTHFLTSFQGFFIALIYCFLNGEVRAVMLKSIAVWLSVRGHPEWAPKRPSMYSGAYNTAPDTDPQLKQ--GD---   473
hydei         FAIWSYVTHFLTSFQGFFIALIYCFLNGEVRAVLLKSLAVWMSVRGHPEWVPKRASMYSAAYNTAPDTEAPAQQT-VD---   601
novamexicana  FAIWSYVTHFLTSFQGFFIALIYCFLNGEVRAVMLKSIAVWLSVRGHPEWAPKRPSMYSGAYNTAPDTDPQLKP--GD---   581
navajoa       FAIWSYVTHFLTSFQGFFIALIYCFLNGEVRAVLLKSLAVWMSVRGHPEWIPKRASMYSAAYNTAPDTEPPAQQS-VE---   602

                        CL4
melanogaster  PSATG---KRISPPNKRLNGRKPSSASIVMIHEPQQRQRLMRLMPRLQNKAREKGKD--RVEK----TDAEA---------   614
takahashi     PSAT----KRISPPHKRLNGRKPSSASIVMIHEPQQRQRLIRLIPRLQNKAREKSRD--RVDKADAETDPQA---------   633
ficusphila    PSAPG---KRISPPNKRLNGRKPSSASIVMIHEPQQRHRLIRLIPRLQNKNQEKGKD--RVEKADIDTEL-----------   623
rhopaloa      PLATG---KRISPPNKRLNGRKPSSASIVMIHEPQQRQRLIRLIPRLQNKAREKGRD--RVEKADAETEPEP---------   612
eugracilis    PSATG---KRISPPNKRLNGRKPSSASIVMIHEPQQRQRLIRLIPRLQNKAREKSKD--RVEKAEPETEP-----------   620
erecta        PSATG---KRISPPNKRLNGRKPSSASIVMIHEPQQRQRLMRLMPRLQNKAREKGKE--RVEK----TDKEA---------   616
simulans      PSATG---KRISPPNKRLNGRKPSSASIVMIHEPQQRQRLMRLMPRMQNKAREKGKD--RVEK----TDAEA---------   660
persimilis    -AATGG--LRTSPINRR-QN---TSASIVMIHEPNHRQRLVRLVQRNHNNNNQESPRRQRGER----TDDEDRIGIAAGAG   532
miranda       -AATGG--LRTSPINRR-QN---TSASIVMIHEPNHRQRLVRLVQRNHNNNNQESPRRQRGER----TDDEDRIGIAAGAG   626
willistoni    LPSTG---KRISPPNKRHTSRKASSASIVIIHEPQQRQRLLRLLQRSSTMNISNAKNLTNQT-----TDIDPEAPGVSPTA   608
bipectinata   NPASG---KRISPPNKRLNGRKASSASIVMIHEPQQRHRLLRLLPRLHSQRKDRDKIKDRDKDNDRDRQRDDRQLE-----   629
albomicans    PATTTAAGKRVSPPNKRLNCRKASSVSIVIAKEPQRQRLVQLVQQQQQQQRNNNN-SRNIMPDDE----------ASAS-   653
grimshawi     AISTS---RRVPPPIKRLKSRKANNVTIVISNEPQQQQQQHQMLQQRNTNNNSTNGSSPRTV--EDDTGS------AMAT-   634
virilis       PQQSG---KRLSQSTKRSNSRKASSVTIVISTEPQIHRYVPYVPRRNNNNRASTGSA--RVRGILKATEEP-----ASGSA   573
hydei         ALSSS---LRISPQNRRLNSRKASSVIIVIANEPQRQRFVQFVQQRNNNNTANSN----------QGKDEP-----ASGSA   619
novamexicana  PQQSG---KRLSQSTKRLNSRKASSVTIVISTEPQIHRYIPYIPRRNNNNRASTDSA--RVRGILKATEEP-----ASGSA   571
navajoa       ALSSN---IRISPQSRRLNCRKASSVIIVIANEPQRQRSVQSVQQRNNNNTANGNGN--------QGKDEP-----ASGSA   618

                        CL5                                            CL6  CL7  CL7
melanogaster  --EPDPTISHIHSKEAG------SARSR-TRGSKWIMG-ICFRGQKVLRVPSA-SSVPPESVVFELSEQ   669
takahashi     ----DPAISRIHSKESSGGGGG-STGSR-NRGSKWIMG-ICFRGQKVLRVPSA-SSVPPESVVFELSEQ   693
ficusphila    ----DPAITRIQSKEST------GSTSR-SRGSKWIMG-ICFRGQKVLRVPSA-SSVPPESVVFELSEQ   679
rhopaloa      ----DPAITRIHSKEAGSTG---GTASRNSRGSKWIMG-ICFRGQKVLRVPSA-SSVPPESVVFELSEQ   671
eugracilis    ----DPAISRIHSKETG------VGGSR-TRGSKWIMG-ICFRGQKVLRVPSA-SSVPPESVVFELSEQ   675
erecta        EPEPDPAISRIHSKEAD------RARSR-TRGSKWIMG-ICFRGQKVLRVPSA-SSVPPESVVFELSEQ   671
simulans      ----EPAIARIHSKEAG------SARSR-TRGSKWIMG-ICFRGQKVLRVPSA-SSVPPESVVFELSEQ   715
persimilis    -VEAEVVVV-QDSVGGAVG---RKRETRIHAGTKWMISGLCFRGQKVLRVPST-SSVPPESVVFELSEQ   591
miranda       -VEAEVVVV-QDSVGGAVG---RKRETRIHAGTKWMISGLCFRGQKVLRVPST-SSVPPESVVFELSEQ   685
willistoni    MMDANAATQRIHSKESS-------ARTNST--PNWMIFVLCFRGQKVLRVPPASSSVPPESVVFELSRQ   663
bipectinata   ETEGDPATTRIHSKETAR--------TGRNRGSKWMMDIICFRGQKVLRVPSA-SSVPPESVVFELSEQ   684
albomicans    -------ATRIQIKDA-----------PSSGHRNWMIG-LCFRGQKVLRVPPA-SSVPPESVVFELSEQ   702
grimshawi     ---------RIRSKES------------SAGRSNWMTN-LCFRGKKVLRVPPA-SSVPPESVVFELSEL   679
virilis       ------VGQRIRSTDDG---------TG--RNSNWMFG-LCFRGQKVLRVPPA-SSVPPESVVFELSEQ   623
hydei         ---------RIRGKEA--------AST-ARNNGNWKFS-LCFRGQKVLRVPPA-SSVPPESVVFELSEQ   668
novamexicana  ------VGQRIRSTDDG-------PDTG--RNSNWMFG-LCFRGQKVLRVPPA-SSVPPESVVFELSEQ   623
navajoa       -----RSGQRIRSKE-T-------ASTG-RSTGNWMFS-LCFHGQKVLRVPPA-SSVPPESVVFELSEQ   671
```

### Supplemental Figure 2

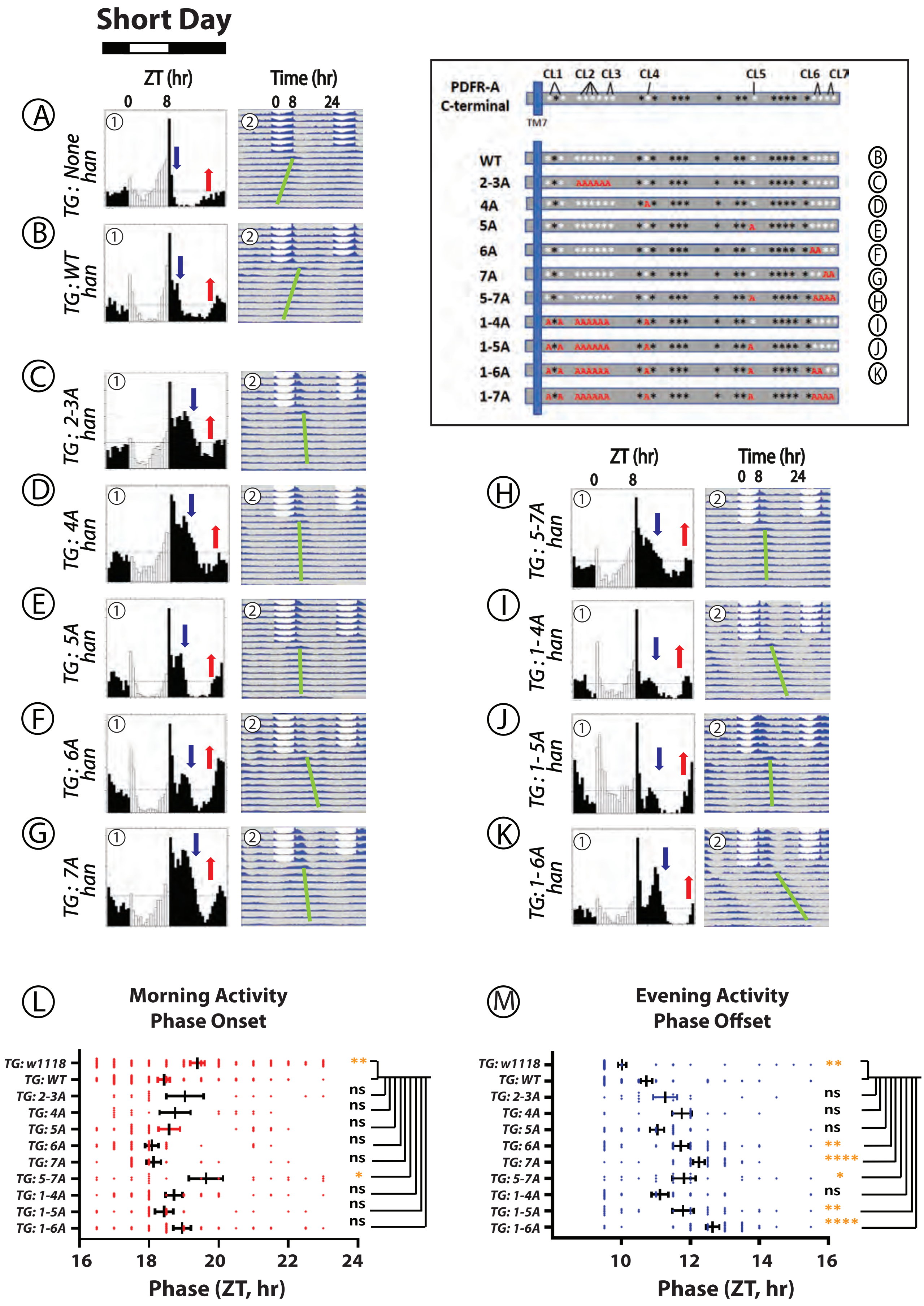

#### Supplemental Figure 3.

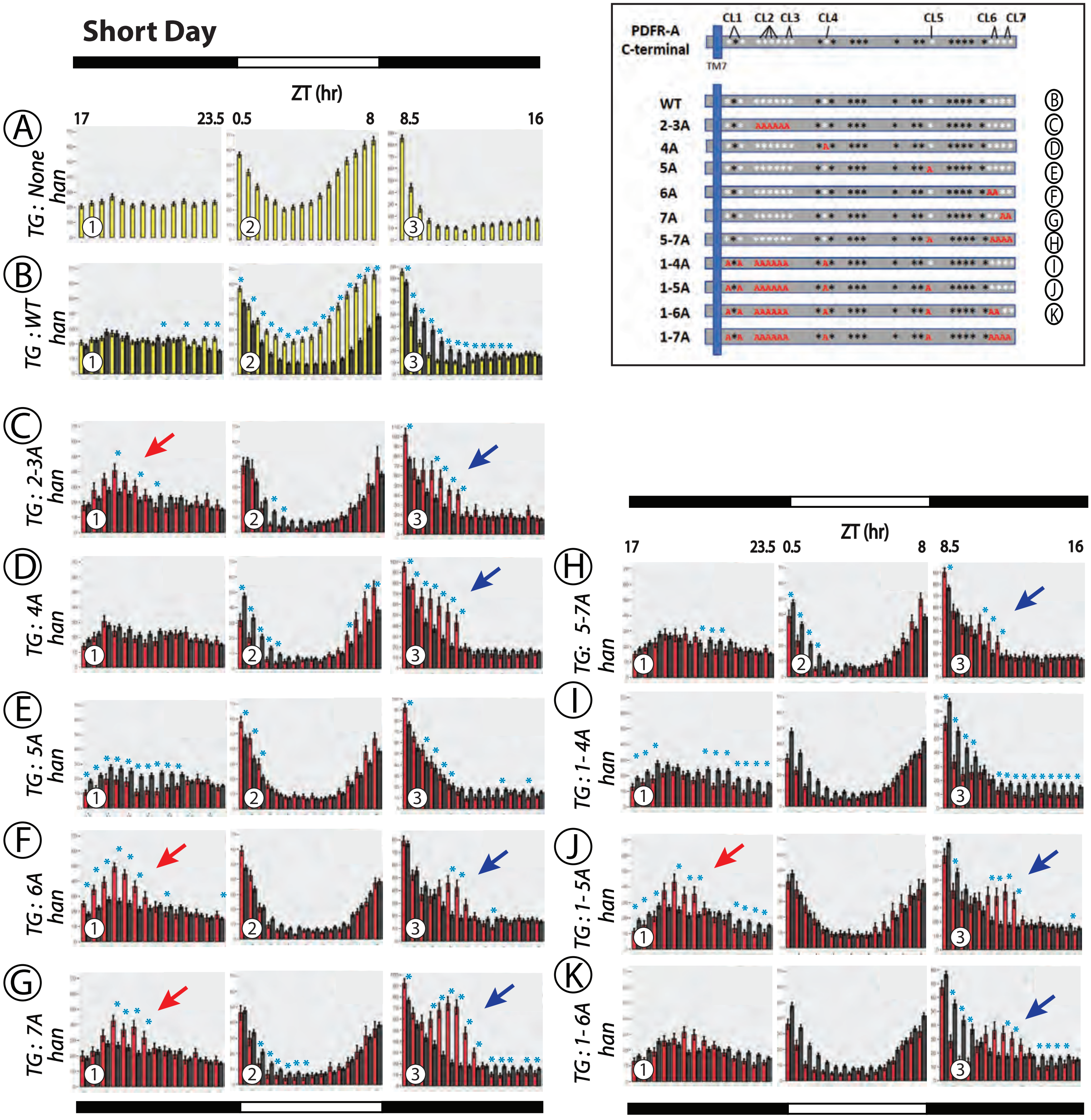

### Supplemental Figure 4

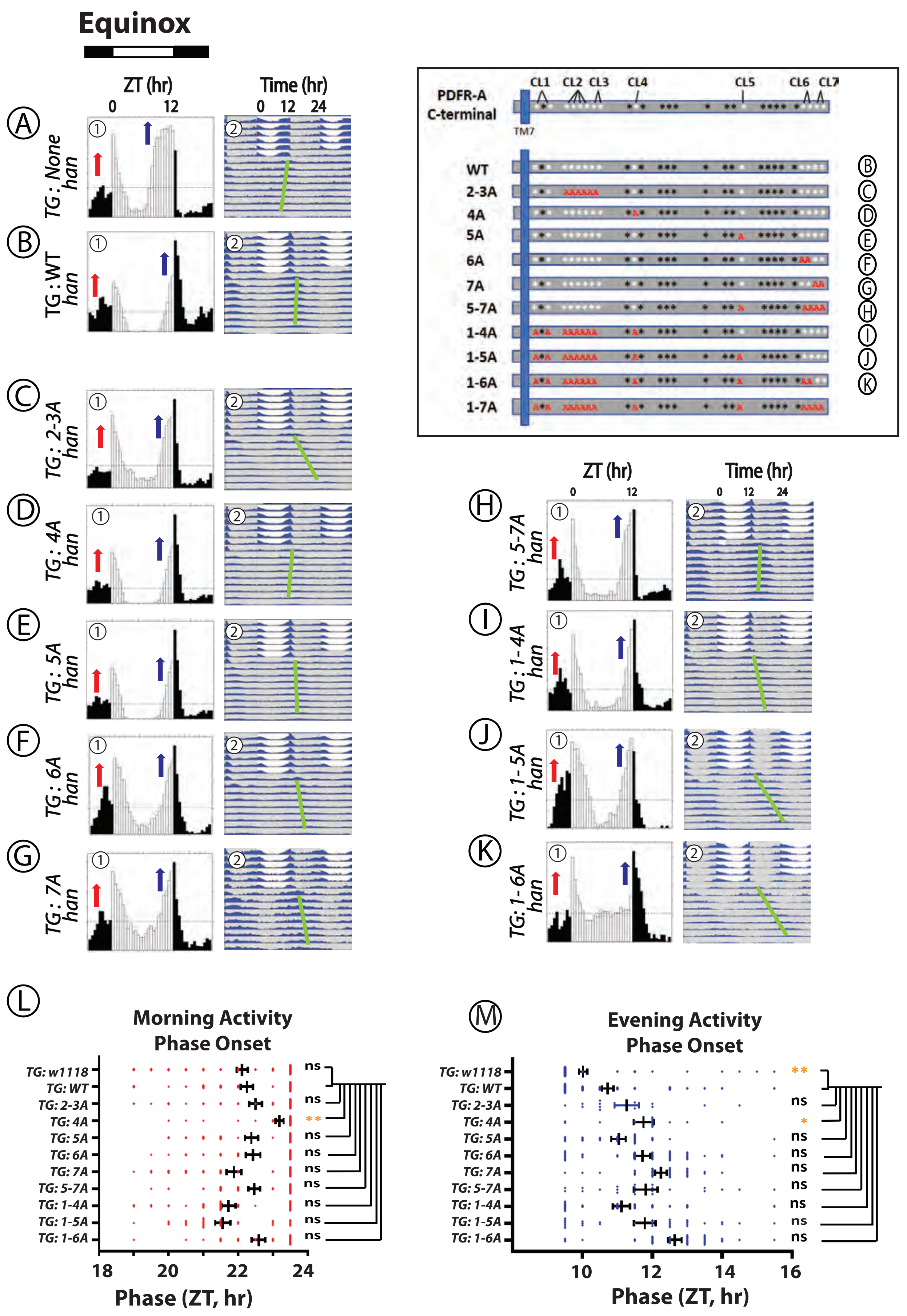

### Supplemental Figure 5.

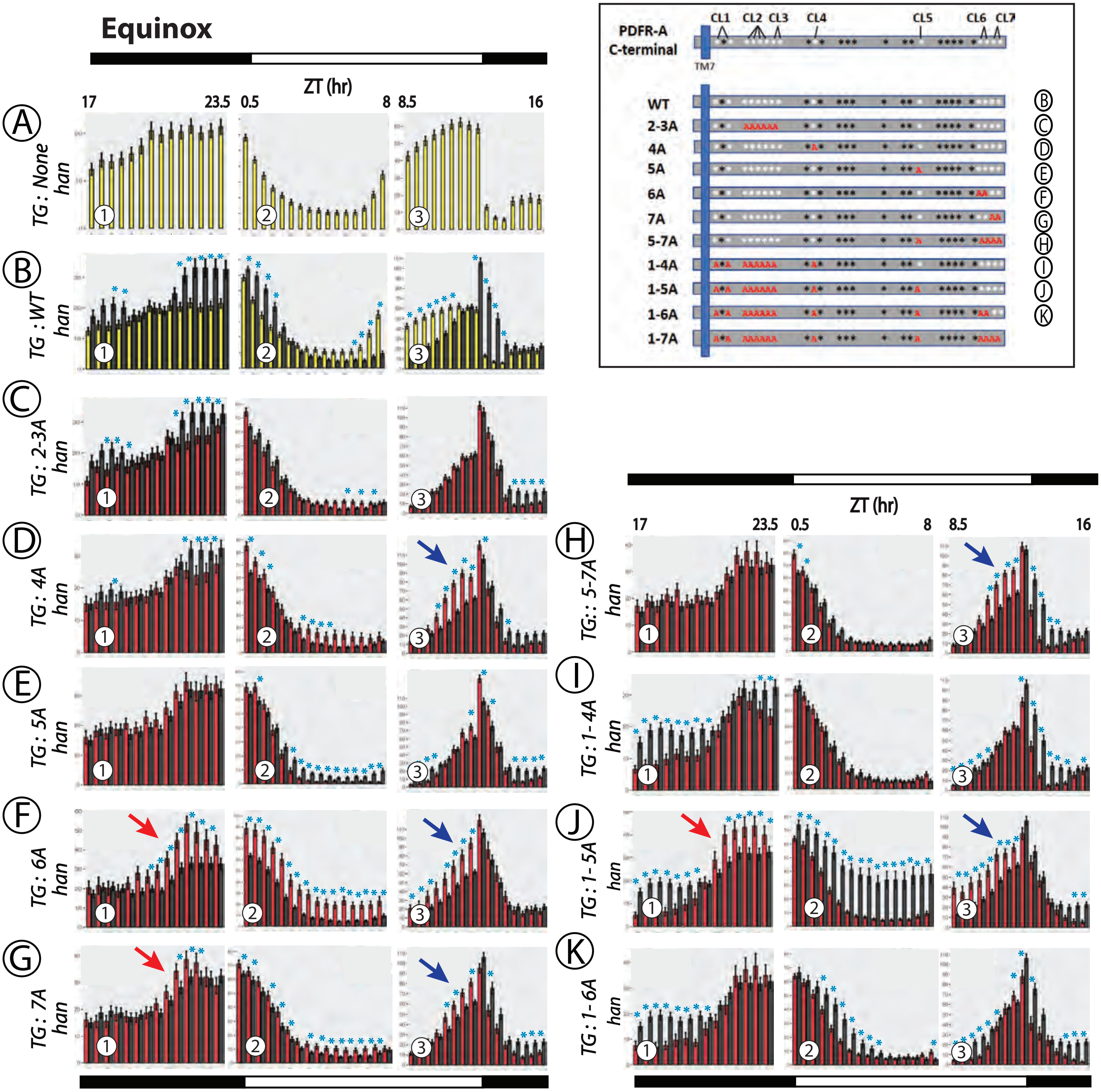

### Supplemental Figure 6

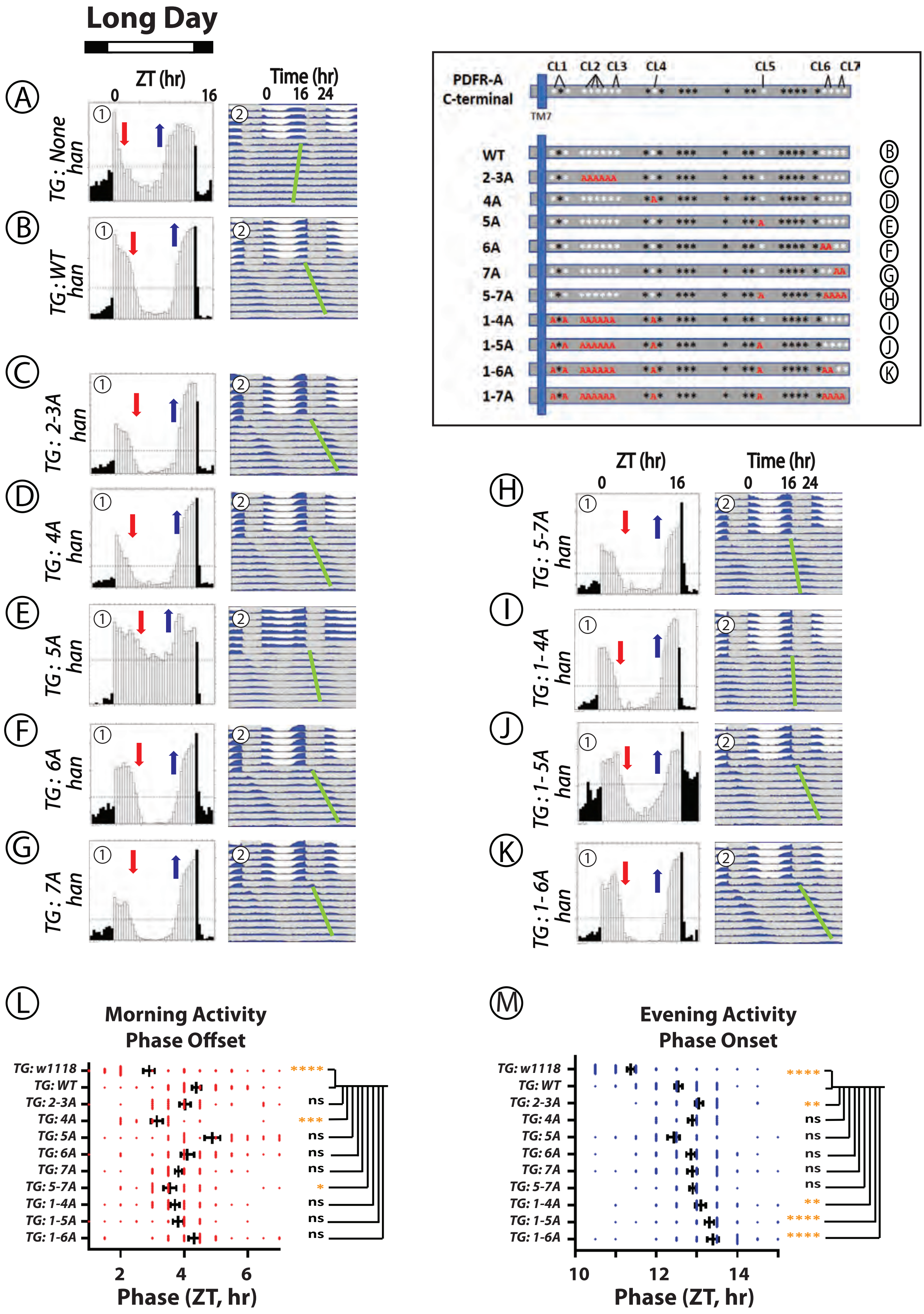

### Supplemental Figure 7.

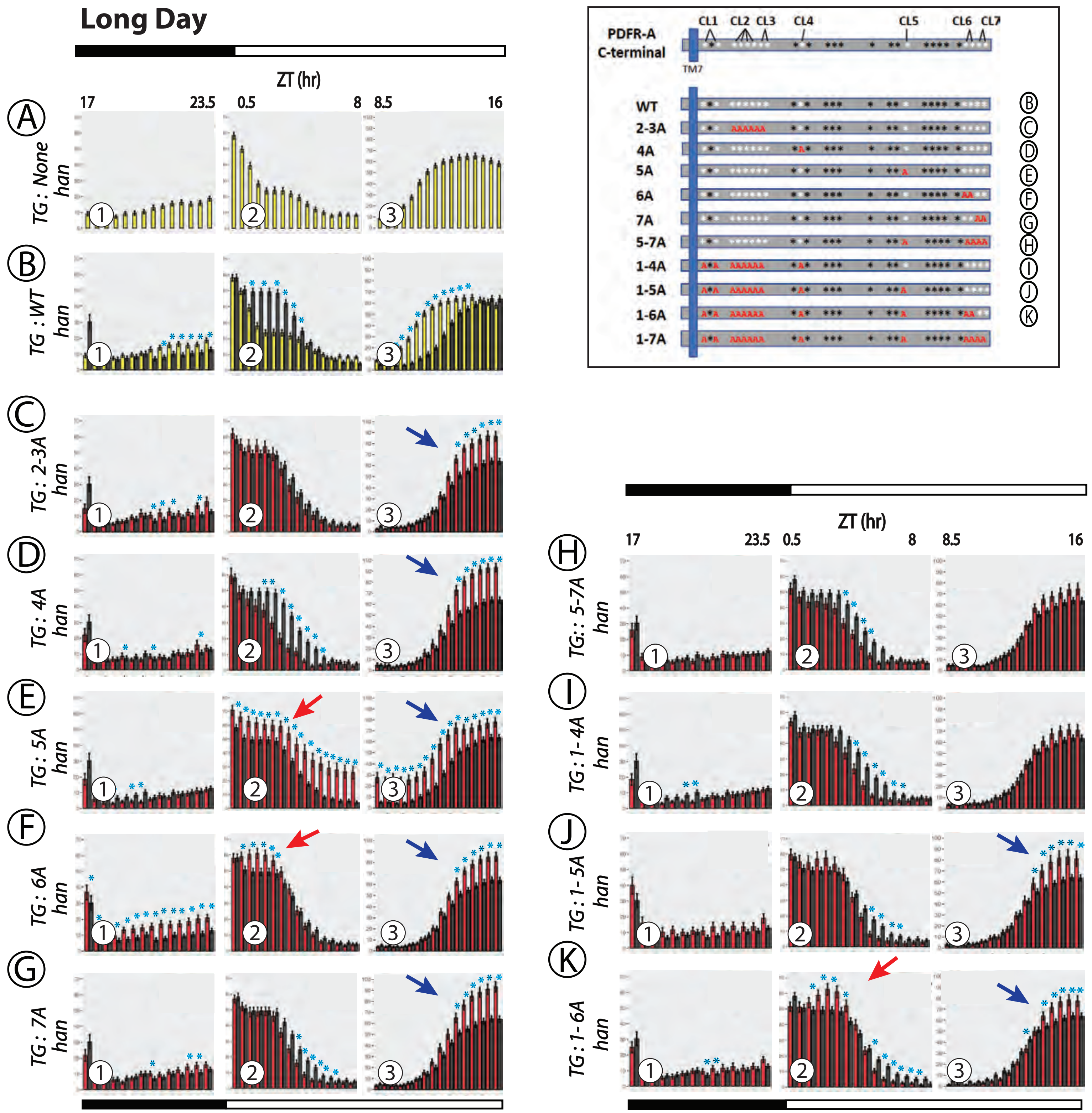

### Supplemental Figure 8.

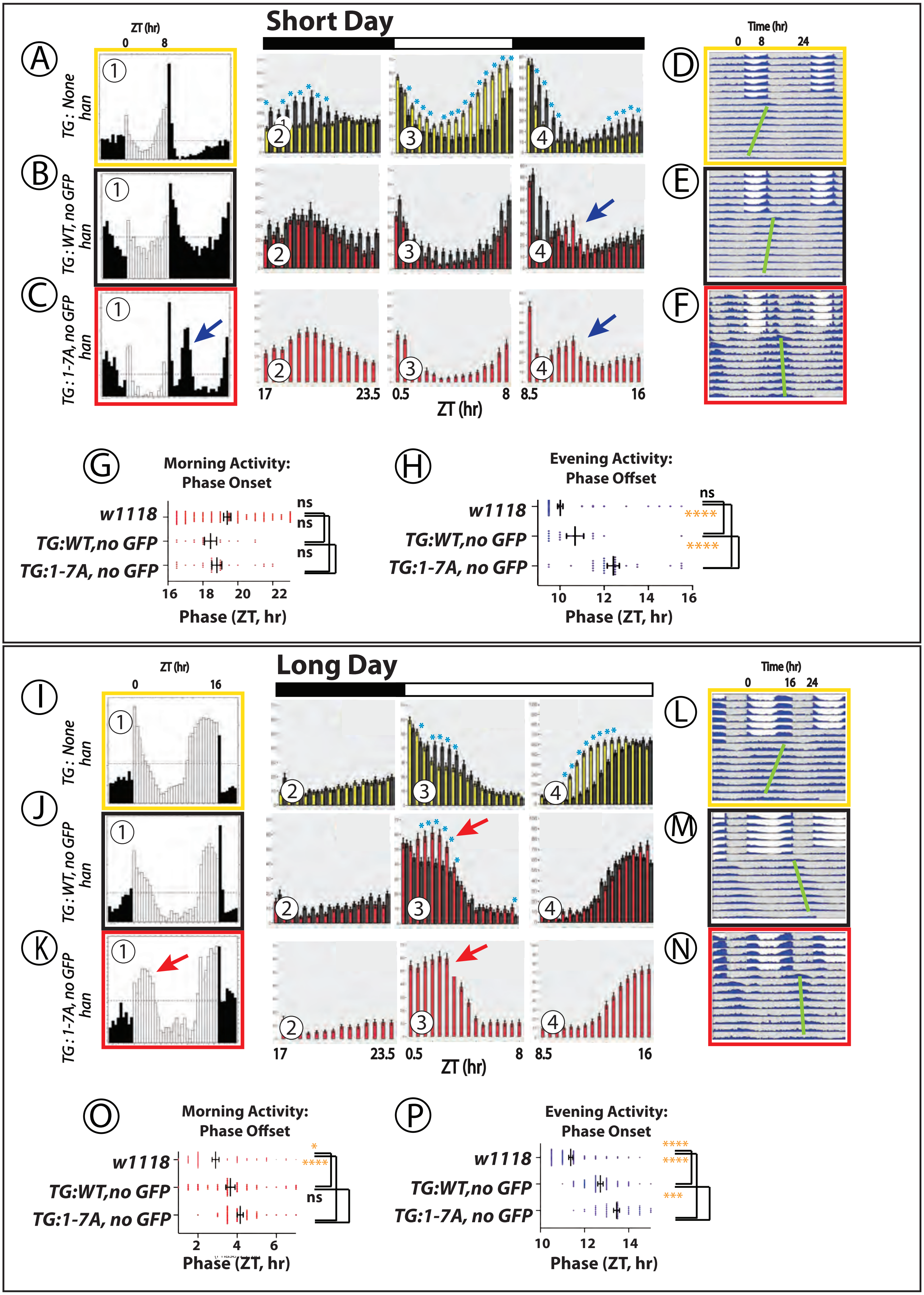

### Supplemental Figure 9

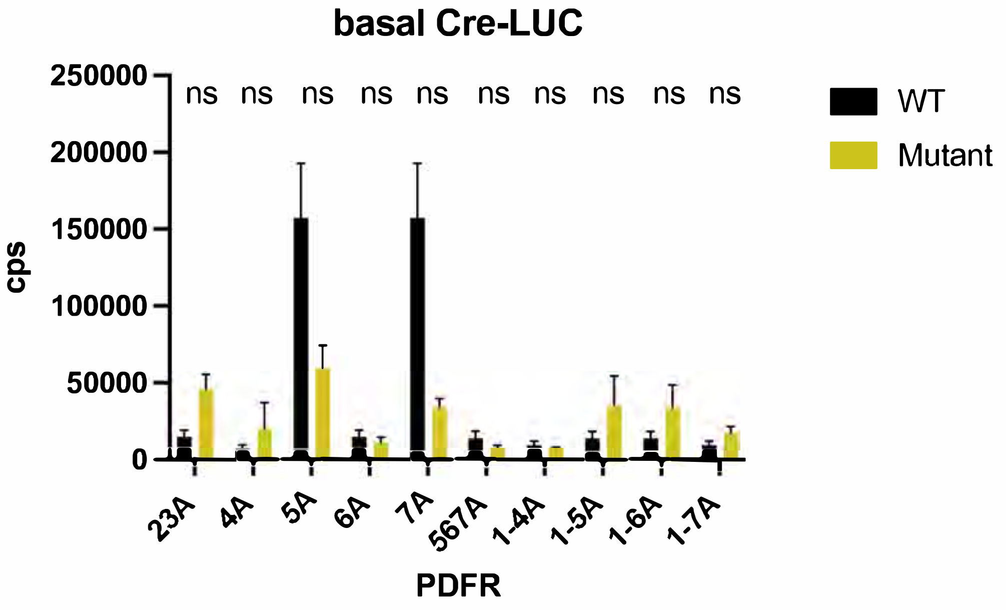

#### Supplemental Figure 10

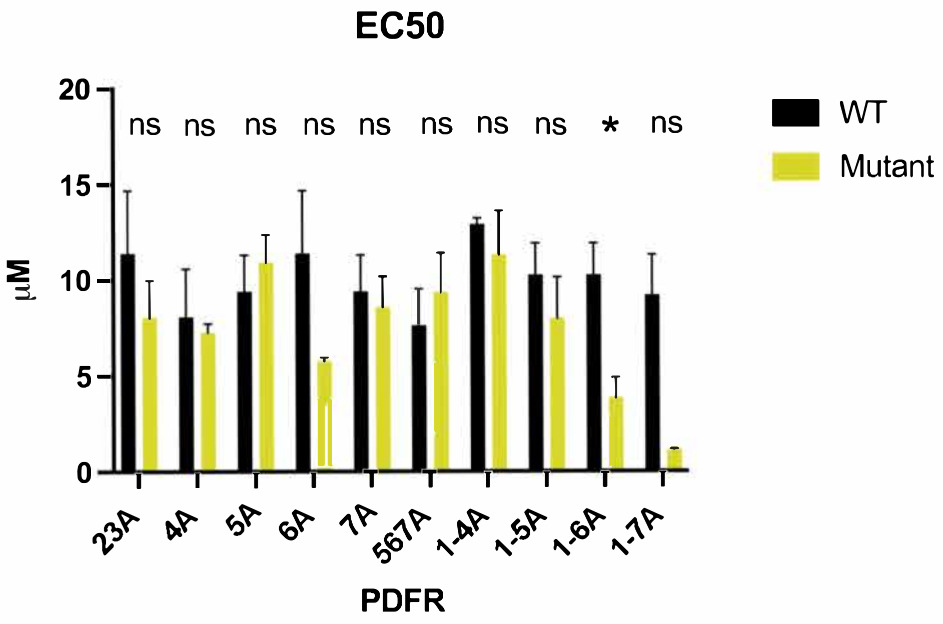

### Supplemental Figure 11

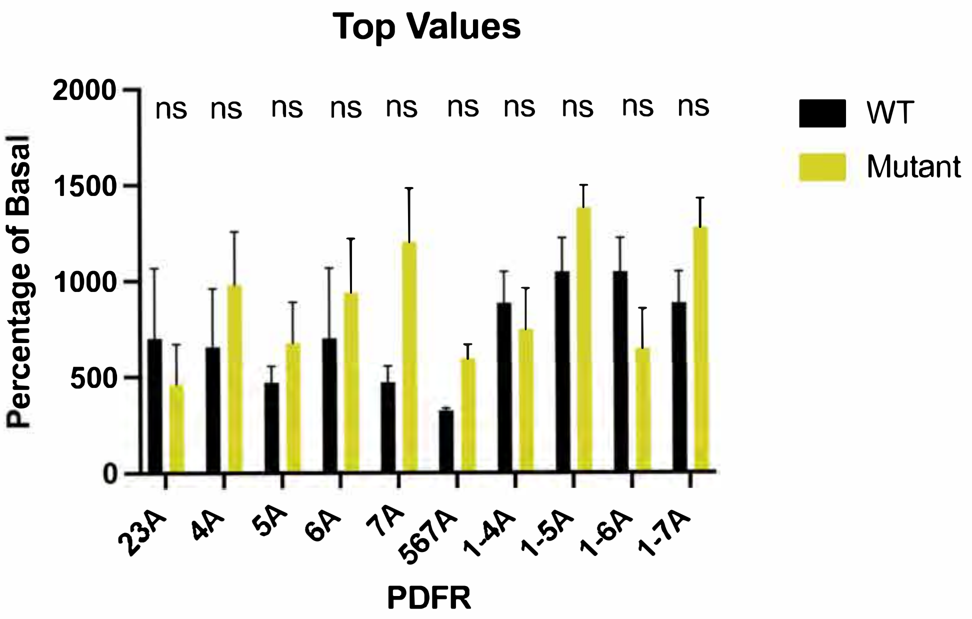

### Supplemental Figure 12

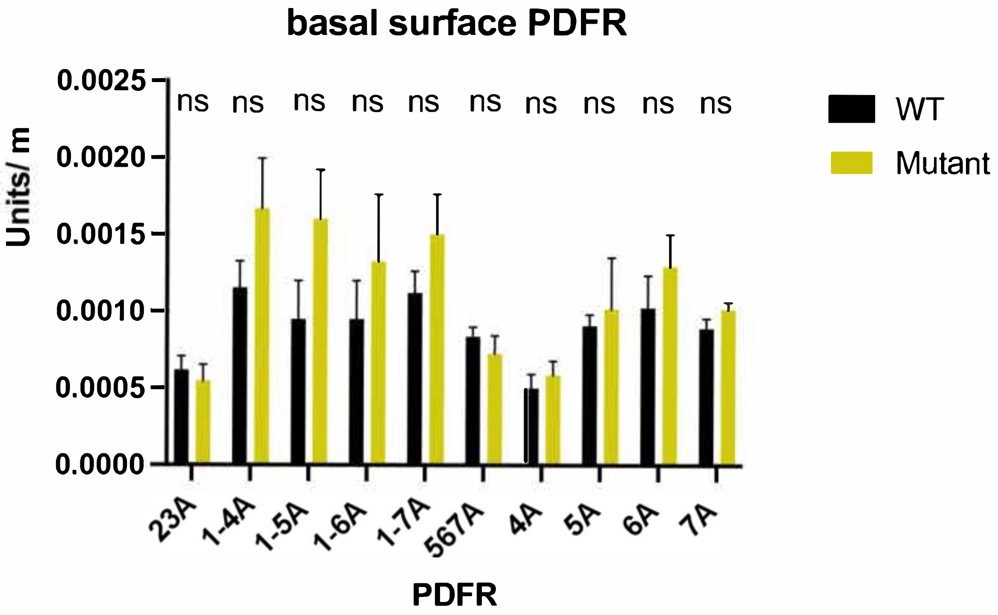

### Supplemental Figure 13

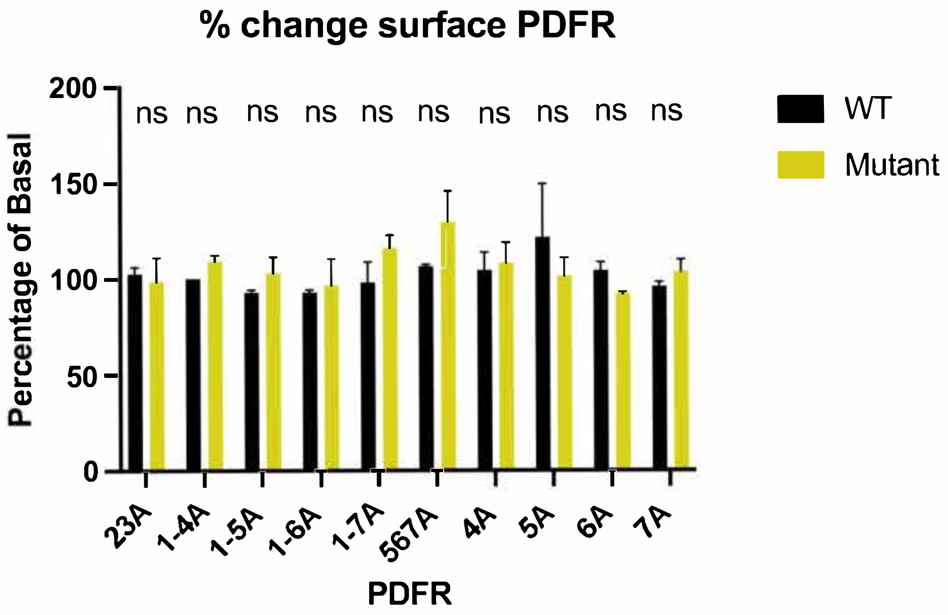

### Supplemental Figure 14

A. Exp 2
PDFR S531 CL2

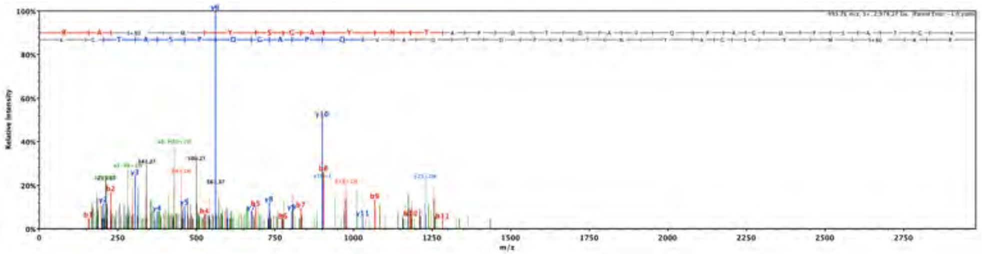

C. Exp. 3
PDFR T543 CL3

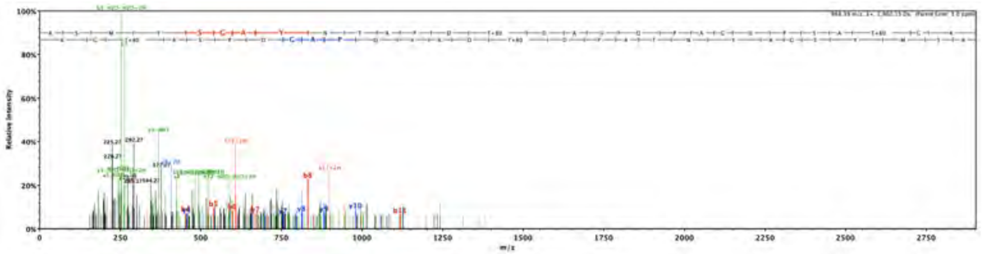

B. Exp. 3
PDFR S531 CL2

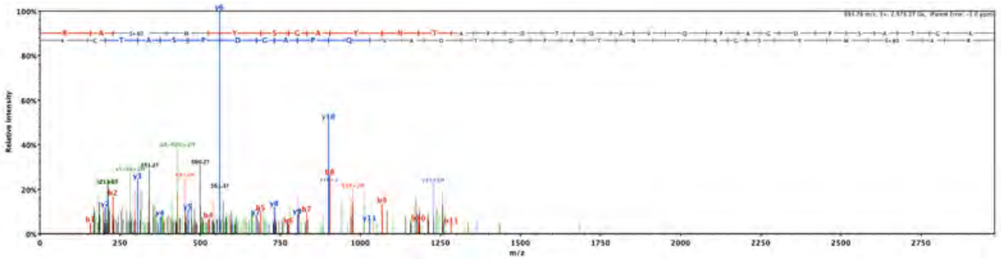

D. Exp 3
PDFR S560

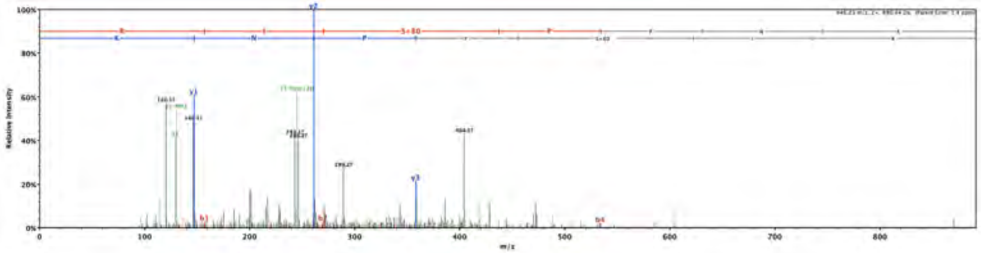

E. Exp. 5
PDFR – S531 CL2

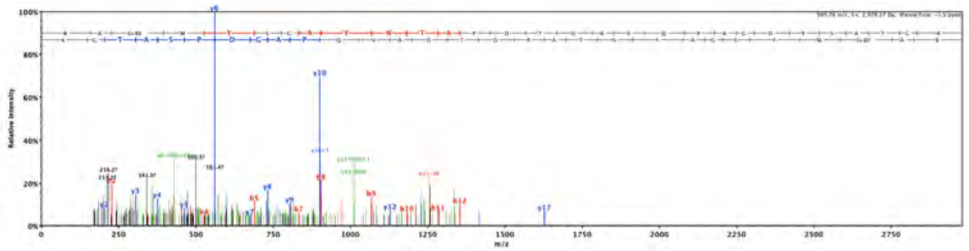

H. Exp. 7
PDFR S653

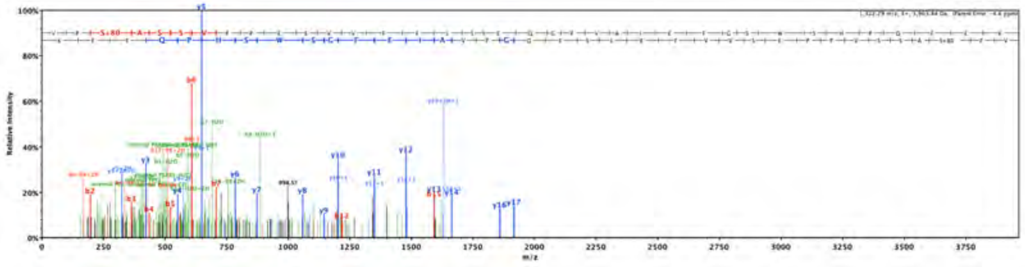

F. Exp. 5
PDFR - S534 CL2

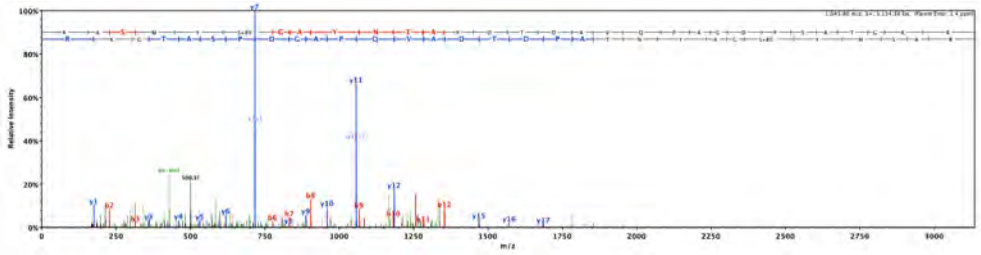

G. Exp 8
PDFR – S531 CL2

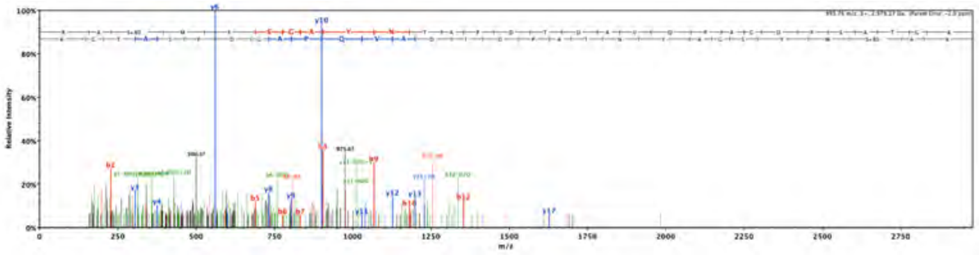

I. Exp. 8
PDFR – S560

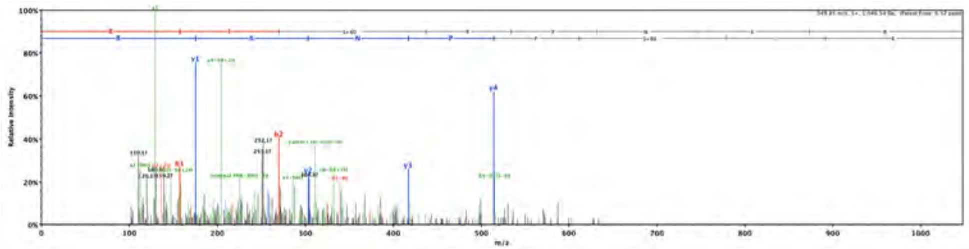

### Supplemental Figure 15

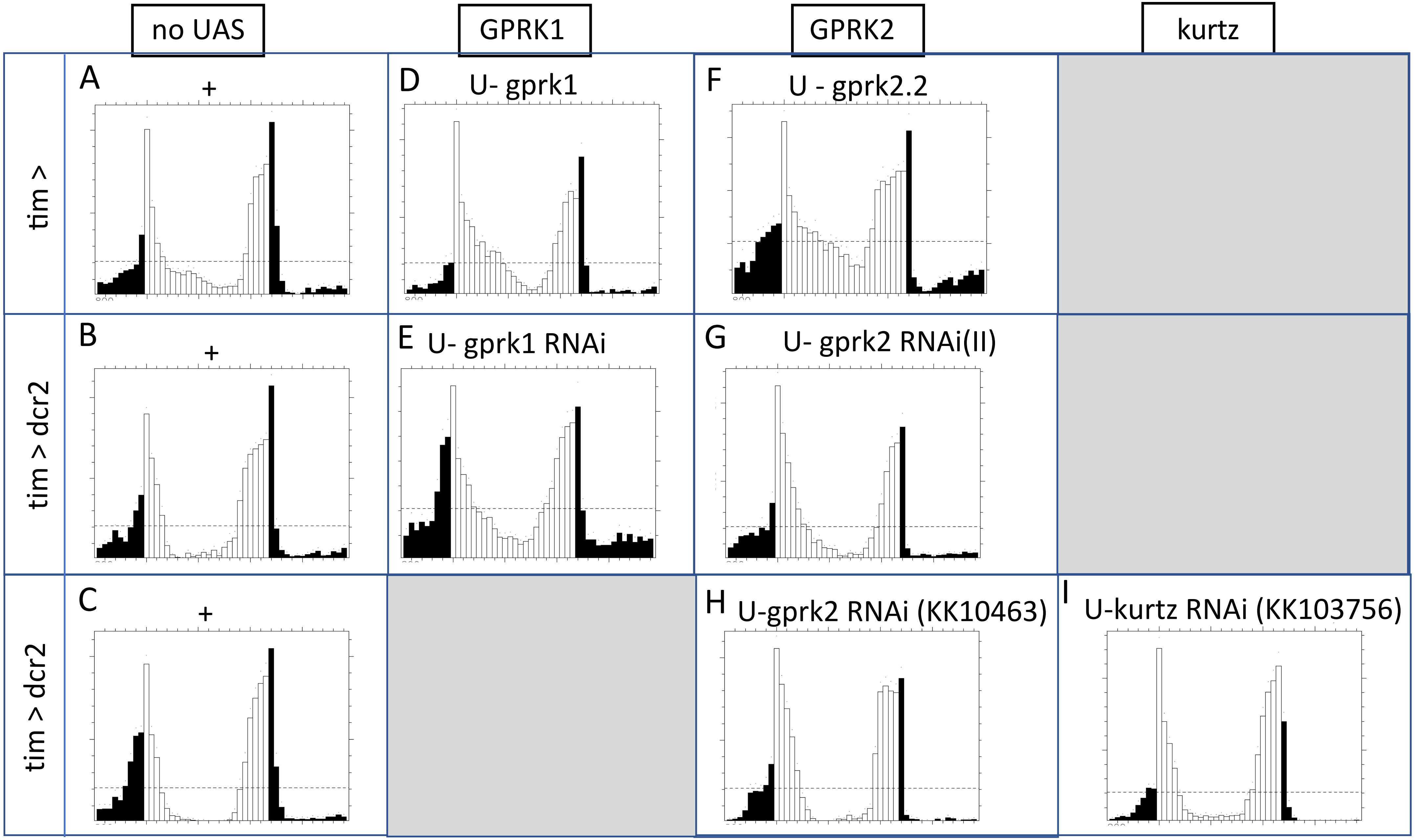
